# BoltzGen: Toward Universal Binder Design

**DOI:** 10.1101/2025.11.20.689494

**Authors:** Hannes Stark, Felix Faltings, MinGyu Choi, Yuxin Xie, Eunsu Hur, Timothy O’Donnell, Anton Bushuiev, Talip Uçar, Saro Passaro, Weian Mao, Mateo Reveiz, Roman Bushuiev, Tally Portnoi, Tomáš Pluskal, Josef Sivic, Karsten Kreis, Arash Vahdat, Shamayeeta Ray, Jonathan T. Goldstein, Andrew Savinov, Jacob A. Hambalek, Anshika Gupta, Diego A. Taquiri-Diaz, Yaotian Zhang, Samuel J. Snyder, A. Katherine Hatstat, Angelika Arada, Nam Hyeong Kim, Haoyu Fan, Ethel Tackie-Yarboi, Dylan Boselli, Lee Schnaider, Chang C. Liu, Gene-Wei Li, Denes Hnisz, David M. Sabatini, William F. DeGrado, Jeremy Wohlwend, Gabriele Corso, Regina Barzilay, Tommi Jaakkola

**Affiliations:** MIT; Boltz; Open Athena; CTU Prague; IOCB Prague; NVIDIA; IOCB Boston; UC Irvine; MPI; UCSF; HHMI; Jameel Clinic

## Abstract

We introduce *BoltzGen*, an all-atom generative model for designing proteins and peptides across all modalities to bind a wide range of biomolecular targets. BoltzGen builds strong structural reasoning capabilities about target-binder interactions into its generative design process. This is achieved by unifying design and structure prediction, resulting in a single model that also reaches state-of-the-art folding performance. BoltzGen’s generation process can be controlled with a flexible design specification language over covalent bonds, structure constraints, binding sites, and more. We experimentally validate these capabilities in eight diverse design campaigns with functional and affinity readouts across 26 targets. In our experiments, binder modalities span from nanobodies to disulfide-bonded peptides, and targets from disordered proteins to small molecules. In particular, we identify nanobody binders for novel targets with low similarity to proteins with already known bound structures. We release model weights, data, and both inference and training code at: https://github.com/HannesStark/boltzgen.

## 1 Introduction

De-novo binder design offers considerable potential for automating drug discovery. A number of previous techniques have been proposed to address parts of this challenge, including [Watson et al.,2023,Pacesa et al.,2024,Bennett et al.,2025,Mille-Fragoso et al.,2025]. However, several key limitations remain. For example, many of the techniques are tailored to specific classes of biomolecules such as nanobodies or peptides. As models learn to emulate physics primarily through examples provided, we believe expanding the generality of the method further improves its design capabilities for specific classes as well. Another key limitation has to do with evaluation as methods are often tested on targets that have closely related complexes in the training data. The potential of de-novo binder design comes precisely from its presumed ability to extrapolate beyond easy targets. We believe design methods should be tested accordingly. Moreover, in real-world discovery campaigns, a number of additional requirements and constraints govern successful designs. It is important to be able to control the design process in a flexible manner.

We introduce BoltzGen as an all-atom binder design method that generalizes across modalities and to novel targets. Extrapolation to unseen targets requires strong structural reasoning grounded in 3D geometry and physicochemical interactions. We hypothesized that models can acquire such reasoning capability by learning conserved patterns across binder and target modalities, such as hydrogen bonds forming the basis of DNA base-pairing, *α*-helix formation in proteins, and water bridges between small molecules and proteins. It is therefore advantageous to operate across modalities with a unified design framework. BoltzGen further reframes binder design as a pure structure prediction task and jointly trains on folding and design objectives. This yields a design model that matches state-of-the-art folding performance (Supplementary Figure 15) and provides the structure reasoning capacity required for high-quality binder design. Our generative process also offers an expressive interface (Figure 1b) for controlling the binder’s interaction with the target, such as specifying binding sites or covalent bonds. The method is described in detail in Section 3.

**Figure 1:**
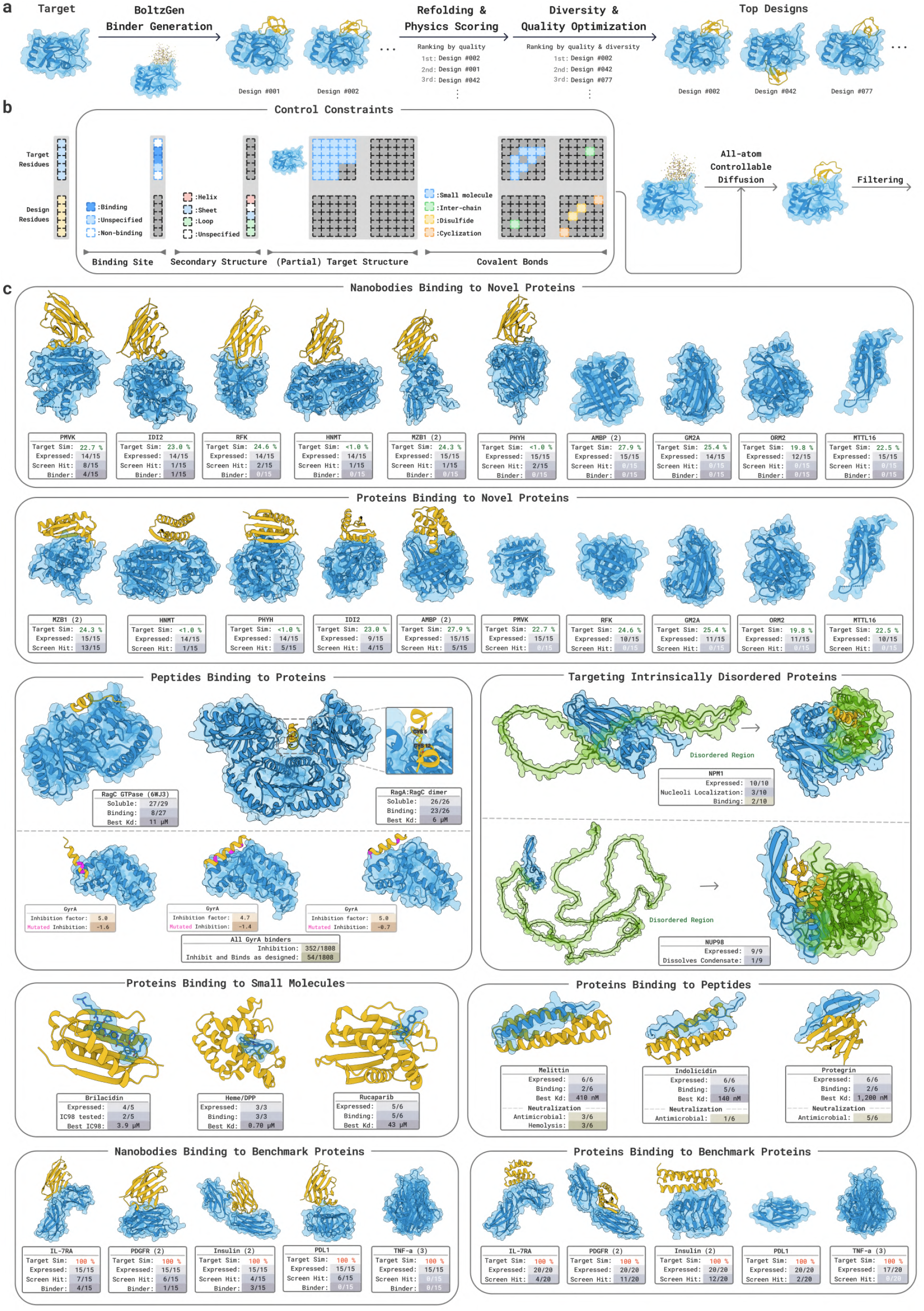
**a)** Schematic representation of the BoltzGen pipeline. A generative model samples tens of thousands of designs that are subsequently ranked and filtered according to computational metrics, resulting in a small set on the order of tens of candidates for experimental validation. **b)** Overview of constraints that can be specified in BoltzGen, including binding site, secondary structure, target structure, and covalent bonds. Target structure and bonds are specified as pairwise constraints between residues. **c)** Experimentally validated BoltzGen designs across 26 targets. In all structures, the designed binder is shown in yellow, and the target in blue. Brackets () indicate assay valency.

### Wetlab Validation

We experimentally validate our designs in a large-scale distributed effort involving multiple wetlabs. Each group selected targets and output modalities relevant to their specific application and then independently validated BoltzGen designs. To measure generalization capacity, we explicitly focus on targets that are dissimilar to any proteins for which bound structures exist.

Here we report the experimental results available to date; additional validation is ongoing. Some data are temporarily confidential at collaborators’ request, and we will update this work as further results become available.

1. Section 2.3: We design nanobodies against 10 novel targets for which there are no proteins with >30% sequence identity in a bound context in the entire PDB. Experimentally validating 15 or fewer designs against each target yields hits for 60% of them. The analogous experiment for protein binder designs results in hits against 50% of targets.
2. Section 2.4: When designing proteins to bind 3 bioactive peptides with diverse structures, we obtain nM binders for 2 and *µ*M binders for the other, while only testing 6 binders per target.
3. Section 2.5: We obtain protein binders and inhibitors against three small molecules.
4. Section 2.6: When designing linear peptides or cyclic disulfide-bonded peptides against two Ragulator proteins, we obtain low micromolar binders with hit rates of 88% and 28%.
5. Section 2.7: We generate and test 10 designs for binding the disordered region of NPM1. Two candidates show binding and co-localization with the target in live cells.
6. Section 2.8: Our campaign to design antimicrobial peptides binding to GyrA results in 19.5% of designs inhibiting cell growth by more than 4×.

### Open Source Release

We release training code, inference code, model weights, and all designs under the MIT License: https://github.com/HannesStark/boltzgen. The design pipeline is freely available, with easy-to-use interface to specify a binder-design problem and run BoltzGen, producing a filtered, ranked, diversity-optimized set of designs ready for experimental validation. We hope that fully open-sourcing the project puts state-of-the-art biomolecular design capabilities in the hands of any researcher and enables the community to build upon BoltzGen or contribute to BoltzGen-2.

## 2 Wetlab Results

This section explains the wetlab results with full methodology laid out in Appendix E. Unless mentioned otherwise, we provide the structure of the targets as input to BoltzGen.

### 2.1 Interpreting Affinities and Expression Numbers

#### Expression

To test a designed binder, one typically first produces the DNA that encodes the design. If the DNA is introduced into an environment with molecular machinery that translates DNA into proteins, the protein is produced, and the design is being *expressed*. Expression can fail for various reasons. For instance, a protein could fail to fold as intended, or a design could contain a large hydrophobic patch that binds to itself, causing aggregation (the proteins “clump up”). Usually, more stable and more soluble proteins have a higher chance of expressing well.

#### Affinity

Binding affinity describes how tightly two molecules stick to each other. It is often quantified via their dissociation constant *K*_*d*_, defined as the binder’s *off-rate* (how often they fall apart) divided by its *on-rate* (how often they come together). A smaller *K*_*d*_ indicates that the molecules stay bound longer and interact more strongly. Natural protein–protein interactions cover a broad range of affinities. In contrast, therapeutic binders are often optimized to achieve tighter binding. Antibodies and nanobodies frequently reach picomolar or low-nanomolar affinities. Representative affinities for selected therapeutic binders are summarized in Table 1. Importantly, a high affinity is only the first step toward an effective therapeutic. It indicates that a molecule can recognize and stably engage its target, but not whether it will reach the target in the body, remain stable, avoid off-target interactions, or produce the desired biological effect.

**Table 1:** Reported binding affinities (*K*_*d*_) of therapeutic antibodies and peptides.

| Drug | Target | $K_d$ (nM) |
| --- | --- | --- |
| Caplacizumab | vWF | 8.5 |
| Brolucizumab | VEGF-A | 0.03 |
| Ozoralizumab | TNFα | 0.02 |
| Degarelix | GnRH-R | 1.68 |
| Desmopressin | AVPR2 | 0.76 |
| Tirzepatide | GIPR | 0.13 |

### 2.2 Our Binder Classification

In an assay such as surface plasmon resonance (SPR) or biolayer interferometry (BLI), the designed protein is attached to a surface, and a solution containing the target is flowed across it, while recording a signal that tracks how the surface changes over time. This results in a set of curves called a sensorgram. In an ideal case, a design that does not bind gives a flat line, whereas for a binder, a clear rise and fall is present, and a set of equations can be fit to the curves to compute an affinity. In practice, other effects besides one-to-one design-target interaction can also produce a signal or noise in the sensorgram^1^.

As a consequence, a primary screen often yields sensorgrams that indicate an interaction but do not confidently confirm binding between the binder and the target. Much of the literature reports such cases as binders. We instead adopt the following categorization:

- **Screening hit:** The sensorgrams indicate an interaction. This could be a clean sensorgram with clear binding, but it also includes sensorgrams with an ambiguous signal. The latter case is reported as a hit from an initial screen, not as a confirmed binder.
- **Confirmed binder:** A design is called a binder only when a clean sensorgram, or similarly conclusive data, confirms binding, and an affinity is reported only when the kinetic data can support one. Such evidence may arise from a first screen, but ambiguous cases require follow-up assays. For instance, varying the capture level of the binder on the plate, adjusting the target’s concentration range, or inverting the assay orientation as to whether the target or the design is attached to the plate.

In this, we consider a sensogram as “clean” only if follows the guidelines:

1. Association times are long enough to resolve curvature clearly.
2. Both the association and dissociation curves can be fitted with an appropriate kinetic model for the valency of the target. For instance, the signal should decay by at least 5% during the dissociation phase to permit a reliable fit.
3. Little to no biphasic behavior is present in either phase if one-to-one binding is expected.
4. The on-rate and off-rate stand in a plausible relationship, avoiding the implausible pairing of very slow association (indicating weaker binding) with slow dissociation (indicating stronger binding).
5. The reference channel does not account for a substantial fraction of the unsubtracted binding-channel response.

### 2.3 Designing Nanobodies and Proteins against 10 Novel Targets

*Experiments carried out by Adaptyv Bio*.

Real-life design campaigns often focus on proteins for which no binders are known. To mirror this real-world scenario, we curated ten novel “hard” targets lacking bound structures in the Protein Data Bank (PDB). Specifically, we chose targets with less than 30% sequence identity to any protein observed in a bound state in the PDB (Supplementary Section E.1), resulting in sequence identities ranging between 1% and 27.9%. This is in contrast to prior works [Bennett et al., 2025, Chai-Discovery et al., 2025, Mille-Fragoso et al., 2025], where the target commonly appears in the training data in a bound context (Supplementary Section C.1). We additionally evaluate our method on five such “easy” targets from Chai-2 [Chai-Discovery et al., 2025], which we refer to as benchmark targets.

For each of the ten novel targets, we used BoltzGen to design 60,000 nanobodies and 60,000 proteins of lengths 80-140 (lengths 80-120 for the easy targets) without specifying a binding site. The 15 highest-ranking nanobodies and proteins per target were advanced to experimental validation. For each design, we conduct a primary binding assay with at least two replicas of surface plasmon resonance (SPR) and one biolayer interferometry (BLI) run (Methods Section E.1) in which the designs are immobilized. For nanobody binders, we further try to confirm the initial screening hits as one-to-one binders with orthogonal assays performed by both Sino Biological and Adaptyv Bio. On average, we obtain six replica binding assays for each design. All data is provided in Supplementary Section F.2.

For 6 of the ten “hard” novel targets, we obtain screening hits when testing nanobodies, several of which were confirmed as one-to-one binders in follow-up assays. Testing mini-protein designs yields screening hits for 5 of 10 novel targets (Figure 2a). Meanwhile, 3 of the 5 easy targets yield screening hits when testing miniproteins and 4 of 5 targets when testing nanobodies with confirmed one-to-one binders against 3 of them.

**Figure 2:**
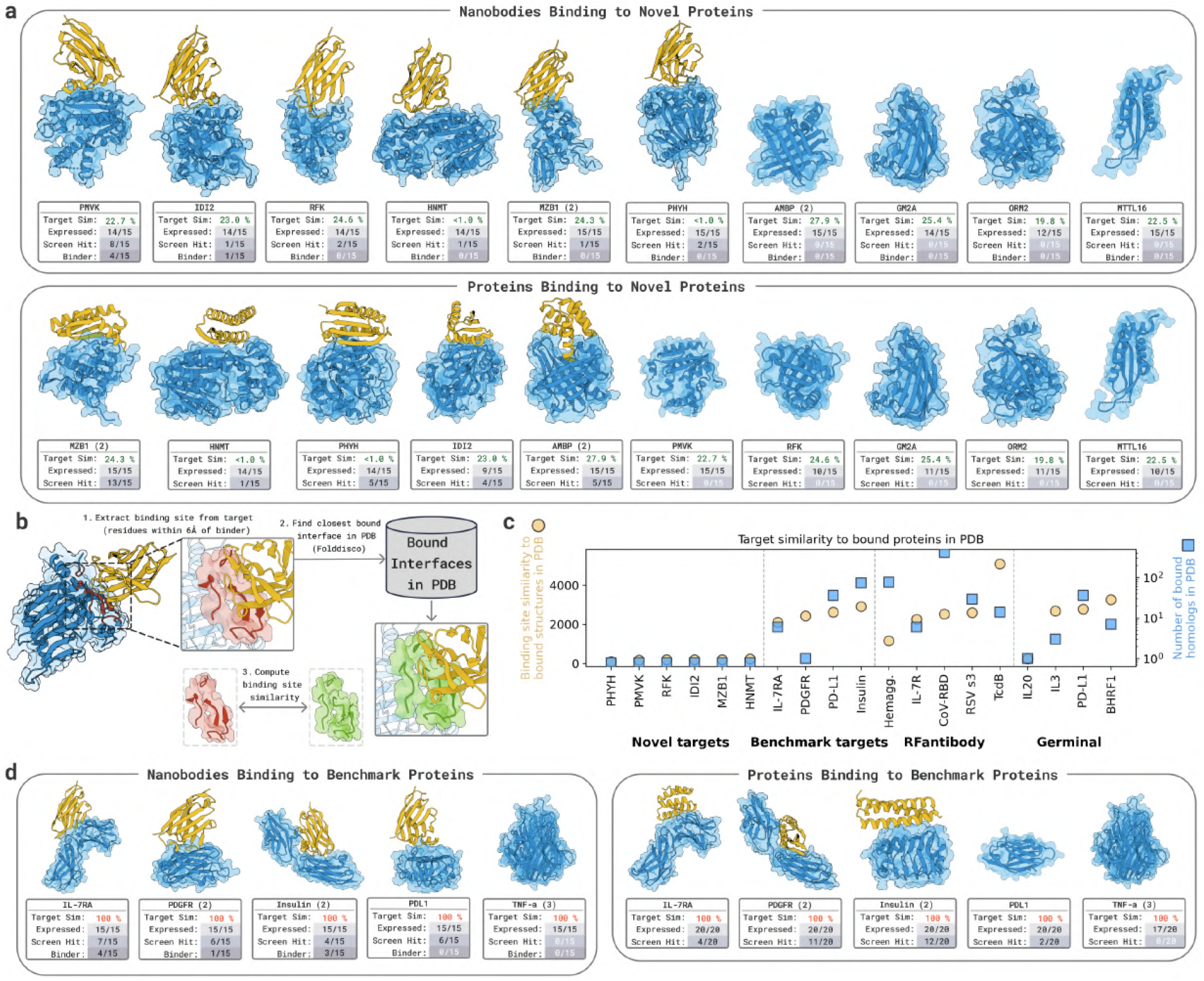
**a)** BoltzGen-designed nanobodies and proteins against ten novel targets. Target Sim. denotes the maximum sequence identity to any bound protein in PDB. Brackets () indicate assay valency. **b)** Binding site similarity computation procedure. We extract the target residues within 6Å of the binder and find the closest existing bound interface in PDB using Folddisco IDF scoring [Kim et al., 2025] to search through PPIRef [Bushuiev et al., 2024]. **c)** Target binding site similarity 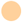 and number of times that the target already appears in a bound context in PDB 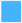, for targets in different nanobody design methods. **d)** BoltzGen-designed nanobodies and proteins against five benchmark targets considered in previous work. Each target appears with 100% sequence identity in at least one bound structure in PDB. Results are also summarized in Figure 1.

We next evaluated whether any of the binders exhibit promiscuous interactions. To this end, we performed identical SPR and BLI assays to test binding against human serum albumin (HSA), a highly abundant and interaction-prone plasma protein commonly used as a proxy for off-target binding. None of the nanobody designs showed detectable binding to HSA. To further probe selectivity, we profiled one mini-protein binder against IDI2 on the HuProt v4.0 Human Proteome Microarray, which spans *>*21,000 full-length human proteins (Figure 21). No off-target interactions were detected, confirming exclusive binding to IDI2 across the human proteome.

Investigating the nanobody binding sites reveals that the target interfaces are dissimilar from any bound target surfaces in all of PDB (Figure 2c). This contrasts with targets in prior nanobody design studies [Bennett et al., 2025, Mille-Fragoso et al., 2025], which were selected for therapeutic relevance rather than novelty and have appeared in a bound context in the PDB at least once, exhibiting highly similar bound interfaces (Figure 2c). This first-of-its-kind systematic evaluation of generalization in binder design shows that BoltzGen can generate high-affinity binders for novel targets without extensive screening, including nanobodies. .

### 2.4 Designing Proteins to Bind Bioactive Peptides with Diverse Structures

*Experiments by A. Katherine Hatstat, Angelika Arada, Nam Hyeong Kim, Ethel Tackie-Yarboi, Dylan Boselli, Lee Schnaider, and William F. DeGrado*.

We tested BoltzGen’s ability to generate binders of biologically active peptides with diverse secondary structures and antimicrobial and/or cytotoxic activity (Figure 3a,b). We targeted three peptides with diverse secondary structures: protegrin (disulfide-bonded beta-hairpin) [Gidalevitz et al., 2003, Steinberg et al., 1997], indolicidin (polyproline II or amphipathic conformation in the presence of bilayers) [Falla et al., 1996, Ladokhin et al., 1997, 1999], and melittin (amphipathic helix in the presence of membranes) [DeGrado et al., 1982, Dempsey, 1990, Guha et al., 2021, Hristova et al., 2001, Vogel and Jähnig, 1986].

**Figure 3:**
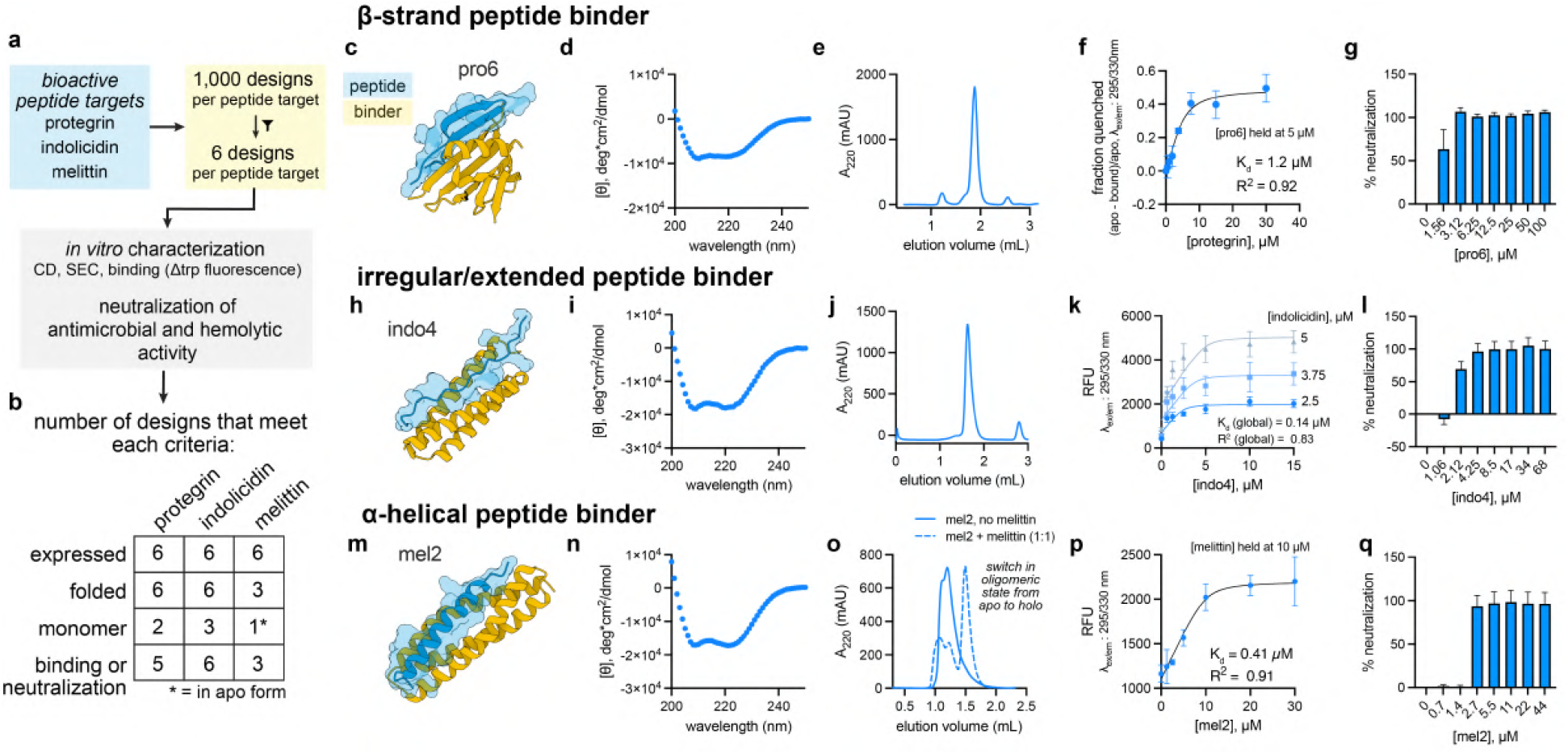
Experimental characterization of peptide binder designs. **A.** Design and characterization workflow for bioactive peptide targets with diverse topologies. **B**. Summary of experimental results for the six designs tested per peptide target. **C**. design model for pro6, the best design candidate for binding beta-strand peptide protegrin. **D–G**. Experimental characterization of pro6 by CD (d), SEC (e), in vitro binding assays (f) and antimicrobial neutralization assays against *B. subtilis* (g). **H**. design model for indo4, the best design candidate for the extended peptide target indolicidin. **M**. design model for mel1, the best design candidate for the peptide target melittin. Results are also summarized in Figure 1.

For each peptide target, we generated 1,000 ranked designs using BoltzGen. We then manually reviewed on the order of tens of candidates and selected six per target for experimental characterization, favoring designs with consistent burial, well-oriented hydrogen-bonding networks, and tightly packed interfaces. Because the binding targets are antimicrobial peptides (AMPs), we screened binders for their ability to neutralize antimicrobial activity against *B. subtilis* and, where applicable, their ability to inhibit peptide-induced hemolysis. The former also serves as a measure of the designs’ proteolytic stability as *B. subtilis* secretes numerous proteases. For each target, at least one binder design had single-digit *µ*M affinity (for protegrin) or nM affinity (for indolicidin and melittin) and neutralized antimicrobial and, for melittin, hemolytic activity (Figure 3b).

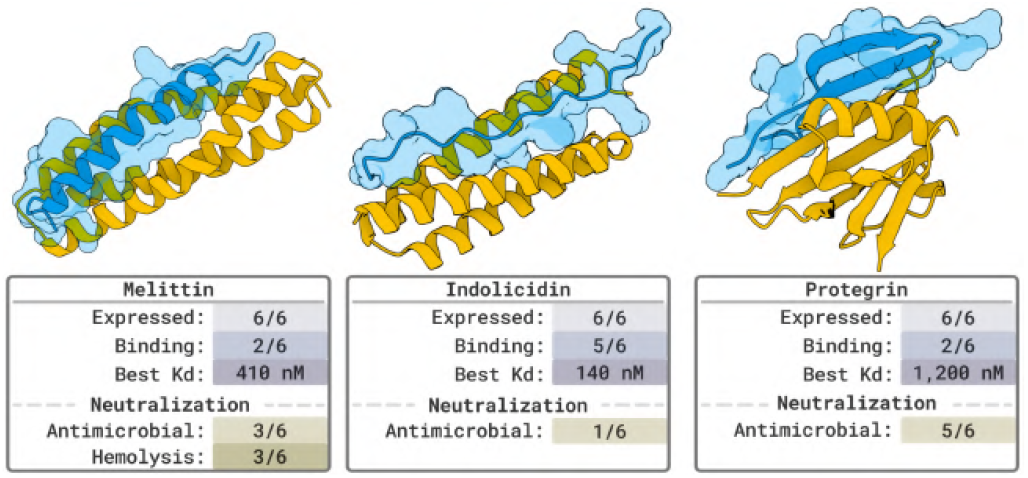

We first assessed the ability of BoltzGen to generate binders to protegrin, a disulfide-bonded, beta-strand antimicrobial peptide (Figure 3c-g; Supplementary Figure 22). All six designs expressed and showed the expected secondary structure by circular dichroism (beta or mixed alpha/beta) (Figure 3d; Supplementary Figure 22b). By Size exclusion chromatography (SEC), two designs formed single monodisperse species consistent with a monomeric state, while the remaining four eluted at volumes consistent with higher-order oligomers or mixed oligomeric states (Figure 3e; Supplementary Figure 22c). Two of the six designs, pro1 and pro6, showed detectable binding to protegrin via changes in binder tryptophan fluorescence, with affinities of 7.2 and 1.2 *µ*M, respectively (Figure 3f, Supplementary Figure 22d). Both pro1 and pro6 neutralized protegrin, with pro6 being a more potent neutralizer. This finding is consistent with the relative affinities of the two designs as measured by tryptophan fluorescence changes. While pro2, pro4, and pro5 showed little spectral shift in tryptophan fluorescence assays, they all neutralized protegrin’s activity against *B. subtilis* (Supplementary Figure 22e). The tryptophan residues in these designs are not predicted to be fully buried in the bound complex; thus, neutralization assays indicate a binding event is occurring, but it may not be detectable by the *in vitro* binding assay used herein (Supplementary Figure 22d,e).

Next, we tested six binder designs against indolicidin, an irregular/extended peptide (Figure 3h; Supplementary Figure 23). All six designs expressed and had helical character as measured by circular dichroism (Supplementary Figure 23b), and three of the designs (indo1, indo3, and indo4) formed single, homogenous species by analytical size exclusion chromatography (Supplementary indolicidin c). All indolicidin binder designs showed indolicdin binding as monitored by changes in indolicidin tryptophan fluorescence (Supplementary Figure 23d; *K*_*d*,indo4_ < *K*_*d*,indo1_ < *K*_*d*,indo5_ < *K*_*d*,indo2_ < *K*_*d*,indo3_ < *K*_*d*,indo6_), with indo 1, 3, and 5 exhibiting affinities < 5*µ*M and indo4 exhibiting sub-*µ*M affinity (Supplementary Figure 23d). The indo4:indolicidin interaction was further analyzed by SPR, which confirmed that the complex binds with nanomolar affinity (Supplementary Figure 23e). Despite all six designs having detectable indolicidin binding, only indo4 showed robust neutralization of indolicidin antimicrobial activity (Supplementary indolicidin f). This may be due to low proteolytic stability of the designs in the presence of *B. subtilis*, which secretes an array of proteases [van Wely et al., 2001].

Finally, we designed melittin binders and tested both their affinity and ability to neutralize melittin’s antimicrobial and hemolytic activity. All six selected melittin binder designs expressed, and mel1–3 displayed the expected helical structure (Supplementary Figure 24a,b). SEC showed that mel1 forms a monodisperse species consistent with a monomer, while mel2 and mel3 tended to aggregate (Supplementary Figure 24c). However, pre-incubation of mel2 and mel3 with melittin before SEC analysis shows that melittin binding acts as a switch, driving mel2 and mel3 from an aggregated state to monodisperse, monomeric species (Figure 3o; Supplementary Figure 24b). Mel4–6 form aggregates in SEC (Supplementary Figure 24b,c).

Mel1–3 were moved forward for assessment of melittin binding through *in vitro* binding assays and neutralization assays (both antimicrobial and hemolysis assays, as melittin has both antimicrobial activity and cytotoxicity) (Supplementary Figure 24d-g). All three binders neutralize antimicrobial activity and hemolysis (Figure 3 q; Supplementary Figure 24e-g). A single molar equivalent of Mel1 and Mel2 reversed the cytotoxic effect of melittin near its HD50 (1.2 *µ*M) for hemolysis of erythrocytes (Supplementary Figure 24f,g). Melittin contains a single tryptophan residue that is predicted to be near or fully buried in the peptide:binder interface in the designed complexes. Thus, we complemented neutralization assays by monitoring changes in intrinsic tryptophan fluorescence *in vitro* to quantify design binding affinity. Mel2 and mel3 have low to sub-*µ*M affinity for melittin (0.41 and 4.4 *µ*M *K*_*d*_ for mel2 and mel3, respectively), while mel1 did not show sufficient spectral shift at the excitation and emission wavelengths monitored to assess a *K*_*d*_ (Figure 3p; Supplementary Figure 24d). Together, these results demonstrate our pipeline can generate binders to bioactive peptides with diverse topologies. For all targets, we test only six designs and obtain single-digit *µ*M affinity or better, and show that the designed binders can inhibit antimicrobial and hemolytic activity of the target peptides.

### 2.5 Proteins Binding and Inhibiting Small Molecules

*Experiments by Nam Hyeong Kim, Haoyu Fan, and William F. DeGrado*.

To evaluate our pipeline’s ability to generate binders for small molecules and inhibitors of small-molecule activity, we tested six or fewer designs against each of brilacidin, heme, and rucaparib.

To assess the ability of BoltzGen to design proteins that bind to and neutralize the biological activity of larger small molecules, we focused on the antibiotic brilacidin, which is a complex arylamide with MW = 937 Da. We generated 9,999 designs and selected the five highest-scoring samples (bri1–bri5) for experimental validation. Four expressed well and were tested for neutralization of brilacidin’s antimicrobial activity following pre-incubation with the antibiotic. All four were active (Supplementary Figure 27), with IC_98%_ (the concentration required to achieve 98% inhibition of brilacidin’s activity). The proteins were further purified by size exclusion chromatography, and bri1 and bri5 showed a major peak eluting at a volume expected for the monomeric protein. After purification, 2.5 molar equivalents of bri1 fully neutralized the activity of 1.56 *µ*M brilacidin, indicating that its *K*_diss_ ≤ 3.9 *µ*M (Figure 4c). The IC_98%_ of bri5 was 31.2 *µ*M, indicating that its *K*_diss_ ≤ 31.2 *µ*M (Supplementary Figure 4b). The sequence identity of bri1 to any protein in PDB is 24.3%. These results demonstrate our pipeline’s ability to design novel and functional small-molecule inhibitors.

**Figure 4:**
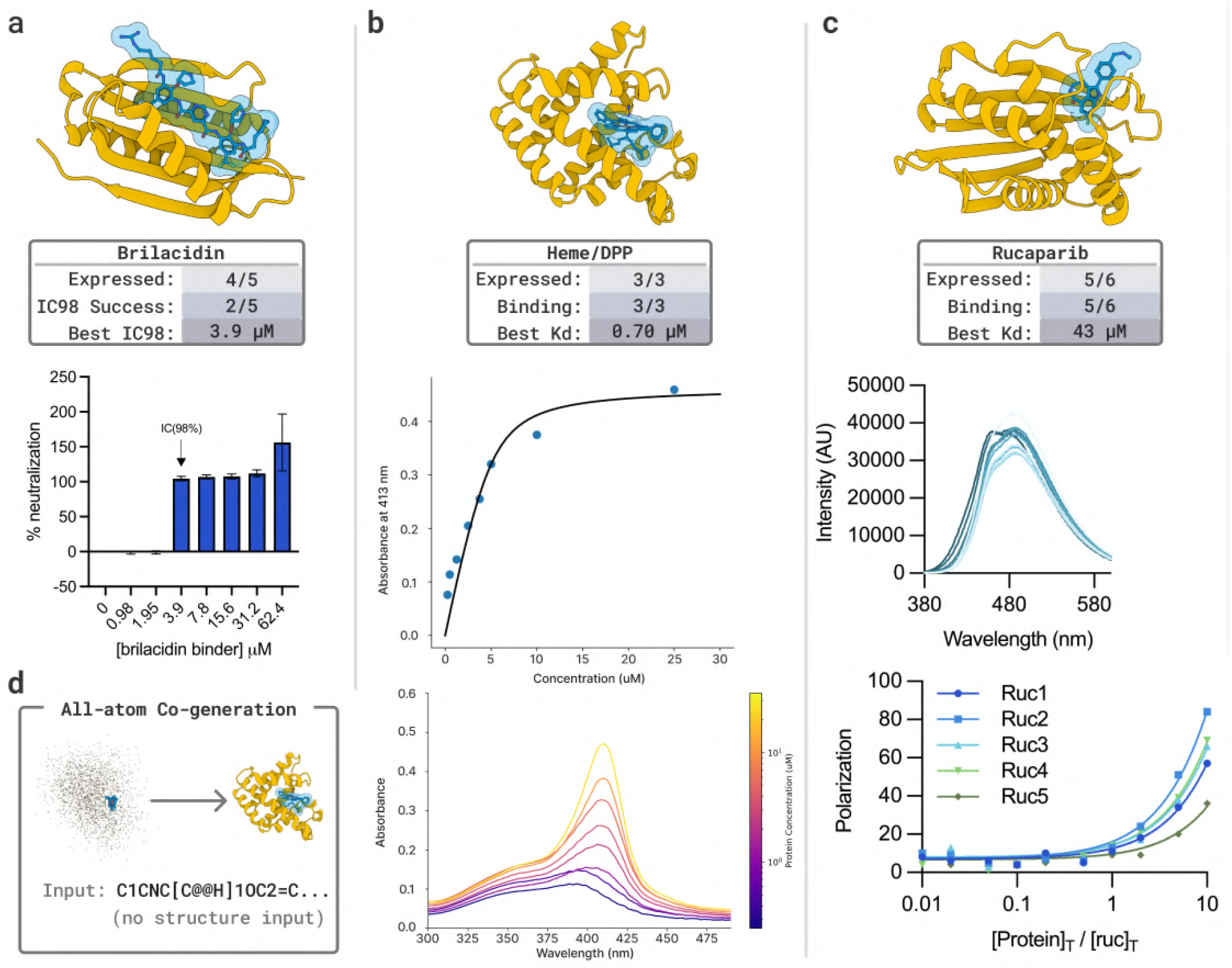
Structural models show the designed binder (yellow) and target (blue). **(a)** Brilacidin target: Neutralization assays (1.56 *µ*M) upon titration with increasing concentrations of designed protein (0–62.4 *µ*M). Neutralization was assessed by: % neutralization = ((*A*_obs_)*/*(*A*_control_)) ∗ 100 where *A*_obs_ and *A*_control_ are, respectively, the background-subtracted endpoint absorbance at 600 nm for the protein-treated sample versus the bacteria-only control (in the absence of both protein and brilacidin). At high concentration, the addition of protein increased the turbidity, causing an increase in absorption relative to control (which resulted in a %neutralization value slightly larger than 100%). **(b)** Heme target: Dissociation constants (*K*_*d*_) were determined from the absorbance at 413 nm by fitting to a quadratic ligand-binding model. UV–vis spectra (5 *µ*M) upon titration with increasing concentrations of the designed proteins (0–5 equivalents) **(c)** Rucaparib target: Fluorescence emission spectra (10 *µ*M) upon titration with increasing concentrations of the designed protein (0–10 equivalents). Fluorescence polarization assay with an excitation wavelength of 405 nm and emission wavelength of 516 nm. **(d)** All-atom bound structure of small molecules are generated without input conformation structures. Results are also summarized in Figure 1.

Next, we selected iron 5,15-diphenylporphyrin (FeDPP) as a benchmark target since de novo binders for both synthetic and natural porphyrins have been previously designed (RFDiffusion All-Atom [Krishna et al., 2024] tested 168 designs against heme and found 33 binders). We experimentally test three, which were the highest scoring of 20,000 BoltzGen designs ranked with its confidence and cavity size for substrate association. All three designs, named d1, d2, and d3, expressed in soluble form (20.4, 17.2, and 13.6 mg/L). However, DPP’s hydrophobicity and propensity to aggregate in aqueous buffer complicated reconstitution, likely because precipitation outpaced binding. Therefore, we turned to the structurally similar ferrous heme (protoporphyrin IX), which is more water-soluble. To assess heme binding, we conducted a titration, in which the visible spectrum of the heme (fixed concentration of 5 *µ*M) was monitored as a function of the concentration of protein. In aqueous buffer, the Soret band of heme is broad, with a peak near 398 nm; upon addition of protein, the peak sharpens, increases in intensity, and shifts to approximately 413 nm. This behavior is typical of mono-histidine ligatedm ferrous heme [Dolphin, 1978]. Dissociation constants (*K*_*d*_) were determined by non-linear least squares analysis of the titration data. The highest affinity was *K*_*d*_ 697±663 nM (Figure 4b), with values of *K*_*d*_ 2.40±1.31 *µ*M and 2.84±1.83 *µ*M for the remaining two designs (Supplementary Figure 26c). The sequence identity of d3 to any protein in PDB is 41.6%. These results indicate that BoltzGen can design nanomolar binders against small molecules when testing fewer than 10 designs.

Last, we chose rucaparib as a benchmark target for which prior work [Lu et al., 2024] achieved high-affinity binders (*K*_*d*_ < 5 nM) using an expert-guided approach specialized to rucaparib. We generated 10,000 designs using BoltzGen and advanced six candidates to experimental validation based on the model’s ranking, while additionally prioritizing designs that form hydrogen bonds with the carboxamide functional groups, as these interactions are considered essential for specific binding. Six proteins were selected for experimental validation by binding assay (Supplementary Figure 25a). Five designs expressed in moderate to good yields (15 – 69.5 mg/L). Incubation of each binder with equimolar concentrations of rucaparib led to a marked blue-shift and an increase in the intensity of its fluorescence spectrum, which is characteristic of the rucaparib indole core being bound in a rigid, solvent-inaccessible site (Supplementary Figure 25b). Fluorescence polarization data showed that ruc1–ruc4 exhibited moderate affinity for rucaparib, with values of 75.9, 43.0, 64.2, and 59.1 *µ*M *K*_*d*_, respectively (Figure 4). Ruc5 showed the weakest binding affinity (151.5 *µ*M *K*_*d*_). Affinities and binder details are summarized in Supplementary Table 12. The sequence identity of the highest affinity binder, ruc2, to any protein in PDB is 24% (with 19.4% of the hit covered by gaps).

### 2.6 Linear and Disulfide Bonded Peptides against Rag GTPase

*Experiments by Shamayeeta Ray, Jonathan T. Goldstein, and David M. Sabatini*.

For a general de novo design platform, the ability to rapidly nominate and synthesize peptides that selectively engage defined pockets on target proteins enables a range of applications. Beyond therapeutic development, such peptides can serve as structural and functional probes of binding-site biology, reagents for biochemical interaction assays, and starting scaffolds for downstream medicinal chemistry and small-molecule design.

The Rag GTPase heterodimer—comprising RagA or RagB bound to RagC or RagD—is a central hub in nutrient signaling that regulates protein synthesis, cell growth, and autophagy in part through control of mTORC1 activity. To evaluate peptide design performance on this system, we synthesized and tested 27 linear peptides with 4 scrambled-sequence controls, as well as 26 disulfide-bridged cyclic peptides designed with BoltzGen to target distinct protein–protein interaction surfaces on the RagA/RagC heterodimer. By surface plasmon resonance (SPR), 8/27 linear (Supplementary Figure 30) and 23/26 cyclic (Supplementary Figure 32) designs exhibited concentration-dependent binding. The affinities of the linear peptide binders span 11 *µ*M to 1073 *µ*M (Supplementary Figure 31), whereas those of the cyclic peptide binders range from 6 *µ*M to 629 *µ*M (Supplementary Figure 33) with the highest affinity binders showing in Figure 5.

**Figure 5:**
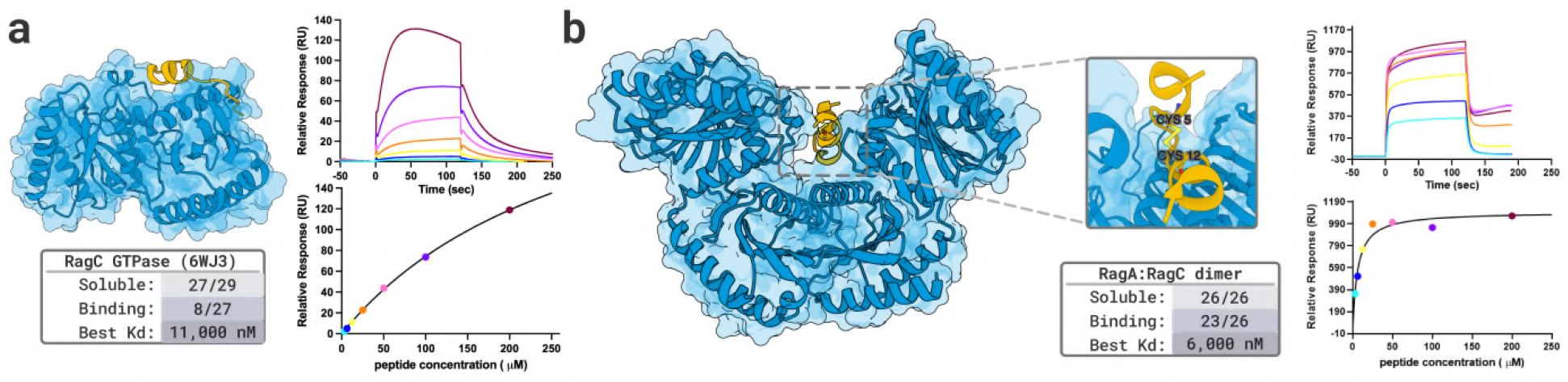
(a) Linear peptide 23 (yellow) bound to RagC GTPase (blue), corresponding subtracted SPR sensorgrams and steady-state affinity curve at concentrations of peptide ranging from 3.25-200 /micro M. (b) Cyclic disulfide-bonded peptide 26 (yellow) bound to RagA:RagC GTPase dimer (blue), corresponding SPR sensorgrams and steady-state affinity curves at concentrations ranging from 3.25-200 /micro M. Results are also summarized in Figure 1.

The SPR workflow clearly discriminated binders from non-binders based on sensorgrams. To further assess binding specificity for a subset of linear hits, we designed scrambled-sequence controls. Peptide 23 produced clean, concentration-dependent sensorgrams that enabled precise affinity determination (KD = 300 µM), whereas its scrambled control (scr23) showed minimal binding at low concentrations and yielded poorly behaved, uninterpretable sensorgrams above 25 µM—consistent with nonspecific interactions at higher analyte concentrations. Overall, these results demonstrate that Boltz can generate short linear and cyclic peptides (6–20 amino acids) that bind Rags, with an experimental confirmation rate of ∼30% for linear designs, and ∼88% for cyclic peptides with dissociation constants in the single-digit micromolar range for best-performing designs. Importantly, prior efforts in our group to identify Rag GTPase binders have been unsuccessful, suggesting that BoltzGen’s de novo peptide design may provide a productive new path toward Rag GTPase chemical biology tools and, ultimately, Rag GTPase modulators.

### 2.7 Peptides Targeting Disordered Proteins

*Experiments by Yaotian Zhang, Samuel J. Snyder, and Denes Hnisz*.

#### NPM1 target

We first considered NPM1, a nucleolar protein in the micron-sized nucleolus condensate of human cell nuclei [Lafontaine et al., 2021]. Ten selected NPM1 IDR binders were fused to msfGFP and transiently expressed in U2OS cells, and their subcellular localization was assessed by GFP fluorescence (Figure 6A). Three binders (NPM1-binder 8, 14, 19) showed nucleolar enrichment (Supplementary Figure 34B), which was confirmed by immunofluorescence colocalization with endogenous NPM1 (Figure 6B and Supplementary Figure 34C). SURF6 staining served as an additional nucleolar marker (Figure 6B). Biolayer interferometry for the three successful candidates confirmed binding for two of them.

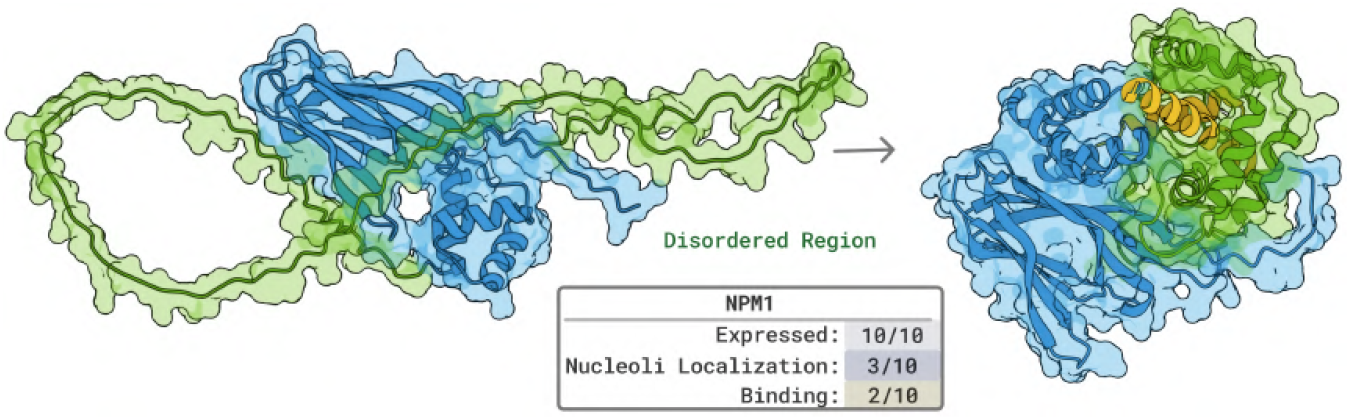

**Figure 6:**
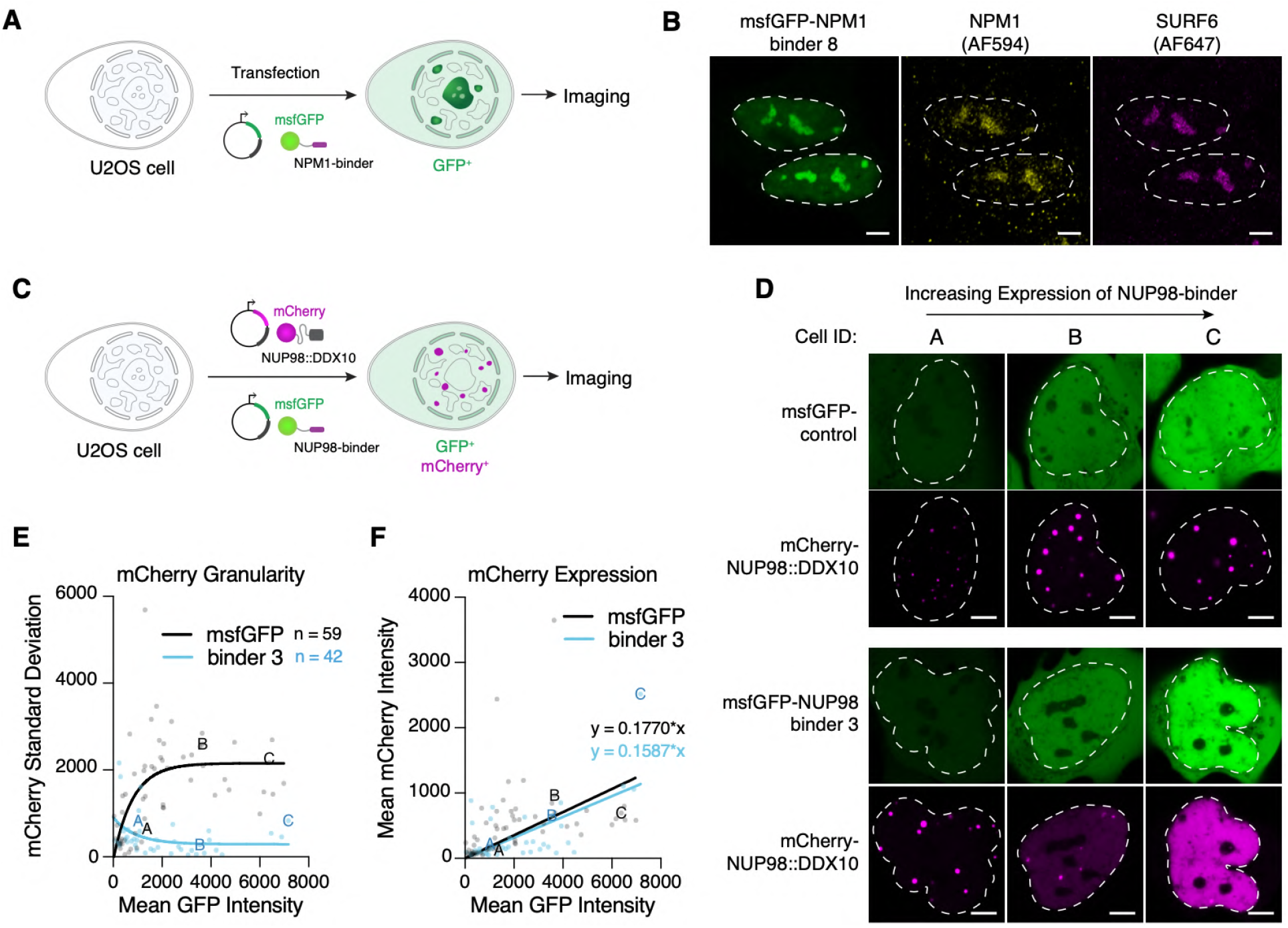
**A.** Schematic model of the msfGFP-tagged NPM1-binder design and cellular assay to visualize subcellular localization of the NPM1-binders. **B**. Representative fixed-cell immunofluorescence of U2OS cells expressing exogenous msfGFP-NPM1-binder 8. Endogenous NPM1 and SURF6 are stained with antibodies. The cell nucleus is highlighted with a dashed white line contour. Scale bar: 5 µm. The experiment was repeated twice independently with similar results. **C**. Schematic of the msfGFP-tagged NUP98-binder constructs and the mCherry-NUP98::DDX10 reporter (containing the NUP98 IDR), and the live-cell co-expression assay used to visualize their subcellular localization. **D**. Representative single-cell examples showing increasing msfGFP-control and msfGFP-NUP98-binder 3 expression (left to right) with corresponding mCherry-NUP98::DDX10 signal distribution. Cell IDs match the labeled data points in E and F. The experiment was repeated twice independently with similar results. **E**. Nuclear mCherry signal standard deviation (granularity) plotted against mean nuclear GFP intensity for msfGFP control and NUP98-binder 3. Labeled points correspond to the cells shown in D. n = 59 (msfGFP control) and n = 42 (msfGFP-NUP98-binder 3) cells from two biologically independent experiments. Curves show one-phase decay fits. **F**. Mean nuclear mCherry intensity plotted against mean nuclear GFP intensity for the cells quantified in E. Labeled points correspond to the cells shown in D. Results are also summarized in Figure 1.

#### NUP98 target

Encouraged by these results, we turned to NUP98, a nucleoporin whose IDR is frequently translocated in myeloid leukemias. NUP98 fusion proteins form micron-sized nuclear condensates in an IDR-dependent manner [Terlecki-Zaniewicz et al., 2021], motivating us to design binders capable of dissolving these condensates. Nine selected NUP98 IDR binders were fused to msfGFP and transiently co-expressed with mCherry-tagged NUP98::DDX10 in U2OS cells, and subcellular localization was assessed by GFP and mCherry fluorescence, respectively (Figure 6C). Binder 3 appeared to dissolve NUP98::DDX10 condensates: at high binder 3 expression, the mCherry–NUP98::DDX10 signal became diffuse, and the extent of dissolution—quantified as the standard deviation of nuclear mCherry intensity—increased with binder 3 abundance (Supplementary Figure 6D–E). Differences in binder expression levels did not account for the observed redistribution of mCherry–NUP98::DDX10 (Figure 6F). Together, these results support target engagement by binders designed against the NPM1 and NUP98 IDRs in live cells. For NPM1, engagement was reflected by nucleolar enrichment and corroborated by colocalization with endogenous NPM1. For NUP98, engagement was reflected by altered nuclear distribution of NUP98::DDX10.

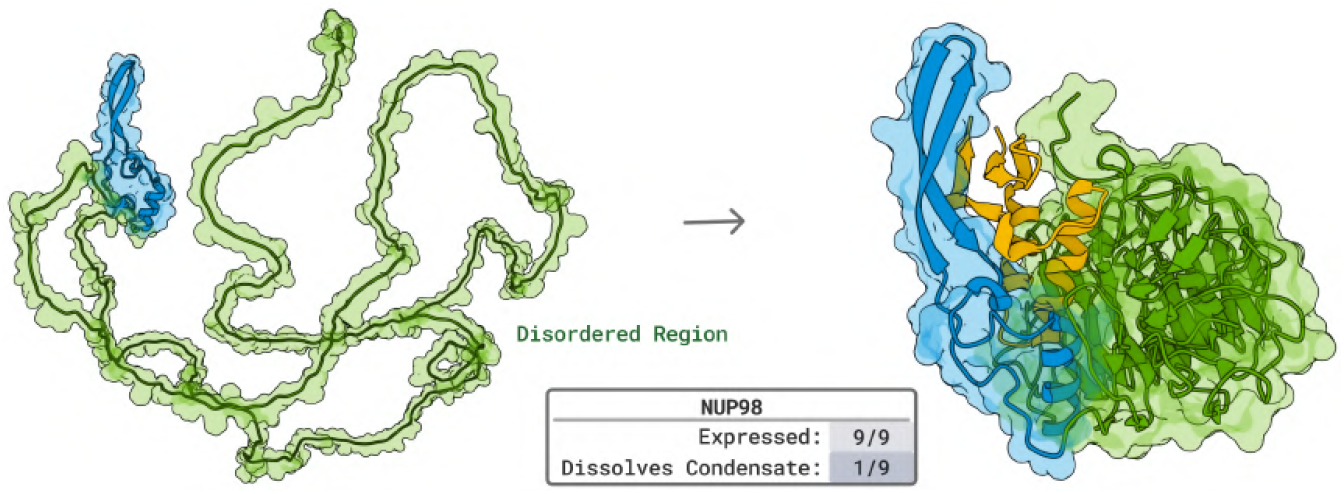

### 2.8 Designing Antimicrobial Peptides that Inhibit the GyrA to GyrA Interaction

*Experiments by Andrew Savinov, and Gene-Wei Li*.

DNA gyrase is a target of considerable interest for developing novel antibiotics. We sought to design antimicrobial peptides based on previous work in which the C-gate closure interaction between two GyrA sub-units was identified as a good target site using inhibitory protein fragments [Savinov et al., 2022, 2025]. Existing compounds, such as clinically employed fluoroquinones, function by trapping gyrase in the DNA-cleavage complex [Khan et al., 2018, Bax et al., 2019] and aminocoumarins interfere with the enzyme’s ATPase activity [Khan et al., 2018], but interfering with C-gate closure represents a complementary drug modality. Indeed, a previously reported monoclonal antibody against *Mycobacterium tuberculosis* GyrA appears to target this same site [Manjunatha et al., 2005].

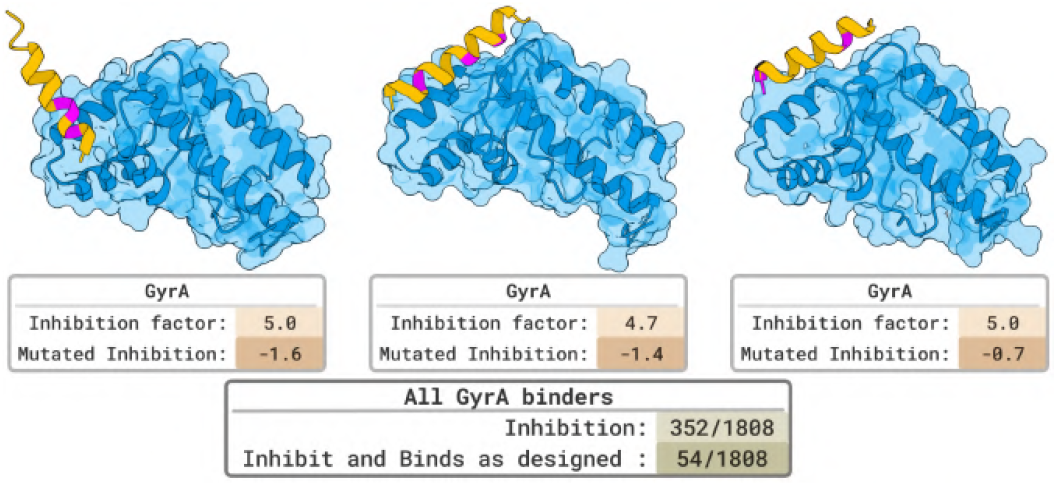

We prompt BoltzGen with residues in the GyrA C-gate closure complex as the binding site and experimentally test 1,808 designed GyrA inhibitors alongside 1,788 mutants of these designs in a massively parallel in-cell assay [Savinov et al., 2022, 2025] (Methods). The mutants were chosen to break the designed binding modes (3 alanine substitutions at the binding interface per design) as a proxy to determine whether the peptides bind GyrA as in the designed structures. These designs were tested alongside 30-aa fragments tiling across GyrA with a 1 aa step size, matching previously published work [Savinov et al., 2022, 2025], as an internal positive control for GyrA-inhibitory fragments; and also 20-aa fragments tiling (1 aa step size) across enhanced GFP (eGFP), approximately matching the average fragment length of the designs (15 ± 3.4 aa, mean ± s.d.). The eGFP fragments provided an internal negative control in the form of fragments not expected to interact specifically with *E. coli* proteins.

Across all tested GyrA binders, 352 (19.5%) substantially inhibited *E. coli* growth, comparable to known inhibitory GyrA fragments (Figure 7a-b; Methods). To quantify inhibition specificity for the designed C-gate interface, we computed Δ(Inhibition) = Inhibition(design) − Inhibition(mutated design), since true C-gate binders should lose activity upon interface mutation (Figure 7c-e). Among the 352 inhibitory designs, 54 (3.0% of total) were strongly specific (Δ(Inhibition) ≥ 2), and 99 (5.5% of total) were significantly specific under a looser threshold (Δ(Inhibition) ≥1). Overall, ∼3-6% of designs appear to inhibit growth via the intended C-gate site, while nearly 20% inhibited growth more broadly. In a control library of 20-aa eGFP fragments, only 0.45% inhibited *E. coli* growth. Relative to these nonspecific peptides, designed GyrA binders showed a ∼43-fold higher rate of growth inhibition and a ∼7-12-fold higher rate of interface-dependent, specific inhibition. Notably, 91% of interface-specific GyrA binders were so potent that they fully dropped out of the cell population upon expression, making these estimates a lower bound on inhibitory activity.

**Figure 7:**
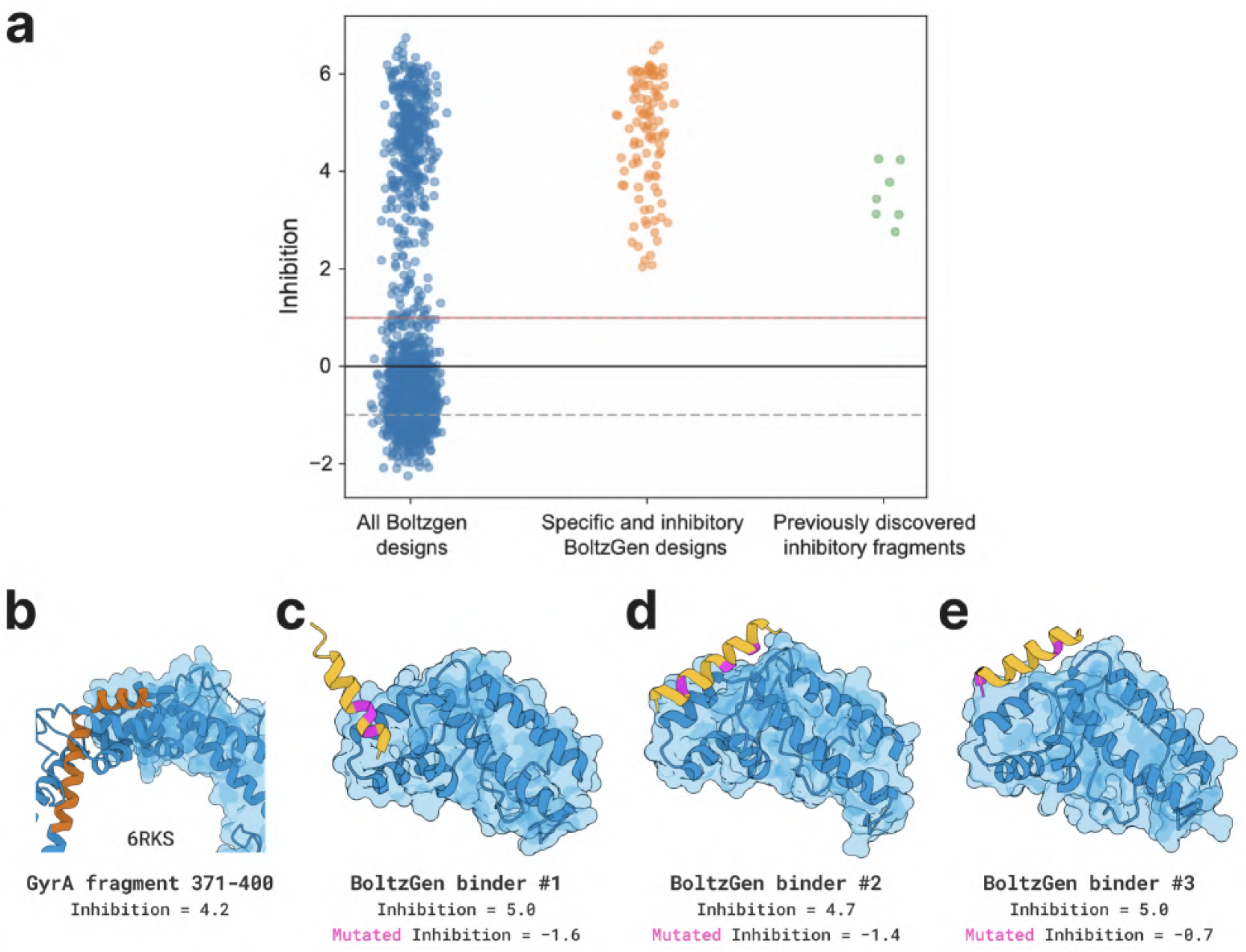
BoltzGen-designed binders are potent DNA gyrase inhibitors *in vivo*. **(a)** Massively parallel in-cell inhibition measurements (Methods) for all designed GyrA binders, the interface-specific inhibitory subset (5.5% of designs), and known inhibitory GyrA fragments targeting the same site [Savinov et al., 2022, 2025]. Reported inhibition for all fragments and 91% of interface-specific designs is a lower bound, as these peptides fully dropped out of the cell population upon expression due to strong growth inhibition. **(b)** Structure of inhibitory GyrA fragment 371-400 (orange; [Savinov et al., 2022, 2025]) bound to full-length GyrA (blue) in the GyrA dimer (PDB ID 6RKS), with *in vivo* inhibition indicated. **(c–e)** Three representative BoltzGen binders targeting this interface show strong, specific inhibition that is abolished by three alanine substitutions at the designed interface (magenta). Results are also summarized in Figure 1.

**Figure 8:**
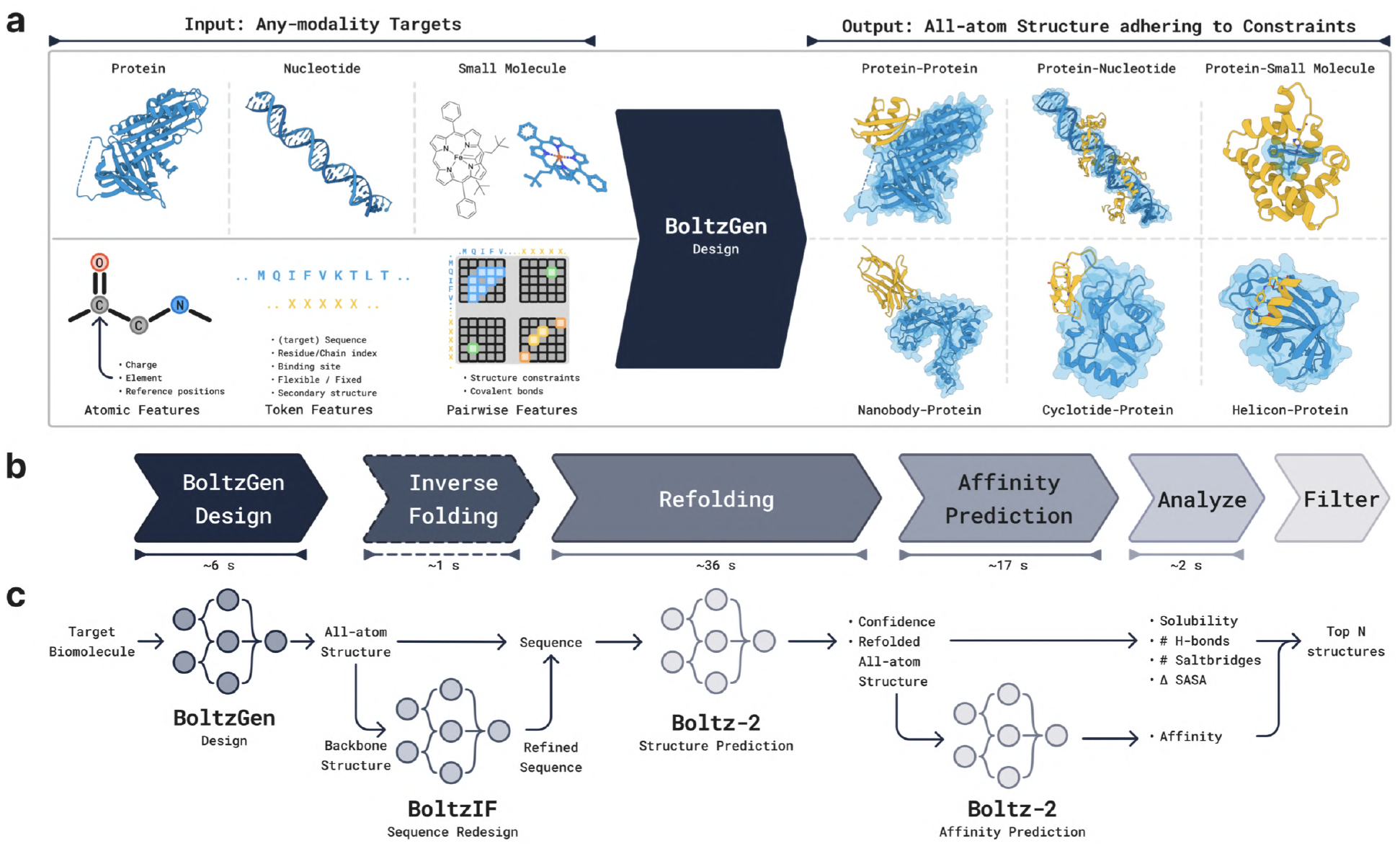
Overview of BoltzGen Pipeline: (**a**) The overall flow from target specification to binder generation. The generative model designs binders for arbitrary targets, including proteins, nucleic acids, and small molecules. It supports various constraints such as covalent bonds, structural motifs, and specific binding sites. The generated designs are passed through a filtering and ranking pipeline to produce a small, diverse set suitable for experimental validation. (**b**) Runtime per stage for a representative case with 200 total residues across the binder and target, illustrating the model’s efficiency. (**c**) A detailed breakdown of each pipeline stage, showing intermediate inputs and outputs.

## 3 Method

BoltzGen is a single all-atom diffusion model capable of performing both structure prediction and protein design. The model takes a set of molecular entities as input and outputs their all-atom three-dimensional structure. Molecular entities include small molecules, RNA, DNA, or protein sequences, along with any post-translational modifications and covalent bonds. New proteins are sampled by specifying *design residues*, for which the model generates both the all-atom structure and amino acid identities. Structure prediction and design capabilities can be exercised in tandem; for example, when generating a binder given only the sequence of the target, the model simultaneously folds the target and designs the binder’s atomic structure, producing a bound complex.

### Unified Design and Structure Prediction

The joint all-atom sequence and structure sampling ability of the model and its scalable training come from its purely geometry-based representation of designed amino-acid types. This representation encodes residue identities according to the position of the “left-over” atoms of side chains (see Figure 9). This change helps maintain a scalable architecture and its associated diffusion training process, similar to state-of-the-art biomolecular structure prediction methods.

**Figure 9:**
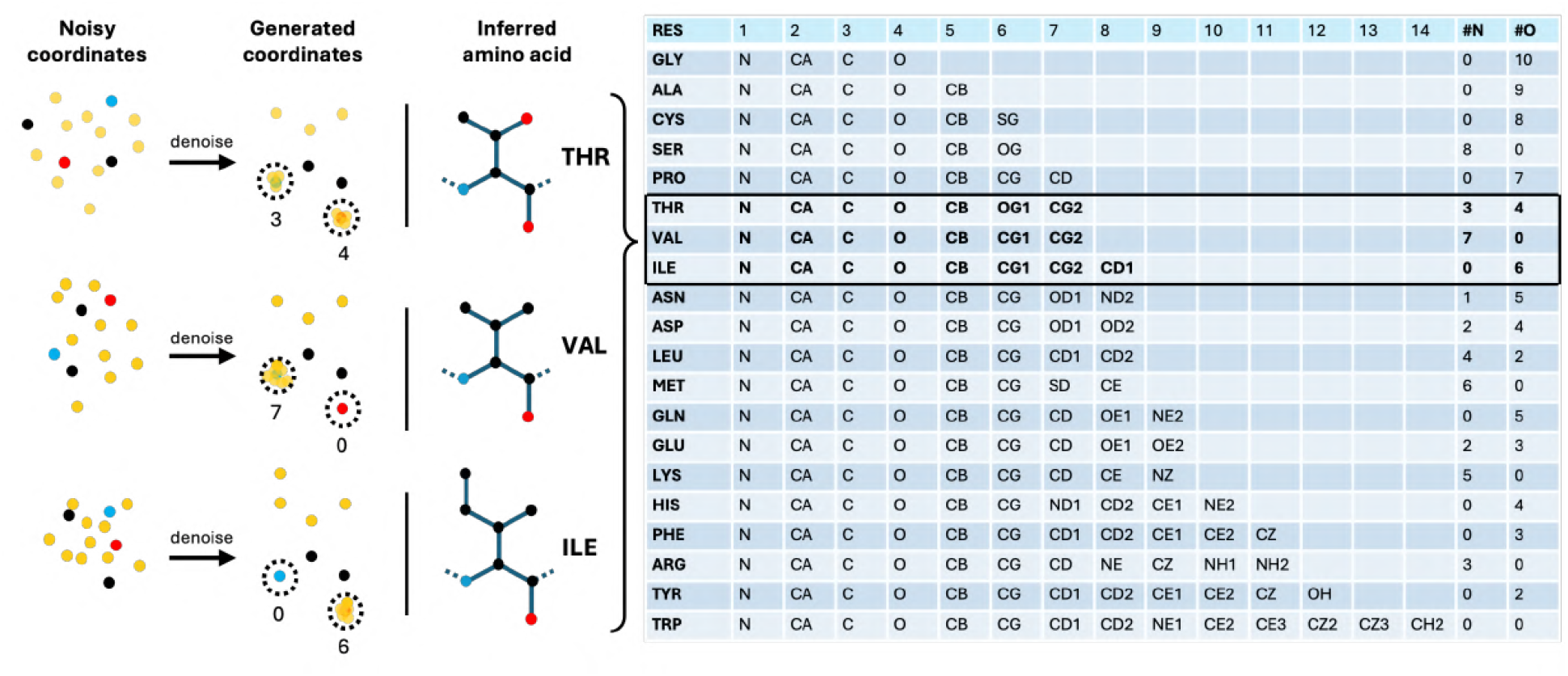
Residue Type Encoding. In BoltzGen, each designed residue is represented using 14 atoms. To determine the residue type, the model is trained to superpose a subset of these atoms onto specific backbone atoms. These auxiliary atoms act as markers and are discarded after decoding. For example, placing three atoms on the backbone nitrogen and four on the backbone oxygen is interpreted as a threonine. The atoms in positions 5, 6, and 7 are then assigned as the threonine CB, OG1, and CG2 atoms, respectively.

### Design Specification Language

Predictions can be conditioned on a broad set of additional inputs. These include standard annotations such as desired residue types and covalent bonds, as well as secondary structure identity, binding site location, and structure templates. The conditions can be incorporated with the help of a rich design specification language, allowing us to support the needs of our experimental collaborators. As a result, we can address a wide range of design challenges, including diverse modalities such as nanobodies, cyclic peptides with various cyclizations, helicons, or cyclotides.

In addition to the core generative model, we provide a comprehensive design pipeline that includes: (1) the initial generation of candidate designs, (2) optional sequence redesign through inverse folding, (3) evaluation of refolding quality and, for small molecule targets, affinity estimation, (4) ranking and filtering of designs, and (5) selection of a final candidate pool with diversity optimization.

### 3.1 All-atom Generative Model Formulation

#### Model Representations

All non-designed molecular entities provided as input to the model are represented at the atomic level, including atom positions, element types, and charge. These atoms are grouped into tokens, such as residues for proteins or nucleotides for RNA and DNA. Designed residues, which do not have a specified amino acid type during generation, use a fixed-size representation consisting of exactly 14 atoms, some of which serve as *virtual* placeholders. Once the model determines the final residue types, these virtual atoms are discarded, as done in other approaches that employ a fixed number of residues to circumnavigate the issue that a residue’s atom count is unknown before its generation [Qu et al., 2025, Chu et al., 2024, Butcher et al., 2025].

#### Geometric Encoding of Residue Type

Instead of generating a discrete residue label, the model encodes the residue identity through the geometry of the 14-atom representation (see Figure 9). Specifically, it learns to place the virtual atoms on top of designated backbone atoms to signal the intended residue type. The first four atoms in each designed residue are fixed as the backbone N, C*α*, C, and O atoms, in that order. As a result, residue types can be inferred directly from the generated structure by counting how many atoms are placed within 0.5 Å of each backbone atom. Any remaining atoms that are not superposed onto the backbone are interpreted as the side chain. For example, proline is encoded by placing 7 atoms on the backbone oxygen, while threonine is represented by placing 3 atoms on the nitrogen and 4 on the oxygen.

This geometric encoding allows the model to operate entirely in a continuous space, avoiding the need to mix discrete and continuous representations. As a result, it enables efficient joint training for both structure prediction and design tasks.

#### Diffusion Process

Because the model operates on a continuous space, we can use the same diffusion process as Boltz-2 [Passaro et al., 2025]. The only difference is that the data samples now contain additional virtual atoms for the designed residues.

Formally, let *X* ∼ *p*_data_, *X* ∈ ℝ^*N ×*3^ the 3D atomic coordinates of a training sample and let *X*_*t*_ follow the forward diffusion process

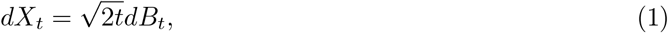

with initial condition *X*_0_ ∼ *p*_data_. For large *T, X*_*T*_ will be approximately Gaussian with variance *T* ^2^. This process can be reversed to obtain samples from the data distribution starting from Gaussian noise. Reversing the diffusion process requires a denoiser *D*_*θ*_(*x, t*; *z*), parameterized by learnable weights *θ*. The goal of the denoiser is to approximate the posterior mean

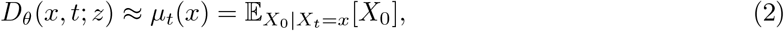

conditioned on trunk features *z*. The model is trained using a standard denoising loss

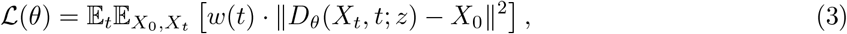

where *w*(*t*) is a weighting function. In the absence of parametric constraints, the minimizer of this loss is *µ*_*t*_, the posterior means.

### 3.2 Architecture

The model preserves the Boltz-2 architecture with some modifications to include additional conditioning inputs, as shown in Fig. 10. The model is split into two parts. The larger *Trunk* produces token and pairwise representations used to condition the *Diffusion Module*, which generates the structure. The trunk is only run once, while the diffusion module is run many times to progressively denoise the 3D coordinates of all atoms.

**Figure 10:**
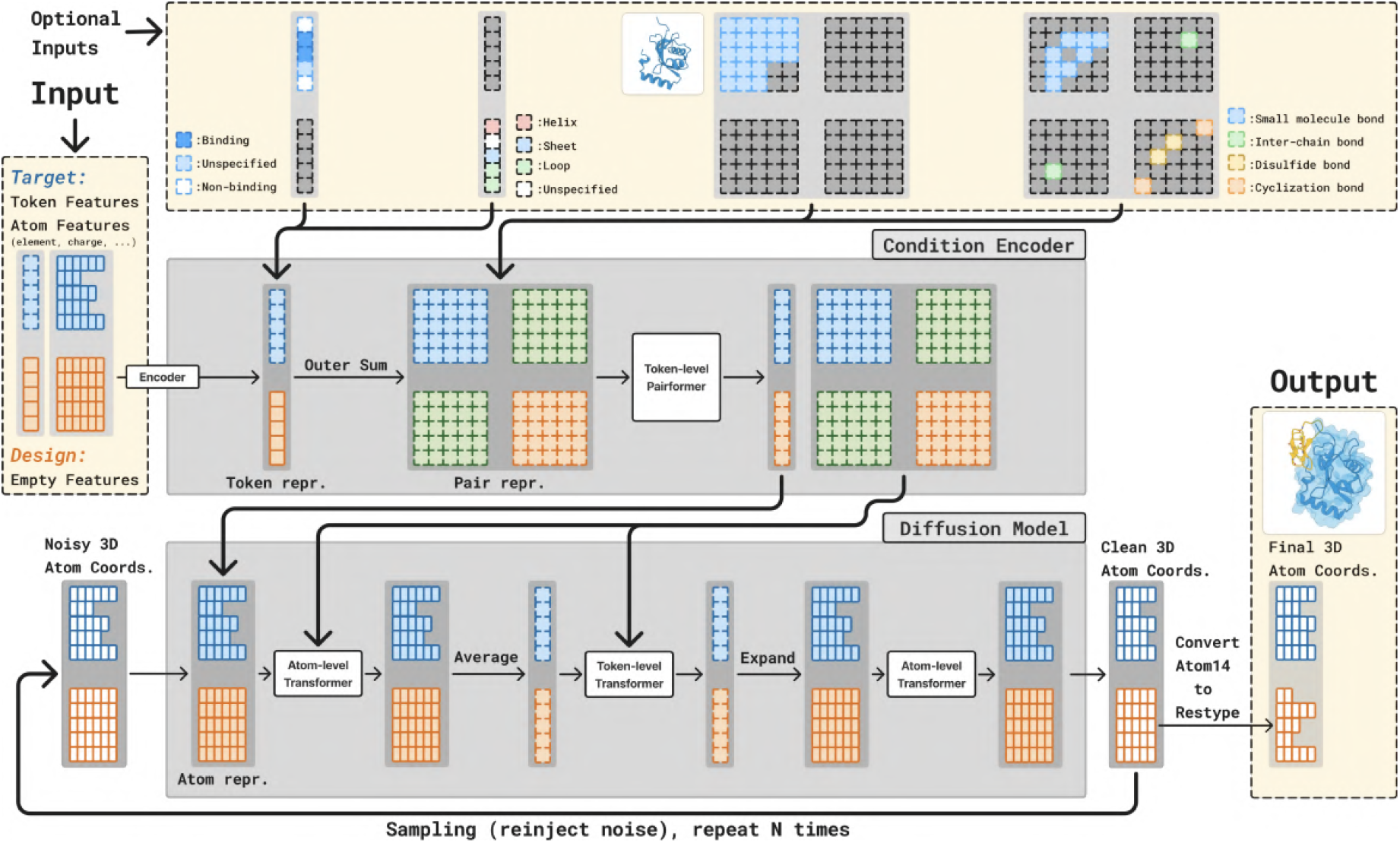
Model Architecture. The architecture preserves the main components of the AlphaFold3 and Boltz-2 architectures, including the condition encoder (trunk) and diffusion model. The main difference lies in the inclusion of design tokens and additional inputs such as binding site and target structure.

#### Trunk

The trunk operates on tokenized structures, where proteins are tokenized into amino acids, RNA and DNA into nucleotides, and small molecules into atoms. Each token consists of a group of atoms with associated features such as charge and element type, along with token-level attributes including residue index, amino acid type, and a flag indicating whether the residue is designed. All features are encoded into vector representations. Atom-level embeddings are averaged to produce token-level vectors, and a pair representation is constructed using an outer-sum of the token embeddings. Both token and pair representations are then passed through a PairFormer stack. Each PairFormer block includes triangle multiplicative and triangle attention layers that update the pair representations, along with a transformer layer that updates the token representations using the pair features as the attention bias.

#### Diffusion Module

The diffusion module takes noisy 3D atomic coordinates as input and predicts denoised coordinates. It uses a standard transformer architecture that operates on both atom and token levels, consisting of 3 atom-level layers, followed by 24 token-level layers, and concluding with another 3 atom-level layers. The atom-level layers utilize sequence-local attention. Transitions between atom and token levels are handled by averaging or expanding the representations. Conditioning information from the trunk is incorporated by adding expanded token-level features to the atom-level input representations, and by biasing the attention layers based on the pair representation.

As in Boltz-2, AlphaFold3 and EDM [Karras et al., 2022], the diffusion module is preconditioned to parameterize the denoiser *D*_*θ*_,

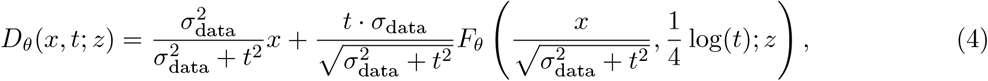

where *F*_*θ*_ is the diffusion module. This parameterization is derived in [Karras et al., 2022] so that (1) the inputs to the network *F*_*θ*_ have unit variance (2) the effective training target of *F*_*θ*_ has unit variance and (3) the scalar multiplier of *F*_*θ*_ in Eqn. 4 is minimal.

#### Controllability

Several additional inputs can optionally be provided to BoltzGen to steer the generation process according to user-specified requirements. Figure 10 shows where these inputs are integrated into the architecture, and Figure 11 illustrates the resulting expressive design specification language.

**Figure 11:**
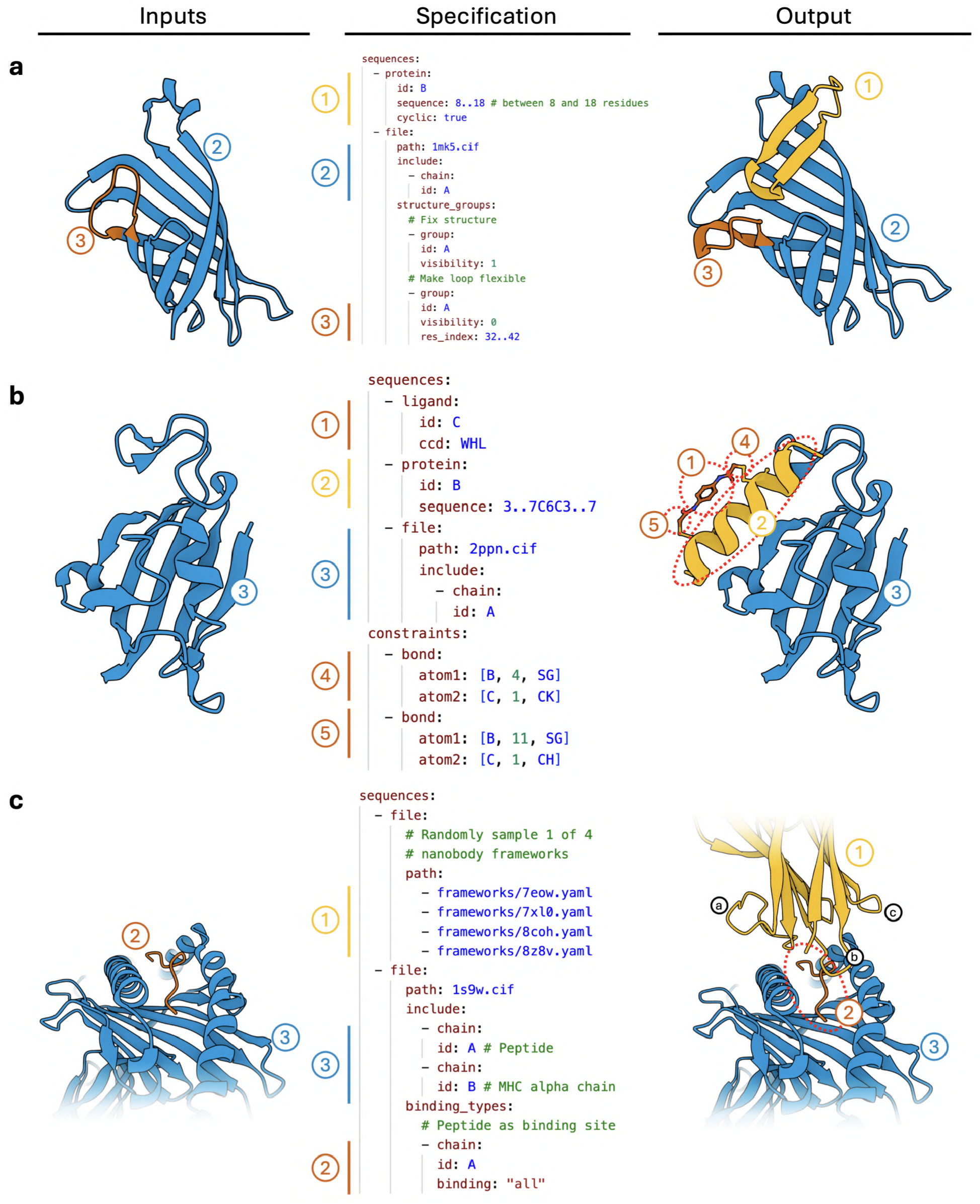
Design Specification Language. 3 examples of how the BoltzGen design specification can be used to solve different design tasks. (**a**) Designing a cyclic peptide against streptavidin. Part of the target structure is left flexible (3, orange) and is predicted to change conformation upon binding the design (1, yellow). (**b**) Designing a helicon binder. The helicon is created by including the staple molecule (1, WHL) and specifying covalent bonds between two cysteines in the design and the staple (4,5, orange). (**c**) Designing a nanobody against a peptide-MHC complex. The peptide (2, orange) is specified as a binding site, and the designed regions are limited to the 3 CDR loops (a,b,c) of the nanobody (1, yellow). The nanobody frameworks are themselves yaml files that specify the PDB structure to use and which are parts to design. With multiple specifications, a random one is sampled for each design.

1. **Covalent Bonds**. Covalent bonds can be specified between individual atoms, in which case the identity of the residues that contain the bound atoms must be specified.
2. **Structure Conditioning**. Parts of the structure can be specified to the model via pairwise distances, for example to perform motif scaffolding. All structures are specified in *structure groups*, where all pairwise distances between atoms in the same structure group are fixed, but not across groups. For example, during nanobody design, the nanobody framework structure and the target structure can be fixed but their relative positions to each other can be left unconstrained by putting them in different structure groups.
3. **Binding Site**. Residues can be specified as binding, or not-binding. The model will try to place designed residues close to binding residues and away from non-binding residues.
4. **Secondary Structure**. Designed residues can also be specified to be part of alpha-helices, beta-sheets, or coils.

Covalent bonds, specified as a matrix of pairwise bonds, and pairwise distances for structure conditioning are both encoded and added to the trunk’s input pair representation. Binding and secondary structure labels are incorporated into the input token representation.

These conditioning options can be used to control the model and address a variety of design tasks. For example,

1. **Cyclic Peptides** can be designed by specifying a covalent bond. This includes disulfide-stapled peptides, head-to-tail cyclic peptides, and any other type of cyclization.
2. **Helicons** can be designed by including a staple molecule in the model’s input and enforcing covalent bonds between the staple and the sulfurs of two cysteines in the design.
3. **Nanobodies** can be designed by enforcing the design to adhere to a given template, allowing the model to generate the CDRs as well as place the scaffold in relation to the target.

Figure 11 also illustrates how to realize these examples using the BoltzGen design interface.

### 3.3 Training

The model is trained with a diffusion objective with a mixture of experimental and self-distilled biomolecular structures. This data is then randomly cropped and employed in a diverse set of tasks by randomly selecting parts of the structure to be designed or conditioned on. The procedure is described in Figure 12. This conditioning sampling process, combined with the diffusion objective, supervises BoltzGen to simultaneously learn folding, binder design, motif scaffolding, and more, resulting in a universal binder design model that maximally extracts information from the data.

**Figure 12:**
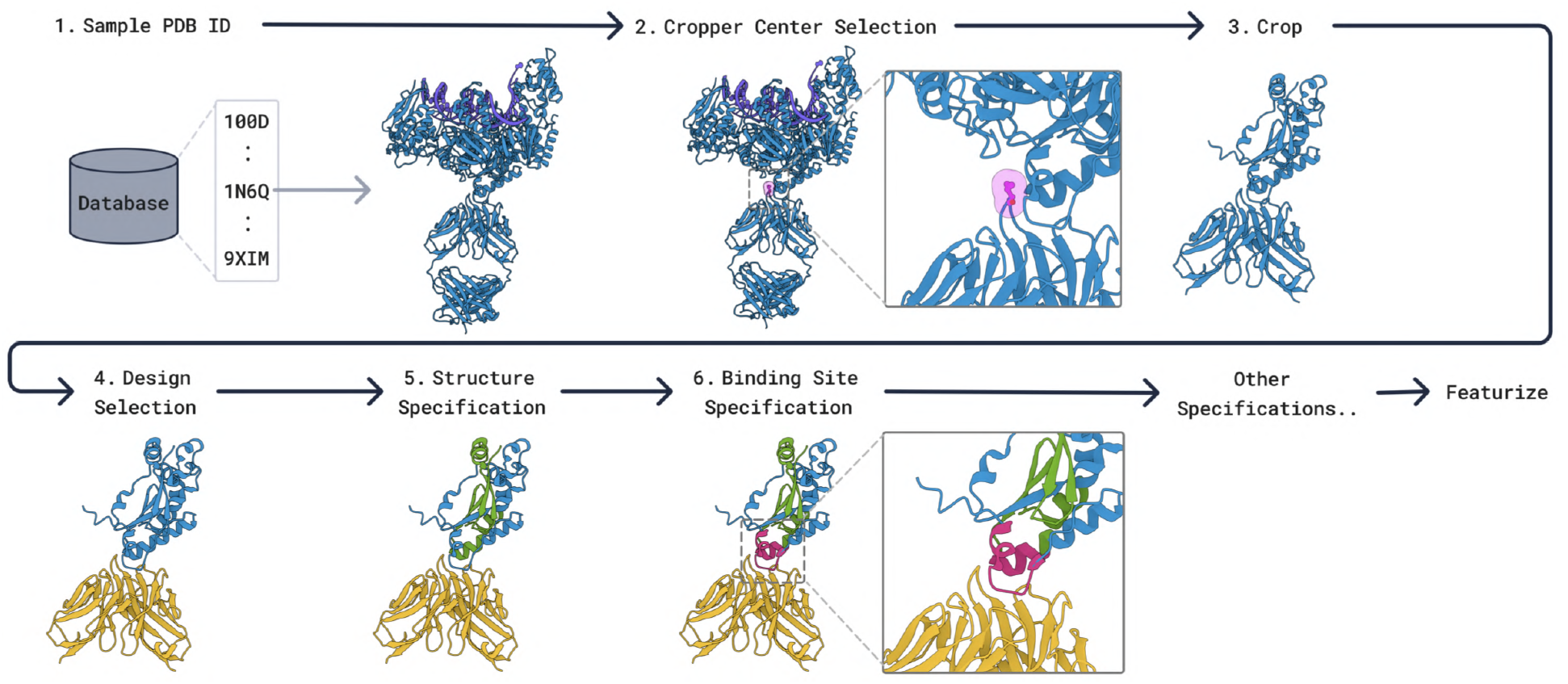
Crop, Selection, and Specification. During training, we sample one PDB structure from the database and crop it using our cropping algorithm 4. On top of this cropped structure, we optionally select chains that will be designed (yellow). Diverse conditions, such as fixed target substructures (green), binding sites (red), and more, are also optionally specified. Cropped, selected, and specified structure is then featurized to be supplied to the model.

**Figure 13:**
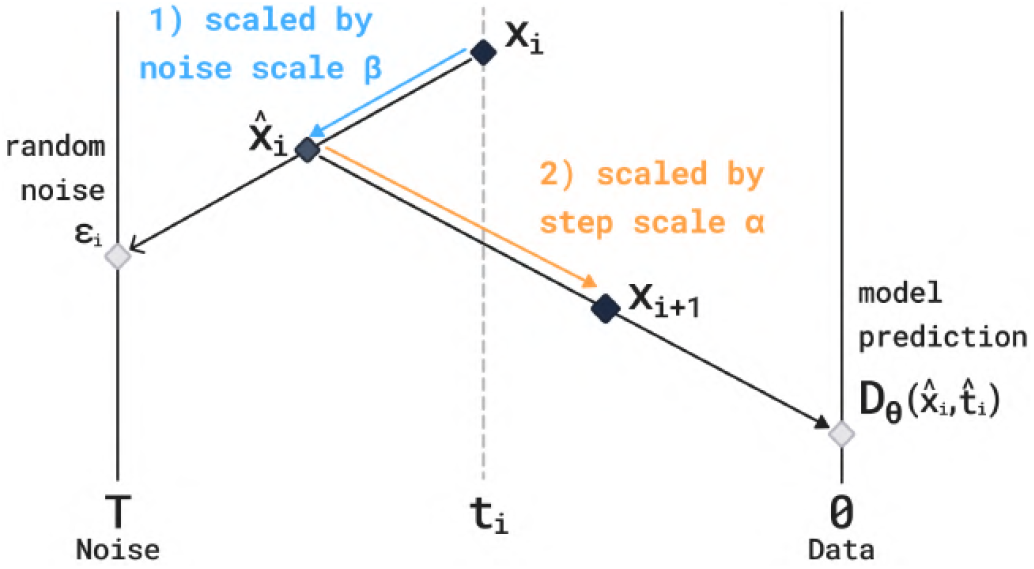
The EDM Sampler steps toward noise and then toward the model prediction in each denoising iteration. The magnitude with which to make these steps can be scaled by ***β*** (noise) and ***α*** (prediction). Using ***α*** ≠ 1, ***β***≠ 1 no longer samples the training distribution, but is a heuristic scaling to sample more diverse (higher ***β***, lower ***α***) or more designable (lower ***β***, higher ***α***) proteins.

**Figure 14:**
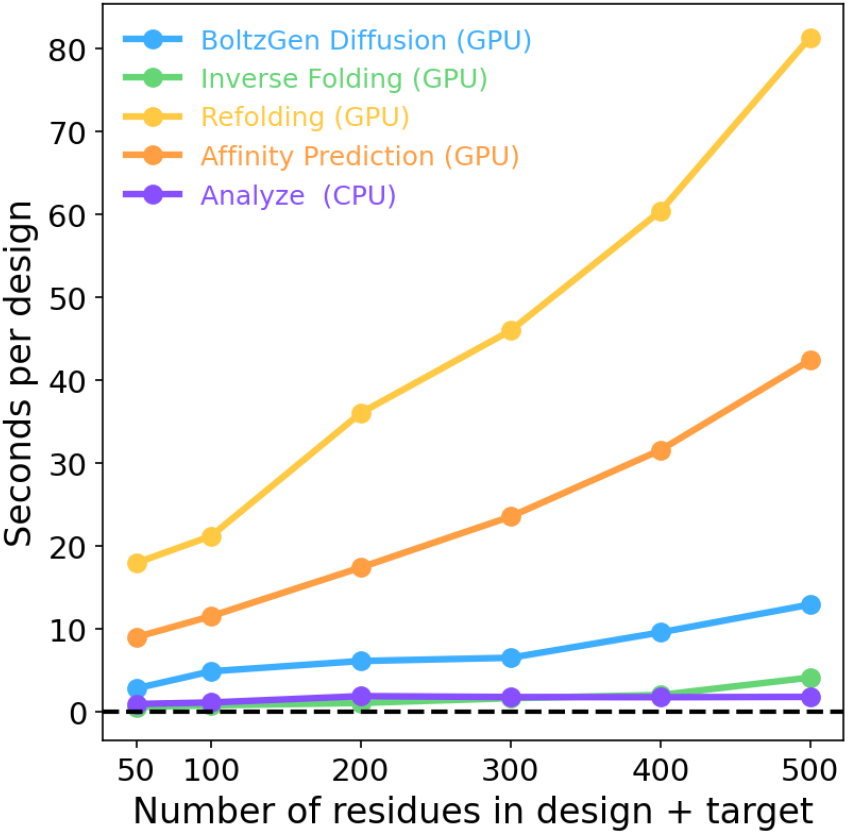
Time consumption of each step in seconds per design for different numbers of residues in the design + target complex (on a single NVIDIA A100 GPU).

#### Training Data

Our data pipeline mostly retains the datasets used in Boltz-2 [Passaro et al., 2025], while adapting the sampling procedure for the task of biomolecular design. Specifically, we use experimental structures from the Protein Data Bank (PDB) [Berman et al., 2000], as well as self-distilled structures from AlphaFold2 (AFDB), as well as Boltz-1 (for protein-ligand, RNA, and DNA-protein structures) (see Appendix B.1 for dataset details). Our data also differs from Boltz-2 by not including the upsampled antibody and TCR datasets, since including them reduces generation diversity.

#### Cropping

The crop size for folding is up to 768 residues, as done in [Abramson et al., 2024, Passaro et al., 2025]. Crop size for generative tasks (binder design, motif scaffolding, and unconditional design) is reduced to 512 residues, to accommodate augmented fake atom representations (Appendix B.2).

#### Diffusion Objective

The loss used to train the model is

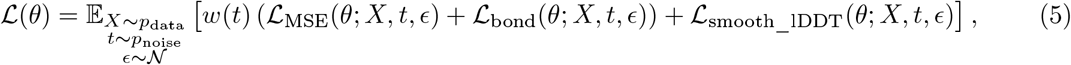

where *p*_noise_ is described below, *ϵ* is standard isotropic Gaussian noise, 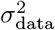 is the variance of the data, ℒ_MSE_,ℒ_bond_, and ℒ_smooth_lDDT_ are the three components of the loss described below and the weighting is defined as

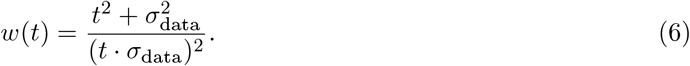

Let 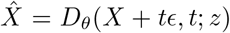 be the output of the denoiser. The MSE loss is a weighted version of the denoising loss including a rigid alignment of the denoiser output and target,

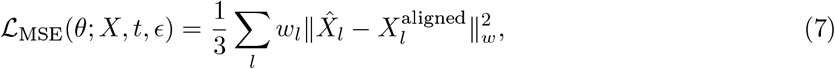

where *l* iterates over atoms in the structure, *X*^aligned^ is rigidly aligned to 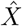, and *w*_*l*_ upweighs nucleotide and ligand atoms,

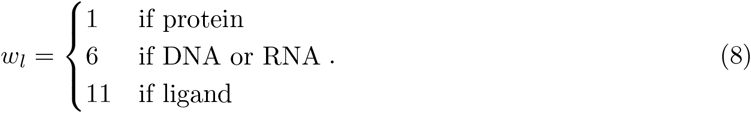

The bond loss encourages bond length correctness,

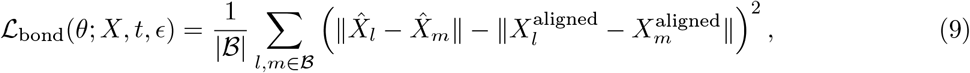

Finally, ℒ_smooth_lDDT_ is described in [Abramson et al., 2024] and is a smooth version of the lDDT which can be directly optimized.

The noise sampling distribution *p*_noise_ used during training is

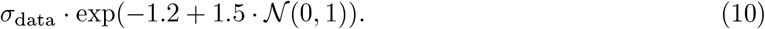

#### Training Tasks

We train BoltzGen on a number of tasks (Appendix B.3 for more details) under the different forms of conditioning above to obtain a general and controllable algorithm. This not only results in generality but could also improve performance for individual tasks as the model is exercised on the data in more contexts, increasing the number of examples to extract generalizable patterns from and learn to emulate physics. Each task is distinguished by which parts of a cropped structure are chosen to be designed. These selected parts are represented using the fixed-size representation with virtual atoms described above, and their residue and atom types are masked.

1. **Folding**. No design residues are specified, corresponding to structure prediction.
2. **Binder Design**. One protein chain in the target structure is chosen to be designed. This corresponds to binder design against whatever other biomolecules are in the target structure, which could be a small molecule, DNA, RNA, or another protein. Sometimes, only the interface of the protein chain with the rest of the structure is designed, corresponding to binder design with a supplied scaffold.
3. **Motif Scaffolding**. Either a crop of the target structure is chosen to be designed, or everything but the crop is designed. This corresponds to completing a scaffold in the first case, and scaffolding a motif in the second case.
4. **Unconditional Design**. Everything is chosen to be designed, corresponding to unconditional protein generation.

Each task is sampled with some probability during training as long as the target structure is appropriate for the task. For example, the binder design tasks are not sampled for structures that contain a single protein monomer.

In addition to the tasks, we also sample different conditioning to supply the model:

1. **Structure Conditioning**. We randomly choose portions of the target structure to supply as input to the model based on crops. These are also randomly grouped together.
2. **Binding Site**. We randomly annotate residues as being part of a binding site or not, based on proximity to any portions of the training structure selected to be designed.
3. **Secondary Structure**. We annotate random residues by their secondary structure.

See Appendix B.3 for full details on the various schemes we use to construct the tasks and sample the different conditioning inputs to give the model.

##### Algorithm 1 EDM Sampler

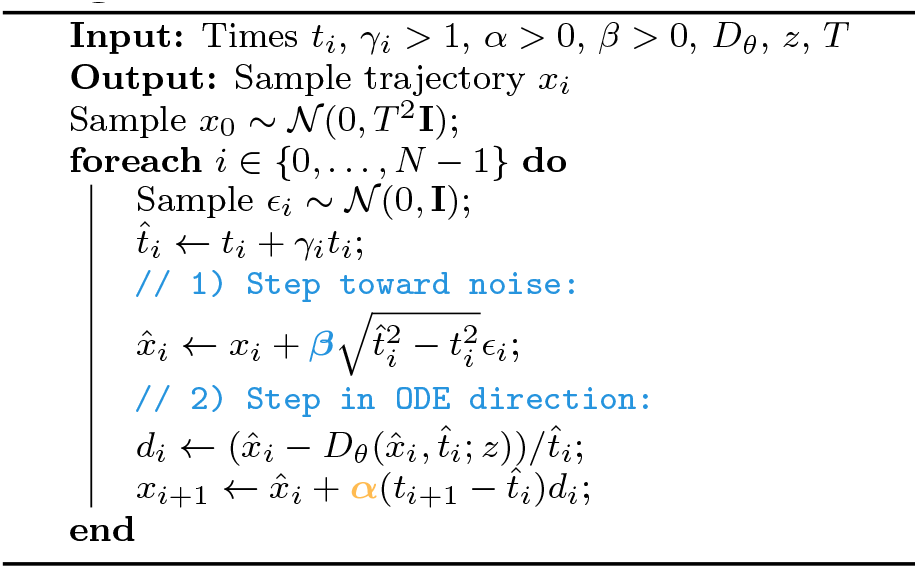

### 3.4 Generation

To sample from the model, the inputs are first passed through the trunk to obtain the token and pair representations *z* that condition the diffusion module. The diffusion module then generates a structure *x* using the stochastic sampler from [Karras et al., 2022]. This sampler starts from random noise and alternates between adding noise and denoising.

The sampler makes use of the probability flow ODE, which in the case of our forward process is,

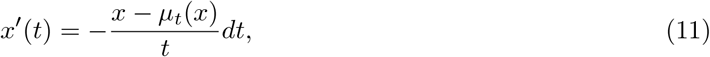

for *t >* 0, where *µ*_*t*_ is the posterior mean in Eqn. 2 that is approximated by the denoiser *D*_*θ*_. Note that this ODE runs forward (towards noise). Let *F* (*x, t, s*), be the associated flow map, i.e. if *x*(*t*) is a solution to the above ODE, then

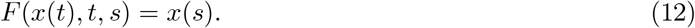

The probability flow ODE satisfies that *F* (*X*_*t*_, *t, s*) has the same marginal distribution as *X*_*s*_. In particular, *F* (*X*_*t*_, *t*, 0) gives samples from the data distribution for any *t >* 0. Rather than simulate the ODE the whole way, the sampler in [Karras et al., 2022] interleaves noising steps to add stochasticity based on the observation that 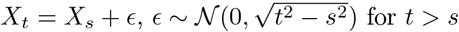. Hence, alternating between adding noise *ϵ* and applying the flow map *F* always gives samples with the correct marginals. The algorithm is given in Alg. 1 which additionally makes use of a step scale *α* and noise scale *β*. To sample more high-designability but lower diversity binder structures, we can increase the step scale or decrease the noise scale (and the opposite to obtain more diverse proteins). When drawing many samples, we vary the step scales and noise scales for each generated protein.

#### Dilated schedule

The time schedule *t*_*i*_ used in AlphaFold3 and Boltz is,

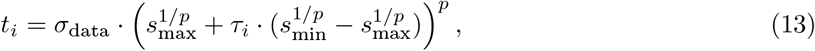

where *σ*_data_ = 16, *s*_min_ = 4 ·10^−4^, *s*_max_ = 160, *p* = 7, and 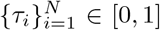 is a sequence of steps. By default, *τ*_*i*_ = *i/N* .

Because of our geometric encoding, we found that the amino acid types of designed residues were determined within a short window, approximately at *τ*_*i*_ ∈ [0.6, 0.8]. In order to spend more function evaluations when generating the residue types, we therefore use a *dilated* schedule where the interval [0.6, 0.8] is stretched out. Concretely, to dilate an interval [*τ*_*s*_, *τ*_*e*_] by 1 < *λ* < 1*/*(*τ*_*e*_ − *τ*_*s*_), we map step *τ* ∈ [0, 1] according to

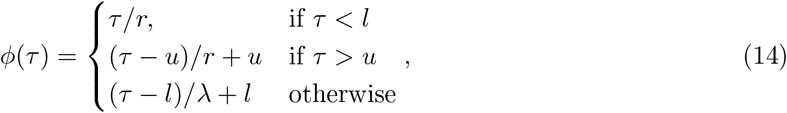

where *r* = (1 − *λ* · (*τ*_*e*_ − *τ*_*s*_))*/*(1 − (*τ*_*e*_ − *τ*_*s*_)), *l* = *r* · *τ*_*s*_, *u* = *l* + *λ*(*τ*_*e*_ − *τ*_*s*_). In practice, we choose *λ* = 8*/*3, *τ*_*s*_ = 0.6, and *τ*_*e*_ = 0.8, with *N* = 300 total function evaluations which we found to work well in experiments.

### 3.5 BoltzGen Pipeline

On top of the generative model, we run a computational pipeline to filter from the potentially thousands of designs sampled by the model down to the most promising candidates. The pipeline consists of the following stages.

1. **BoltzGen Diffusion (GPU)**. Given a design specification, we generate a large number of designed binders with BoltzGen.
2. **Inverse Folding (GPU)**. We optionally inverse fold the designed binders. This step tends to create sequences that are more likely to be predicted to refold into the designed structure. It is also likely to improve solubility (the inverse folding model was only trained on soluble proteins). We use BoltzIF (detailed below), which is similar to SolubleMPNN [Goverde et al., 2024].
3. **Folding (GPU)**. We predict the structure of the design in complex with the target using Boltz-2, which is provided the template of the target structure (produced in step 1) and no MSAs. We compute the RMSD between the predicted structure and the structure produced by step 1 for later filtering (if the refolded structure is similar to the designed structure, the design is more likely to be a binder). This also produces confidence metrics (pTMs, pAEs) used in filtering. In scenarios where we design globular proteins, we perform an additional refolding step where we only predict the structure of the designed binder (in absence of the target), and compute the same RMSD metrics. As detailed in D, we found this necessary to assess whether the designed protein’s structure can be achieved in absence of the target, indicating that the design can express well and fold on its own.
4. **Affinity Prediction (GPU)**. When designing proteins that bind small-molecules, we predict their affinity using Boltz-2’s affinity module.
5. **Analyze (CPU)**. We compute physics-based metrics that provide information about the binder-target interaction strength and developability properties of the design. Here we describe those that are used in the filtering step by default. A complete list is in Appendix B.4. We count the hydrogen bonds and salt bridges between design and target and assess the buried surface area (how much contact is there and how much shape complementarity) between the design and target. Additionally, we compute a sequence based solubility metric and the area of the largest hydrophobic patch on the surface of the designed protein (large hydrophobic patches can cause protein aggregation and difficulty during purification or expression).
6. **Filter (CPU)**. Based on the metrics computed in the previous stage, we produce a ranking of the designs (detailed below). This ranking is used in a quality-diversity algorithm (detailed below), which returns a final set of filtered (to a desired budget *k*), ranked, and diversity optimized candidates. This step takes around 20 seconds, independent of the number of designs.

#### Inverse Folding Model

Our inverse folding model is largely similar to SolubleMPNN [Goverde et al., 2024]. The main differences are that (1) we use 6 encoder layers instead of the 3 of SolubleMPNN, (2) we exclude fibril proteins from the training set next to excluding transmembrane proteins and (3) we train on structures cropped to 1024 amino acids rather than excluding larger instances. We verify that our inverse folding model performs similar to ProteinMPNN and SolubleMPNN in Section C.4.

#### Ranking

To rank designs, we first compute two groups of metrics that we expect to correlate with experimental success: Boltz-2 confidence scores [Passaro et al., 2025] and interaction-type scores such as the number of hydrogen bonds. These are aggregated into a single quality score *q* by taking the worst rank across all metrics (Algorithm 2). Metrics are given varying importance by weighting their respective ranks, where the weights are calibrated on a benchmark of validated binders (see Appendix C.2). This prioritizes designs with the best worst-case performance across all metrics.

##### Algorithm 2 Top-*k* binder designs selection by worst weighted metric rank

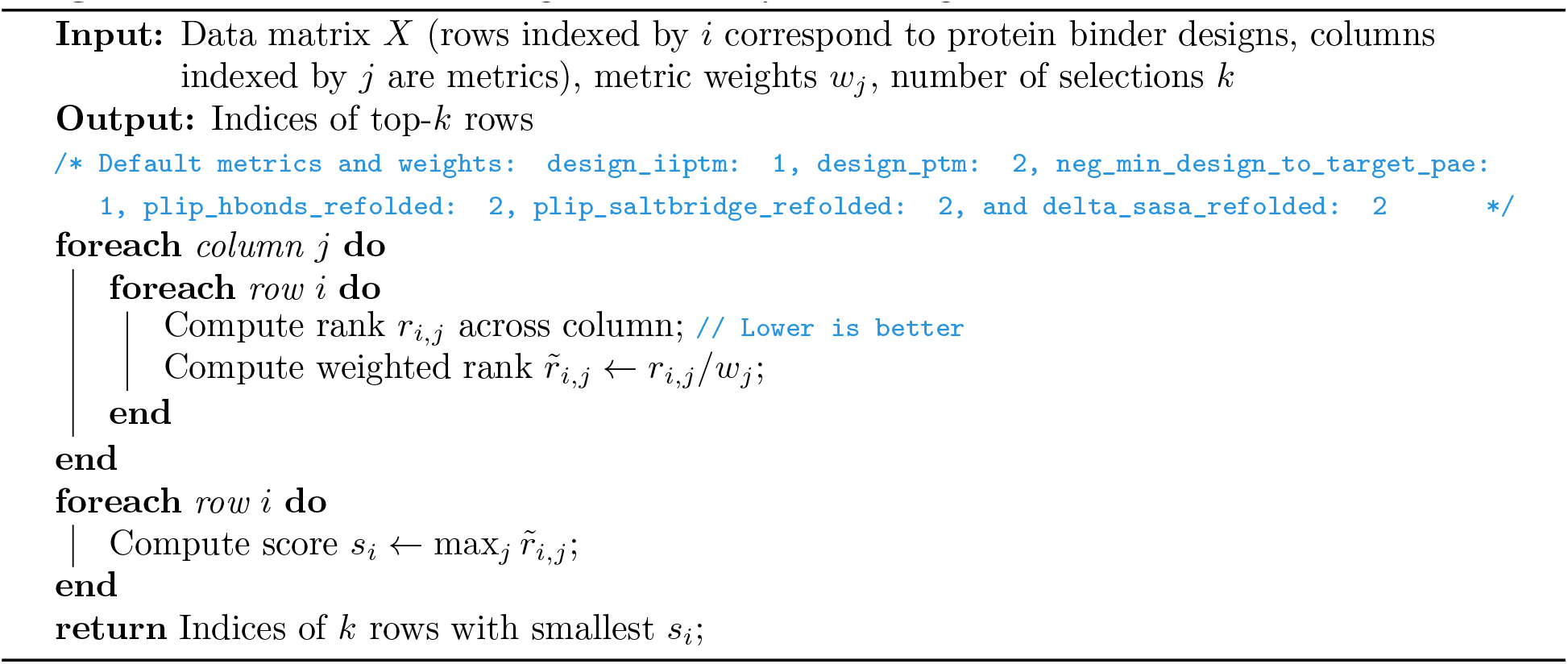

#### Quality-Diversity Selection

In order to select a *diverse* set of high-ranking designs, we use a greedy selection algorithm described in Alg. 3. Each design *x* produced by the model is ranked according to the aggregated score *q*(*x*) introduced above. For a given set of candidates *S*, we also measure the similarity of design *x* to all designs in the set *S* based on a mixture of the TM-score and sequence similarity,

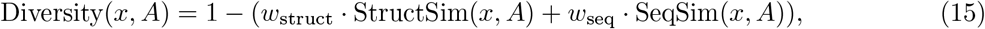

where

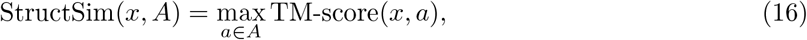

and

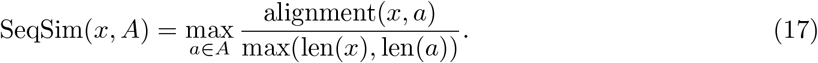

The algorithm then proceeds by greedily adding designs that maximize a combination of quality and diversity with respect to the current set of candidates.

##### Algorithm 3 Greedy Quality-Diversity (QD) Selection

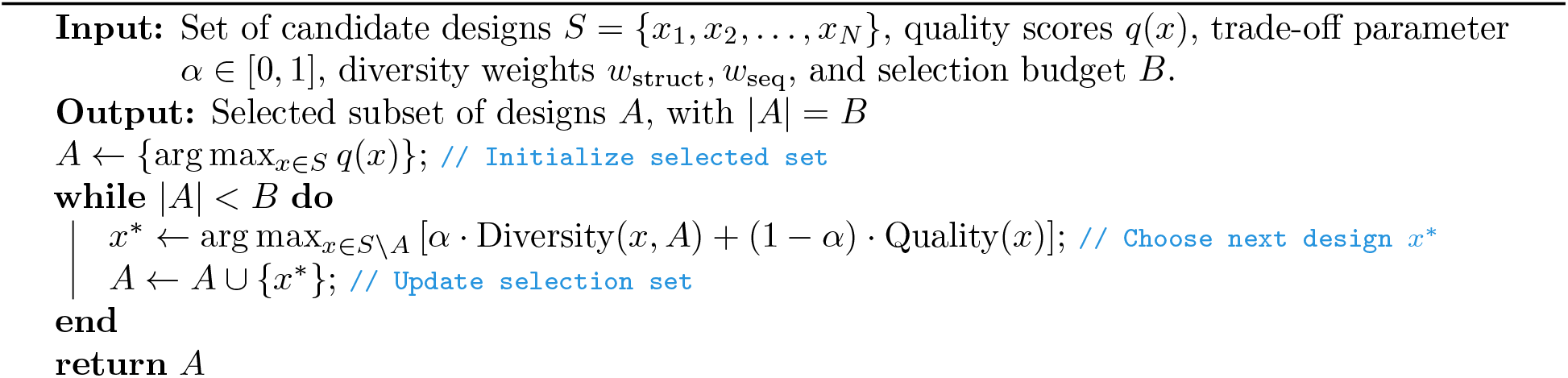

## 4 Computational Results

### 4.1 Structure Prediction

BoltzGen’s development was driven by the hypothesis that designing high-affinity binders requires strong structure prediction and reasoning capabilities. Expressive features that allow for accurate structure prediction are also crucial for design and enable the model to place atoms that tightly interact with the target. Thus, we assess BoltzGen’s structure prediction performance.

Figure 15 shows that BoltzGen’s folding performance matches Boltz-2 on its test set (minus 187 complexes that did not fit on a 40 GB GPU). This test set is based on clustering sequences with a 40% similarity threshold (details in B.1.2).

**Figure 15:**
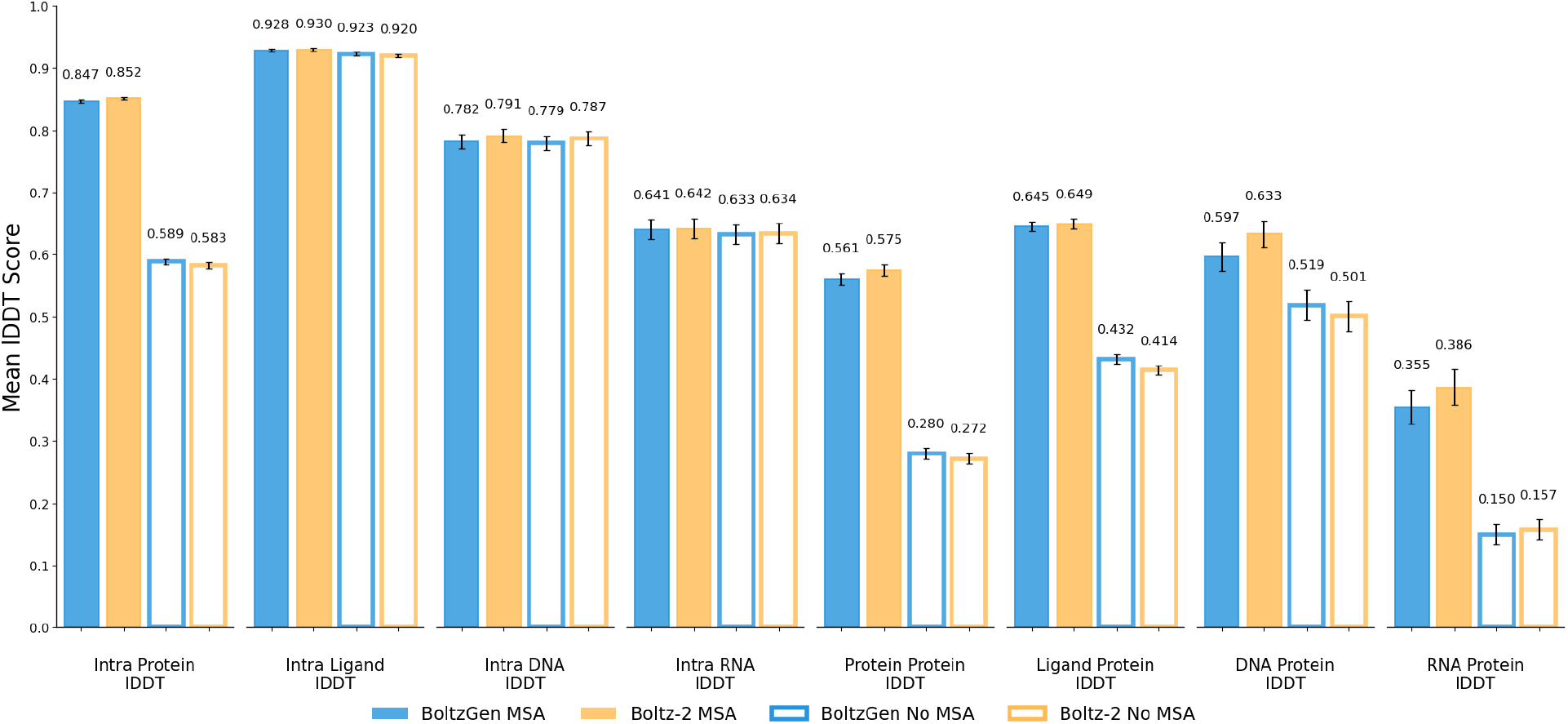
Structure Prediction Evaluation. We reason that structure-based binder design requires strong structure prediction and reasoning capability. BoltzGen can perform design *and* folding. Its folding performance matches Boltz-2. Shown is the best lDDT out of 5 diffusion samples for each complex.

### 4.2 Computational Binder Design

In a comparison with RFdiffusion [Watson et al., 2023] and its extension RFdiffusionAA [Krishna et al., 2024] we aim to assess the degree to which the models ignore the target and produce the same structures independent of their conditioning. To evaluate a model’s target dependence, we collect a set of targets and draw one binder design per target and filter that set, only keeping the designs that match the model’s generated structure after refolding with Boltz-2 (based on an RMSD threshold of 2.5 Å). We then assess the diversity of the resulting set as the Vendi score [Friedman and Dieng, 2022] with TM-score as the similarity kernel. Methods that tend to produce the same structures irrespective of the target will obtain a lower diversity.

We carry this evaluation out for monomeric proteins and small molecules as targets. The monomer set is selected based on sequence similarity clustering and June 2023 as a date cutoffs (details in B.1). The small molecule set is a random sample from all ligands that satisfy standard drug-like property criteria (Lipinski’s rule of five [Lipinski et al., 2001]) and have a Tanimoto similarity of less than 0.35 to any small molecule with the same date cutoff. The comparison in figure 16 shows that BoltzGen “pays more attention” to the target.

**Figure 16:**
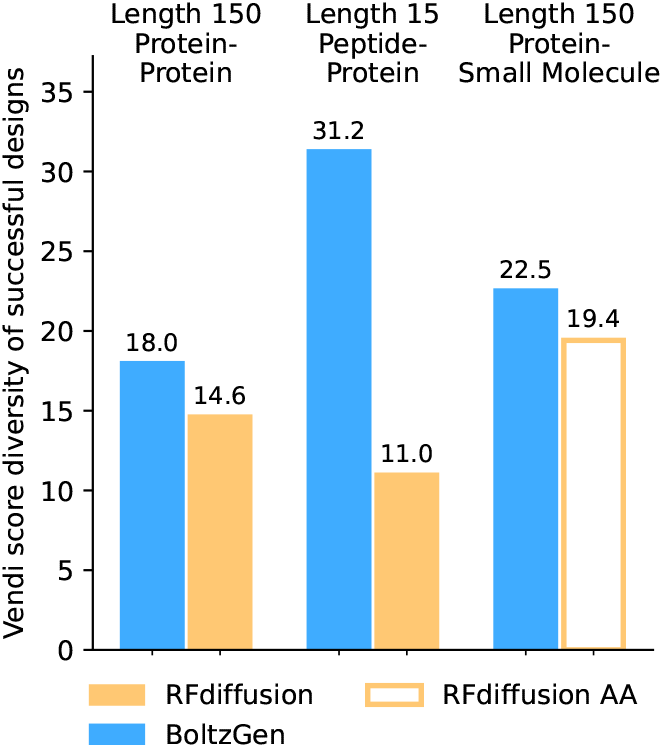
Target conditioning quantification. The diversity of successfully refolded complexes when designing a single binder against 110 targets. This assesses the degree to which the models are conditioned on the target instead of generating designs independent of the target.

## 5 Limitations

For therapeutic development, generating high-affinity binders is only the first step. Whether a design will achieve the intended function depends on a range of additional properties, including selectivity, developability, and the precise characteristics of the target. While predictors for these could be included in BoltzGen’s filtering steps, the *selective* binding problem may be particularly ripe for direct integration into the generative process. Classifier-free guidance techniques could be used for combining the scores of BoltzGen with different targets as conditioning to achieve guiding toward one target while steering away from off-targets. Alternatively, BoltzDesign1 [Cho et al., 2025a] could be used to suggest mutations that prevent binding to off-targets.

More specific to BoltzGen, we note that there is a memorization issue when designing binders of length 73-76. For certain protein targets, BoltzGen’s generation diversity collapses in this length range and it nearly exclusively samples ubiquitin as the binder. For future BoltzGen training runs, ubiquitin should be downsampled (appears more than 900 times in the PDB). More details about this ubiquitin memorization is provided in Appendix C.6.

Lastly, we comment on how there is a tendency in the field to claim that binder design models are “zero-shot” and “plug-and-play” solutions without a chance for failure. We do not make this claim and encourage users to use BoltzGen thoughtfully, carefully inspect the generated structures, and potentially rerun the pipeline multiple times, first at smaller and then larger scales. BoltzGen’s rich design specification language provides a large degree of control that should be experimented with for optimal results.

## 6 Conclusion

BoltzGen is a general-purpose generative model for biomolecular binder design, supporting a broad range of modalities, including proteins, peptides, nanobodies, and related modalities, which can be designed to target virtually any biomolecule, such as proteins, small molecules, and nucleic acids. While the core of BoltzGen is a diffusion generative model, we provide it as part of a broader framework that includes tools for specifying design tasks, generating candidates, and filtering, ranking, and optimizing for diversity. This makes it a practical, end-to-end solution for several binder design problems.

We release the entire BoltzGen package under the MIT license, including model weights, training and inference code, and a user-friendly pipeline for design and evaluation. By providing a complete and accessible solution, we hope BoltzGen serves as a practical tool and a foundation for future work in general-purpose biomolecular design.

## Acknowledgments

We would like to thank Rachel Wu, Sergey Ovchinnikov, Wenxian Shi, Tally Portnoi, Umesh Padia, Peter Mikhael, Edwin Neumann, Christopher Garcia, Kevin Jude, Juno Nam, Cora Wendlandt, Dane Wittrup, Bradley Pentelute, Jigar Patel, Reda Rawi, Sarah Fahlberg, Kellon Belfon, Giannis Daras, Sushil Mishra, Mathai Mammen, Noah Getz, Bowen Jing, Donovan Chin, Thomas Schwartz, Jeremie Alexander, Jonathan Stokes, Robert Hurt, Matthew Shoulders, Robbie Wilson, Aneesh Karatt-Vellatt, and Soojung Yang for insightful discussions and feedback. We thank Jiri Damborsky for providing insights into early versions of the model. D.H. thanks Henri Niskanen and Alexandre Magalhaes for consultation on experimental design. We thank Philip J. Kranzusch and Samuel J. Hobbs for providing the penguinpox cGAMP PDE target and Celia W. Goulding and Christine D. Hardy for providing the filamentous hemagglutinin target. A.S. and G-W.L. acknowledge members of the Li lab for helpful discussions.

For the compute resources required to complete the project, we thank Novo Nordisk, the National Energy Research Scientific Computing Center (NERSC), a Department of Energy User Facility using NERSC award ERCAP0035823, Anantha P. Chandrakasan, Chris Hill and the Office of Research Computing and Data (ORCD) via a Seed Fund grant, the EuroHPC Joint Undertaking for access to the EuroHPC supercomputer LUMI, hosted by CSC (Finland) and the LUMI consortium through a EuroHPC Regular Access call. We thank The Infrastructure Group (TIG) for managing our MIT computer cluster.

We acknowledge support the Abdul Latif Jameel Clinic from the Machine Learning for Pharmaceutical Discovery and Synthesis (MLPDS) consortium, the MATCHMAKERS project supported by the Cancer Grand Challenges partnership (funded by Cancer Research UK (CGCATF2023/100001), the National Cancer Institute (OT2CA297463) and The Mark Foundation for Cancer Research), the DTRA Discovery of Medical Countermeasures Against New and Emerging (DOMANE) threats program, the NSF Expeditions grant (award 1918839) Understanding the World Through Code, and the DSO Singapore grant on next generation techniques for protein ligand binding.

J.S. acknowledges support by the European Union (CLARA (101136607), ERC FRONTIER (101097822), ELIAS (101120237)) and by the Czech Technology Agency (TA CR) projects RETEMED (TN02000122) and TEREP TN02000122/001N). T.P. acknowledges support by the Czech Science Foundation (GA CR) grant 21-11563M and by the European Union’s Horizon Europe program (ERC, TerpenCode, 101170268). Views and opinions expressed are, however, those of the author(s) only and do not necessarily reflect those of the European Union or the European Research Council. Neither the European Union nor the granting authority can be held responsible for them. This work was also supported by awards K99 GM148718 (to A.S.) and R35 GM124732 (to. G.-W.L.) from the NIH. G.-W.L. is an investigator of the HHMI. W.F.D. acknowledges NIH R35GM122603. L.S. acknowledges NIH K99GM157551. A.K.H acknowledges NIH K99GM155611. E.T.Y. acknowledges NIH F32GM150255. W. D. acknowledges NSF 2306190, NIH 2R35 GM122603, NSF 2448848 Yang Yang. Work in the D.H. lab is supported by the Mark Foundation for Cancer Research. We thank the Institute for Rapid Antibody Engineering and Evolution, part of the Engineering+Health Initiative of the UCI Samueli School of Engineering, for support.

## Supplementary Material

## A Related Work

### Generative Models for Protein Design

After initial proposals for protein structure generative models [Trippe et al., 2022, Anand and Achim, 2022], the first experimental evidence for their efficacy came to light, Watson et al. [2023], Ingraham et al. [2023], causing a range of subsequent works that extensively explored this modeling space [Yim et al., 2023b,a, Bose et al., 2024, Liu et al., 2024, Campbell et al., 2024, Wu et al., 2024, Yim et al., 2024]. Further work has expanded these approaches to all-atom target representations [Krishna et al., 2024, Ren et al., 2025] and enzyme design via atomic-motif scaffolding [Ahern et al., 2025]. The latest generation, including BoltzGen and RFDiffusion3 [Butcher et al., 2025], now enables all-atom binder design against targets of almost any modality. The open-source nature of these methods allows the community to scrutinize the results, validate the underlying assumptions, and draw independent conclusions, which ultimately accelerates progress across the field. Several closed-source models also exist [Chai-Discovery et al., 2025, Team et al., 2025, Bio, 2025].

### All-Atom Protein Structure Generation

Early structure-generative models primarily operated on backbone atoms [Watson et al., 2023], whereas recent approaches perform full all-atom generation [Chu et al., 2024, Qu et al., 2025, Geffner et al., 2025a, Butcher et al., 2025]. A key challenge is that the structure’s number of atoms depends on an unknown sequence. Geffner et al. [2025a] circumvent this by encoding side chains in fixed-size latent vectors, but this prevents explicit atom-level reasoning. Chu et al. [2024] represent all possible side chains simultaneously, yet only update one at each denoising step, limiting joint reasoning over residue alternatives. Qu et al. [2025], Butcher et al. [2025], and BoltzGen instead adopt a padded atom14 representation where each designed residue has 14 atoms. While Qu et al. [2025] and Butcher et al. [2025] require a separate predictor for residue types, BoltzGen encodes residue identity directly in its generated geometry, ensuring explicit residue choices throughout denoising.

### Inverse Folding

Samples from protein structure generative models are typically inverse-folded to obtain sequences compatible with their structures, including in all-atom settings. Most commonly employed methods (but not all [Hsu et al., 2022]) follow the GNN-based formulation of Ingraham et al. [2019], which underpins ProteinMPNN [Dauparas et al., 2022] and its many extensions [Hsu et al., 2022, Goverde et al., 2024, Dauparas et al., 2025]. The BoltzGen inverse folding model is mostly based on these previous works, but is trained on our dataset.

### Hallucination-Based Design

Hallucination approaches have also shown strong performance in design tasks [Anishchenko et al., 2021, Wicky et al., 2022, Pacesa et al., 2024, Cho et al., 2025a,b, Fang et al., 2025]. Unlike generative modeling approaches, these methods directly optimize sequences using gradients from a folding model.

## B Computational Method Details

### B.1 Datasets

#### B.1.1 Training

Our data pipeline builds upon Boltz-2 [Passaro et al., 2025], while adapting the sampling procedure for the task of biomolecular design.

Table 2 summarizes the datasets used for sampling during training, including their sources, sampling cluster types, and associated weights. BoltzGen is primarily trained on entries from the Protein Data Bank (PDB) [Berman et al., 2000] and the AlphaFold Protein Structure Database (AlphaFold DB) [Varadi et al., 2022], while leveraging Boltz-1 distillation [Wohlwend et al., 2025] to enhance performance on underrepresented modalities. For additional information on individual datasets, please see Passaro et al. [2025].

**Table 2:**
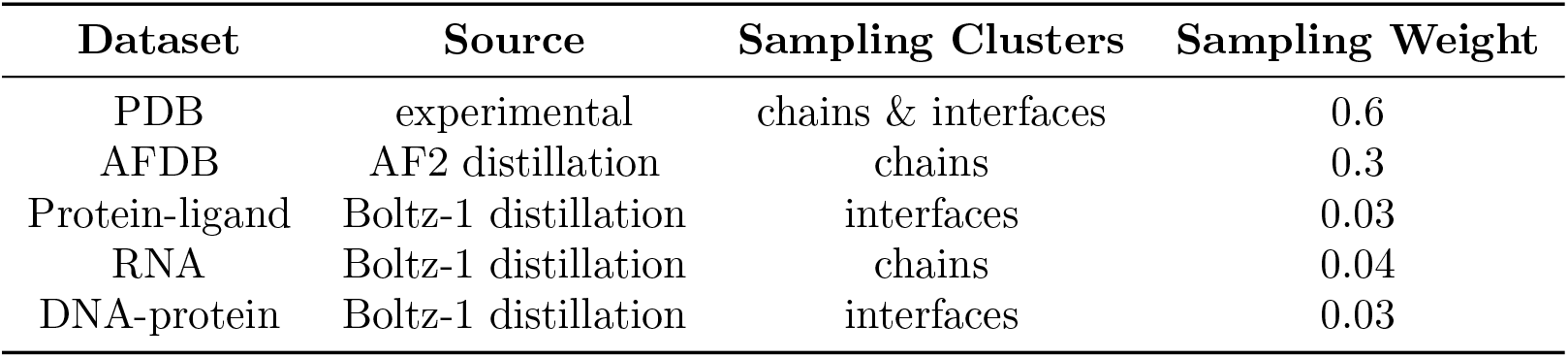
Training data composition.

| Dataset | Source | Sampling Clusters | Sampling Weight |
| --- | --- | --- | --- |
| PDB | experimental | chains & interfaces | 0.6 |
| AFDB | AF2 distillation | chains | 0.3 |
| Protein-ligand | Boltz-1 distillation | interfaces | 0.03 |
| RNA | Boltz-1 distillation | chains | 0.04 |
| DNA-protein | Boltz-1 distillation | interfaces | 0.03 |

##### PDB

We process every structure in the PDB following a pipeline similar to those previously described in Boltz-2 [Passaro et al.,2025]:

- We use every PDB structure up to the training date cutoff of 06/01/2023. We parse the Biological Assembly 1 from these structures.
- For each polymer chain, we use the reference sequence and align it to the residues available in the structure.
- For ligands, we refer to the CCD dictionary to get the reference ligand and atom composition. We compute up to 10 3D conformers per ligand and sample one at random during training.
- We remove large complexes that are over 7MB or with more than 5000 residues.
- We apply the same filters as AlphaFold3, namely excluding crystallization aids and other non-biologically relevant ligands, removing clashing chains, and filtering out chains with fewer than 4 resolved residues or composed only of unknown residues.
- We compute multiple-sequence alignments for every protein chain (and only protein chains) using ColabFold search. Once monomeric MSAs are produced, we assign a taxonomy ID to every sequence in every MSA using their Uniref100 IDs as reference, if any. The preprocessing of the MSAs is analogous to AlphaFold3.
- We produce template hits for protein chains as described in AlphaFold3, using hmmbuild and hmmsearch on PDB sequences deposited at least 60 days prior to any given query’s deposition date.

##### Distilled datasets

We use Boltz-2 distilled datasets, in particular:

- *AlphaFold Database (AFDB) distillation:* In order to construct a protein monomer distillation set, we begin with uniref30 and find the overlap between those sequences and the uniclust multiple sequence alignments provided by OpenFold. We then fetch structures from the AFDB where we impose a minimum global lDDT of 0.5. This procedure results in a monomer distillation of about 5 million proteins.
- *Protein-Ligand distillation*: We construct a dataset of protein-ligand distillation from BindingDB and ChEMBL that were excluded from the main hit-to-lead affinity training set of Boltz-2 [Passaro et al.,2025]. The distillation set was formed by filtering Boltz-1 predictions to examples with a maximum interface predicted distance error (iPDE) ≤ 1.0 and a minimum interface predicted TM-Score (ipTM) ≥ 0.9.
- *RNA distillation*: Following AlphaFold3, we clustered Rfam (v14.9) [Kalvari et al.,2021] using MMSeqs2 [Steinegger and Söding,2017] with 90% identity and 80% coverage. To form the distillation set, Boltz-1 predictions for cluster representatives are filtered to those where the maximum average predicted distance error (PDE) ≤ 2.0.
- *Protein-DNA distillation*: The protein-DNA distillation data is constructed similarly to AlphaFold3. Using the JASPAR 2024 release (specifically, the CORE collection), we first find transcription factor profiles with matching gene IDs across two high-throughput SELEX datasets [Jolma et al.,2015,Yin et al.,2017]. For each filtered profile, a protein sequence is assigned in two ways: i) using the canonical protein sequence under the profile’s Uniprot ID and ii) searching for the sequence in the two SELEX datasets (with matching gene ID) with the highest similarity to the Uniprot sequence. Sequence similarity is calculated using KAlign v2.0, computed as the number of non-gap matches between the two sequences divided by the minimum length of pre-aligned sequences. Unlike AlphaFold3, we did not apply any sequence clustering. To generate binding DNA sequences for each protein sequence, we use the corresponding JASPAR profile’s position frequency matrix (PFM) to sample 10 single-stranded motifs. For each distillation example, the inputs include the protein sequence, the single-strand DNA sequence and its corresponding reverse complement. After generating Boltz-1 predictions, we filtered examples to those that satisfied all the following conditions PDE ≤ 2.0, maximum interface predicted distance error (iPDE) ≤ 1.0 and minimum interface predicted TM-Score (ipTM) ≥ 0.7.

#### B.1.2 Structure Prediction Test Set

For structure prediction, the test set in table 15 was constructed following Boltz-2 [Passaro et al., 2025]:

1. Initial release date is between 2024-01-01 and 2024-12-31.
2. Resolution is below 4.5Å.
3. We select all polymer chains that have less than 40% similarity to training polymer chains.
4. We select all interfaces where at least one of the two chains is dissimilar from the training chains.
5. Given these chains and interfaces, we get all the relevant full targets and always predict assembly 1.

We exclude 187 complexes since they do not fit on 40GB GPUs.

### B.2 Cropping

Each sampled training entry is randomly cropped. While AlphaFold3 [Abramson et al., 2024] alternates between contiguous, spatial, and interface-spatial cropping, we use the single strategy of Boltz-1 [Wohlwend et al., 2025] tailored to biomolecule design. Algorithm 4 shows how training tokens *T* are processed: given a center index *c* determined by the sampling cluster type (Table 2), we iteratively add contiguous fragments of size *W* from chains nearest to the center until reaching the target crop size *L*. The maximum crop size is 768 for folding, matching AlphaFold3 and Boltz-2, and 512 for design tasks to accommodate additional memory for fake atoms.

#### Algorithm 4 CropNeighborhood

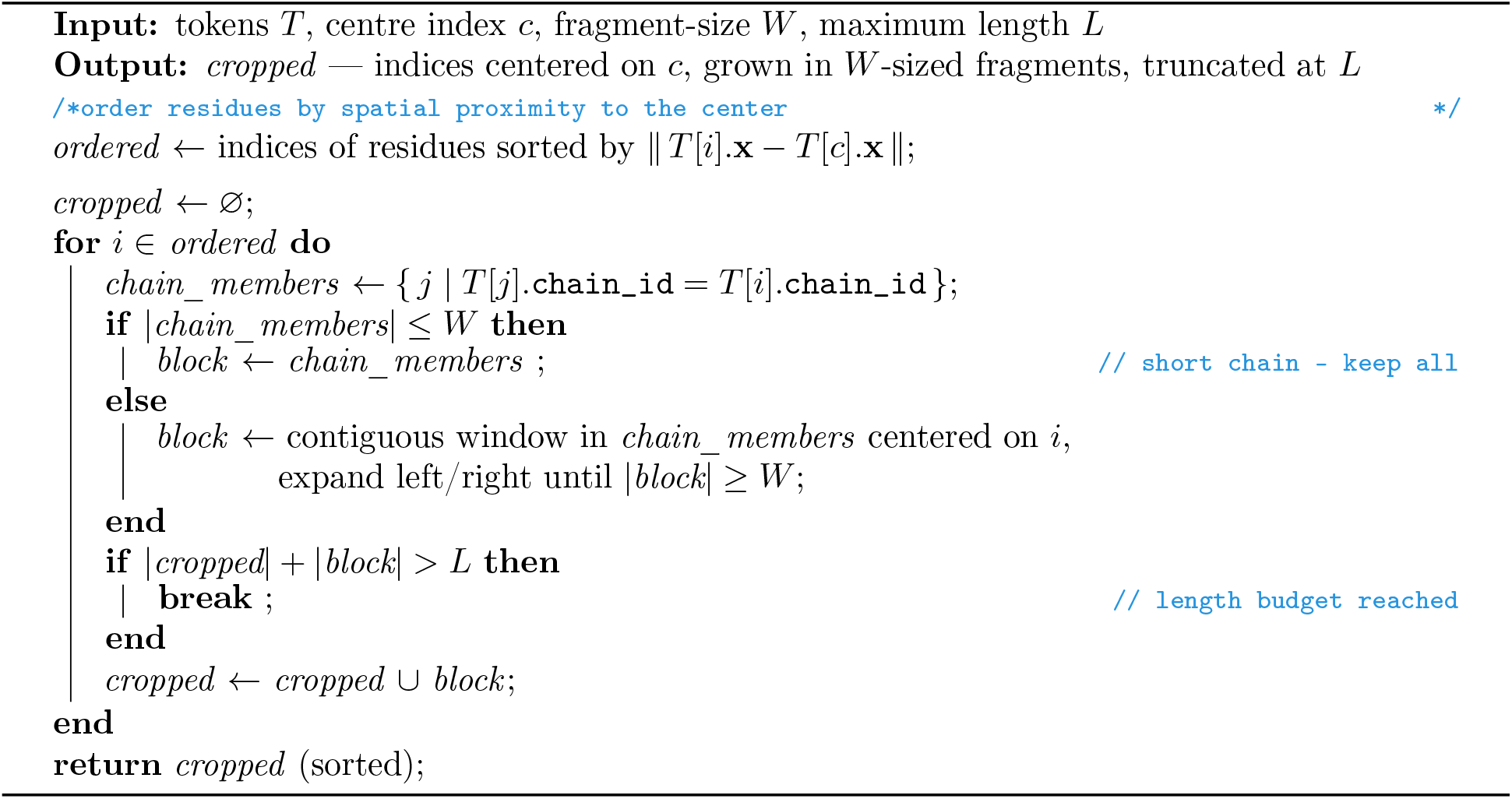

### B.3 Training Tasks

During training, we select different parts of the data sample to be designed. We do this according to different training tasks, which correspond to common use cases such as binder design, or motif scaffolding, as described in Sec. 3.3. The tasks are outlined in Table. 3.

**Table 3:**
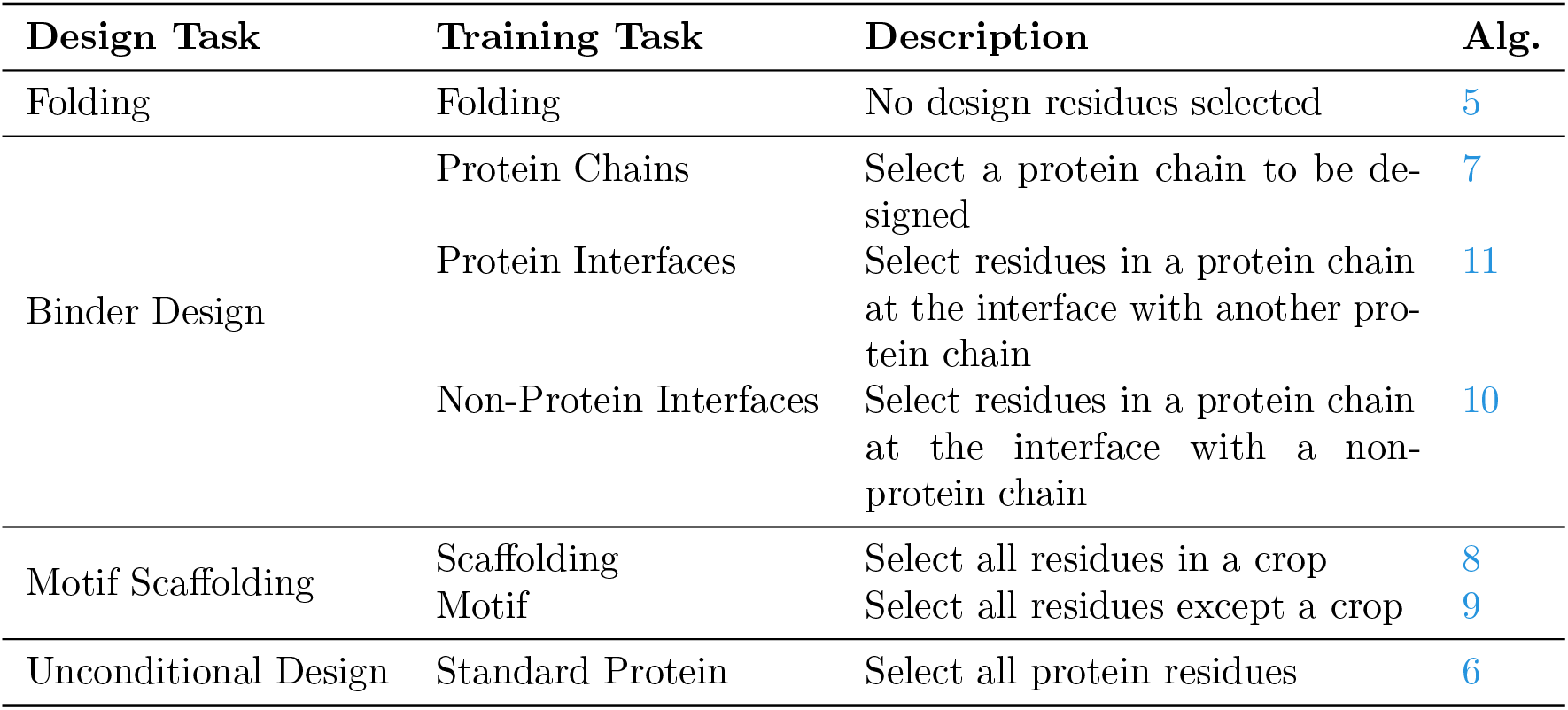
Training Tasks Descriptions.

| Design Task | Training Task | Description | Alg. |
| --- | --- | --- | --- |
| Folding | Folding | No design residues selected | 5 |
| Binder Design | Protein Chains | Select a protein chain to be designed | 7 |
|  | Protein Interfaces | Select residues in a protein chain at the interface with another protein chain | 11 |
|  | Non-Protein Interfaces | Select residues in a protein chain at the interface with a non-protein chain | 10 |
| Motif Scaffolding | Scaffolding | Select all residues in a crop | 8 |
|  | Motif | Select all residues except a crop | 9 |
| Unconditional Design | Standard Protein | Select all protein residues | 6 |

Each task is sampled with a certain probability depending on the data sample. These sampling probabilities are given in Table 4.

**Table 4:** Training Tasks Distribution.

| Task | Condition |  |  |  |  |
| --- | --- | --- | --- | --- | --- |
|  | 0 Non-Protein |  |  | > 0 Non-Protein |  |
|  | 0 Protein | 1 Protein | > 1 Protein | 1 Protein | > 1 Protein |
| Folding (Alg. 5) | 1 | 0.1 | 0.05 | 0.05 | 0.05 |
| Scaffolding (Alg. 9) | 0 | 0.5 | 0.2 | 0.2 | 0.2 |
| Motif (Alg. 8) | 0 | 0.3 | 0.15 | 0.15 | 0.1 |
| Non-Protein Interface (Alg. 10) | 0 | 0 | 0 | 0.2 | 0.05 |
| Standard Protein (Alg. 6) | 0 | 0.1 | 0.1 | 0.4 | 0.1 |
| Protein Interfaces (Alg. 11) | 0 | 0 | 0.1 | 0 | 0.1 |
| Protein Chains (Alg. 7) | 0 | 0 | 0.4 | 0 | 0.4 |

In addition to selecting which residues will be designed, we also sample other conditioning features, such as binding site specifications. The procedures for each feature are listed in Table 5 along with references to algorithmic descriptions.

**Table 5:**
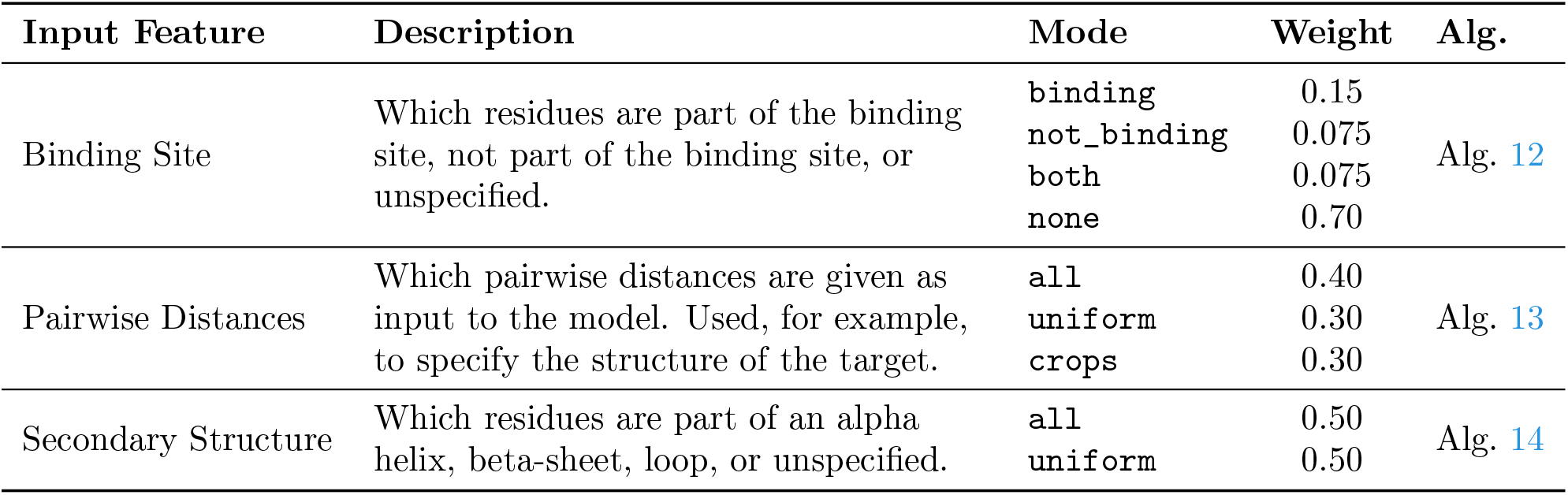
Conditioning Inputs Sampling.

#### Algorithm 5 select_none

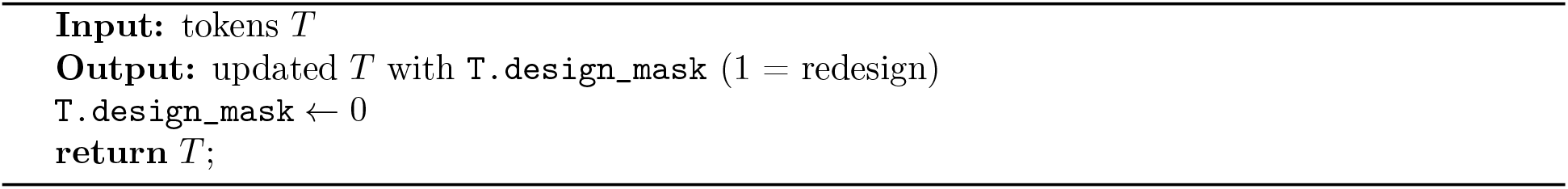

#### Algorithm 6 select_standard_prot

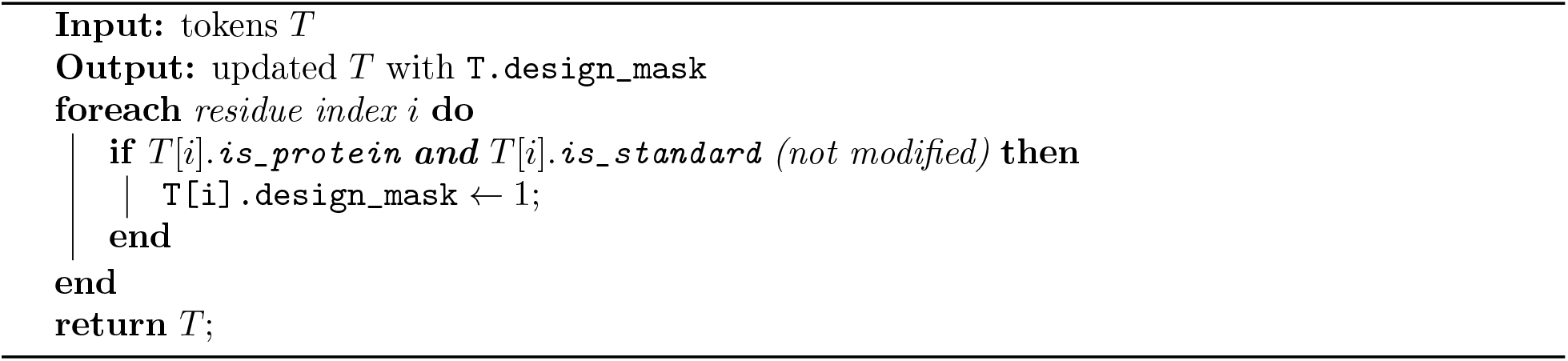

#### Algorithm 7 select_protein_chains

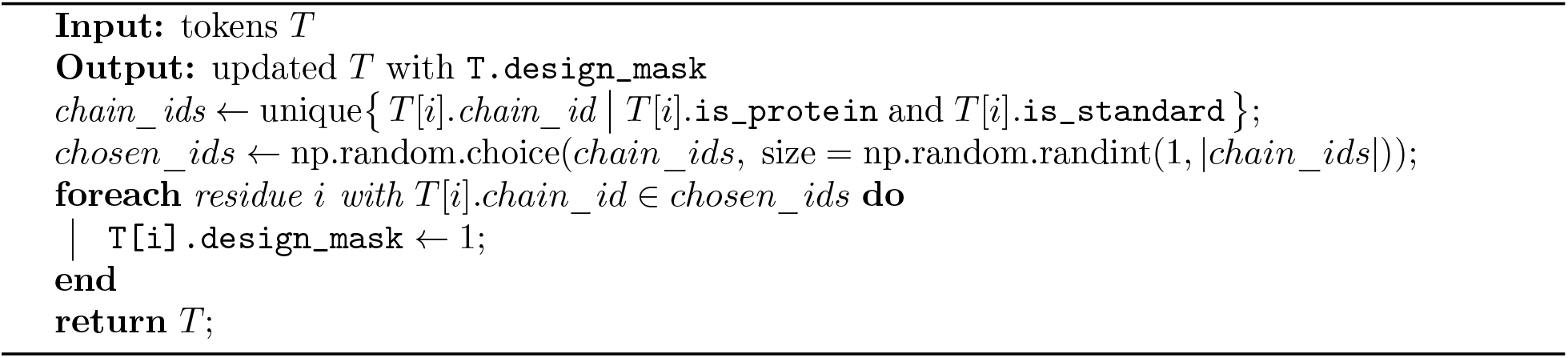

#### Algorithm 8 select_motif

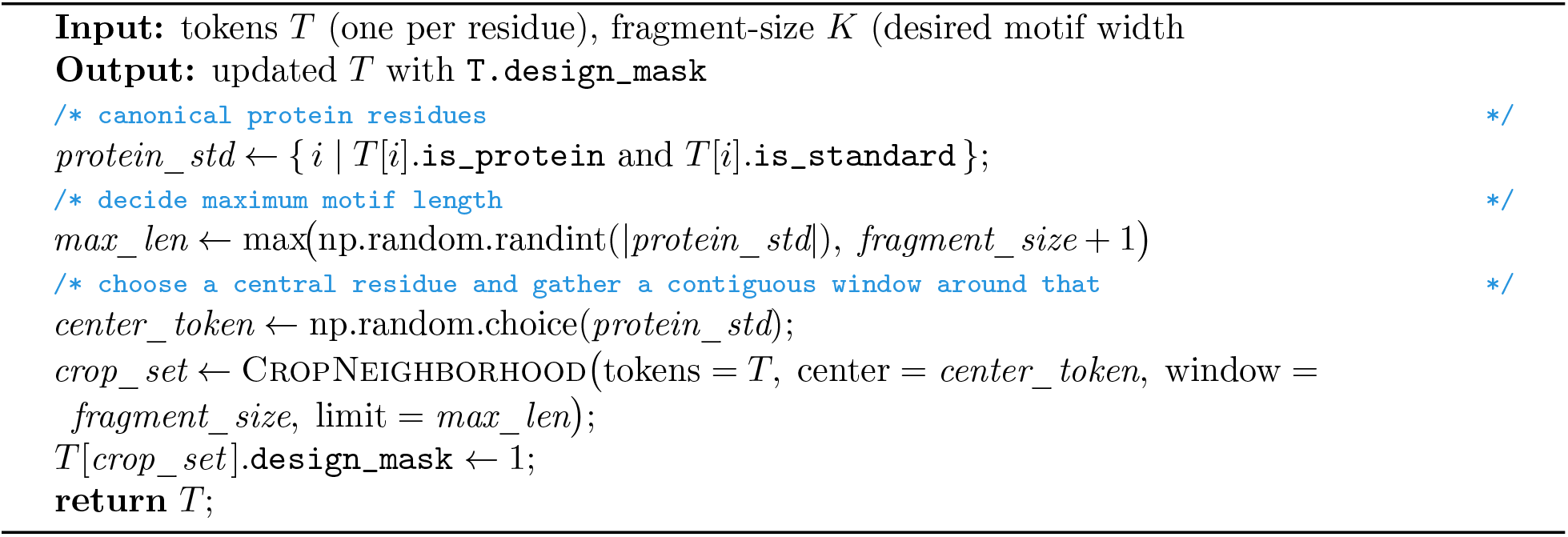

#### Algorithm 9 select_scaffold

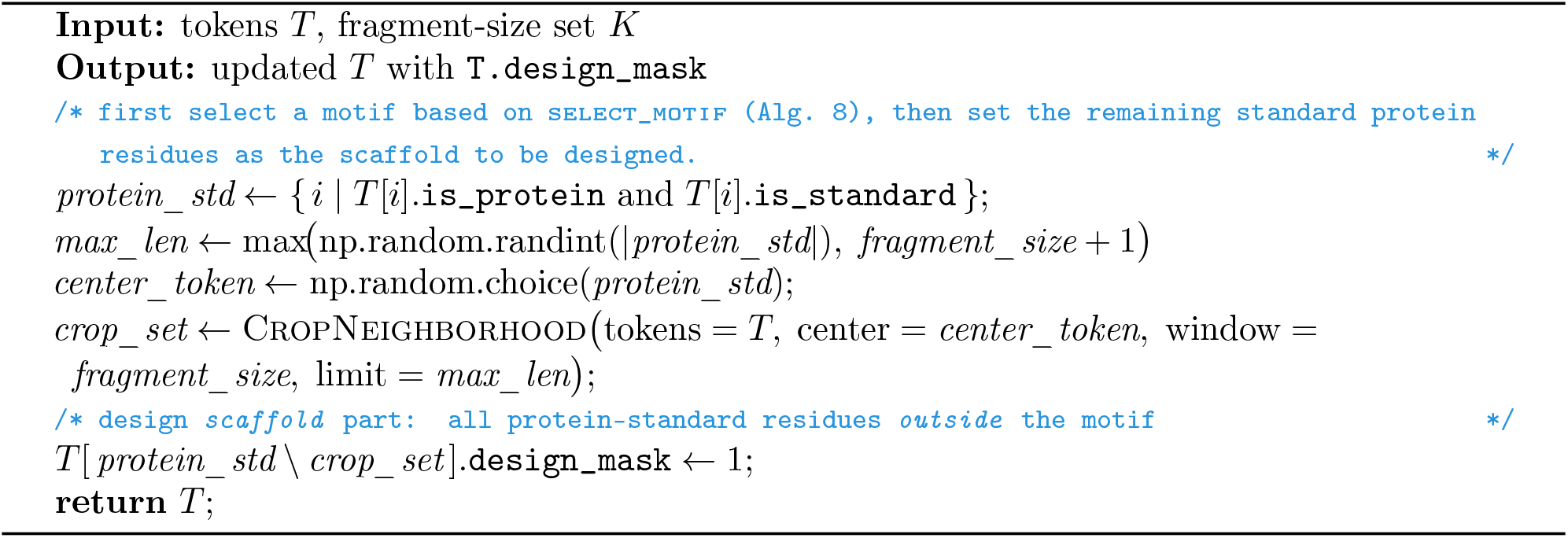

#### Algorithm 10 select_nonprot_interface

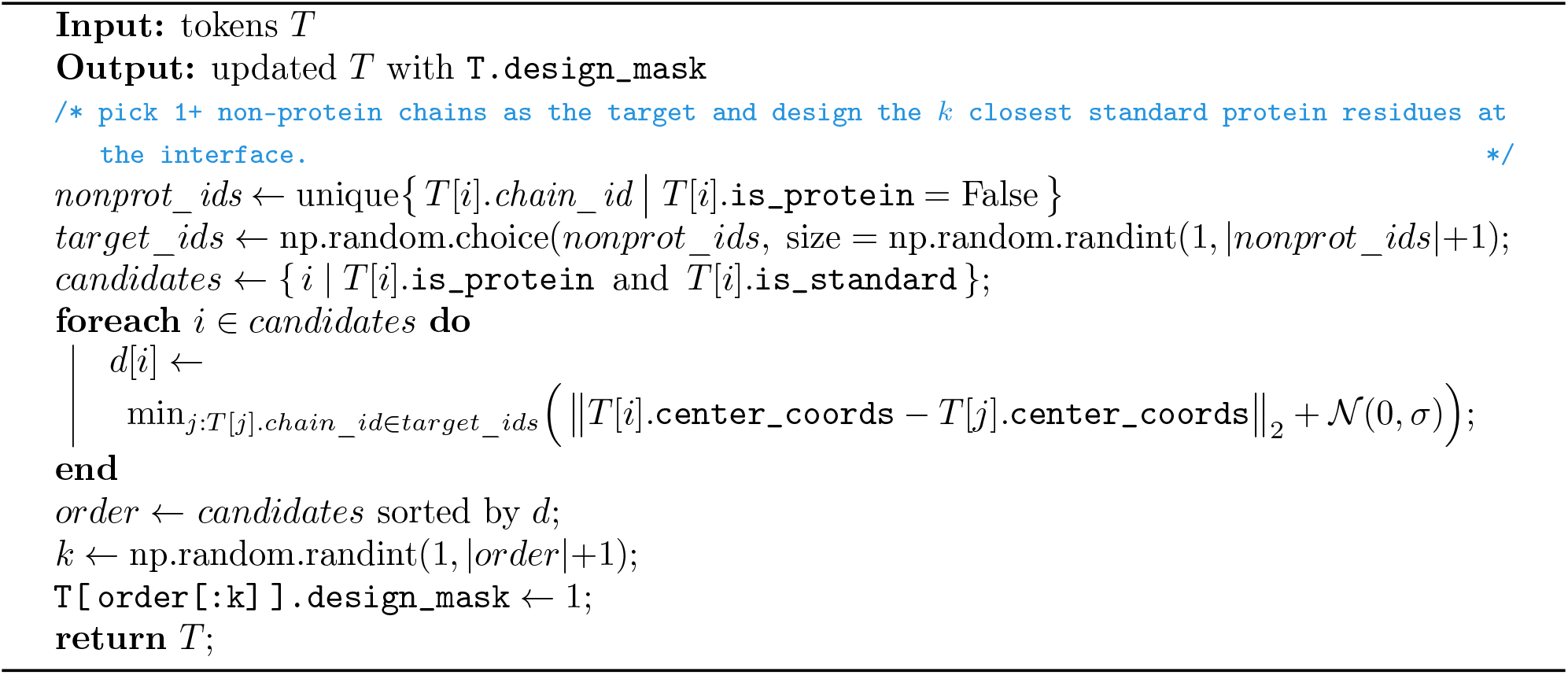

#### Algorithm 11 select_protein_interfaces

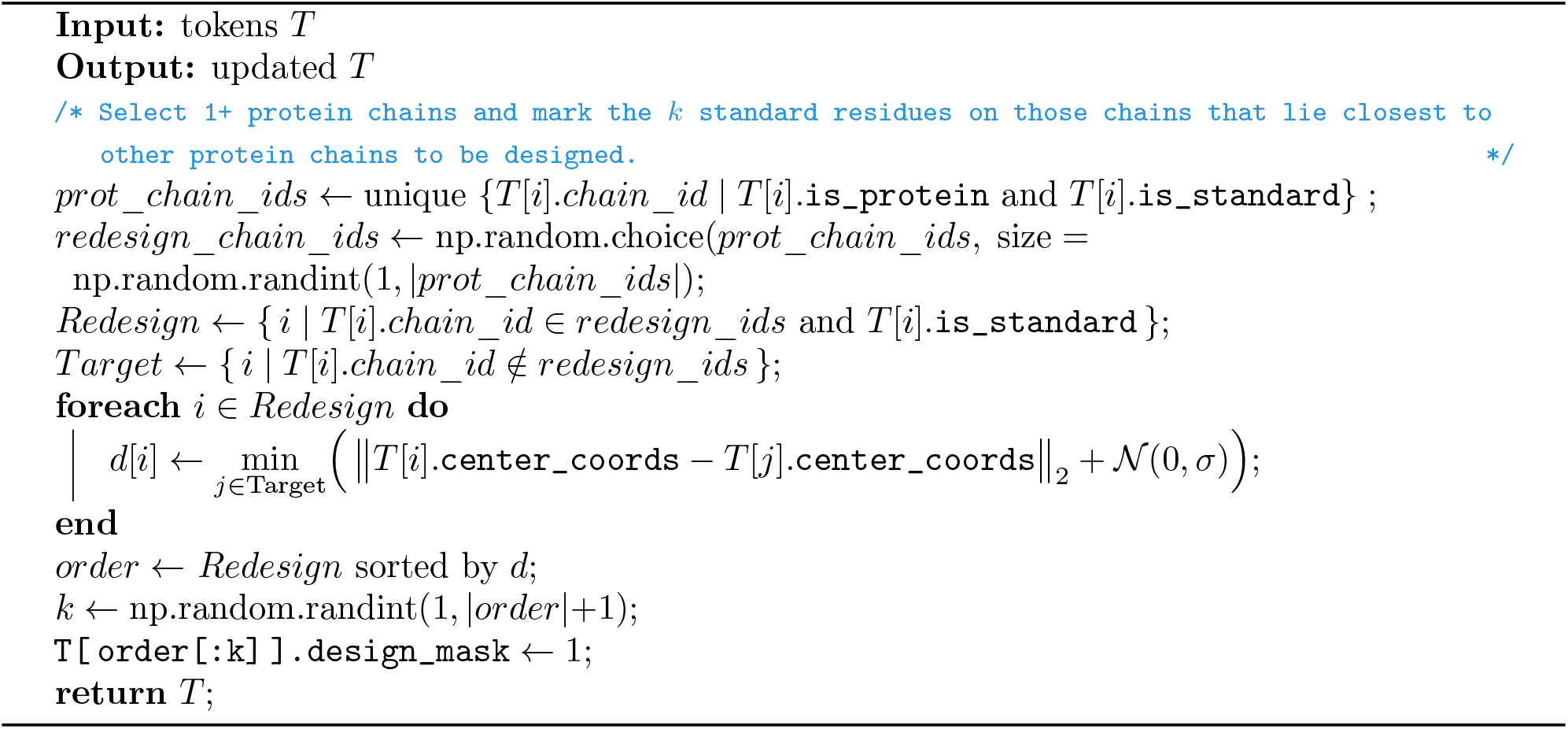

#### Algorithm 12 specify_binding_site

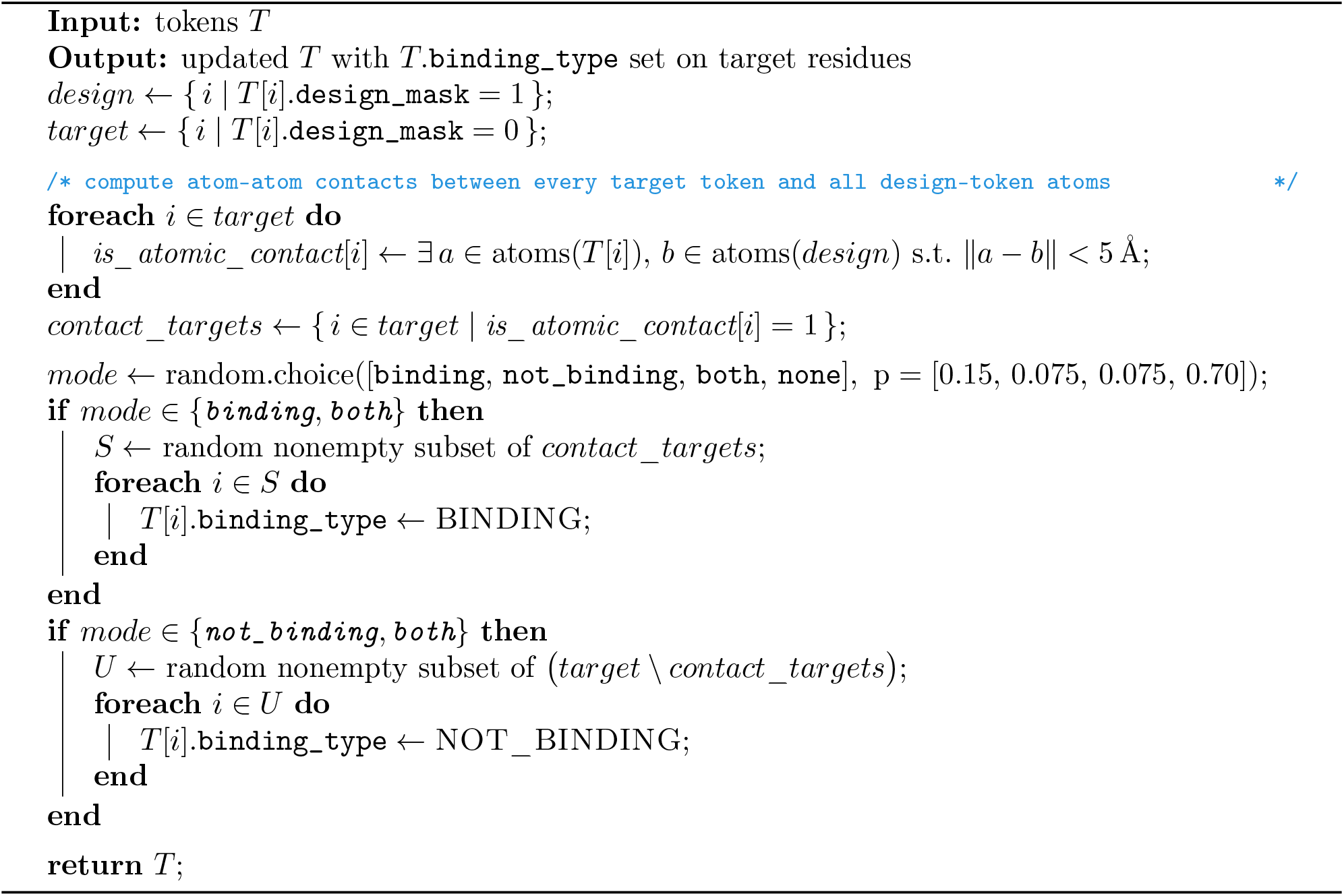

#### Algorithm 13 specify_structure_groups (pairwise–distance conditioning)

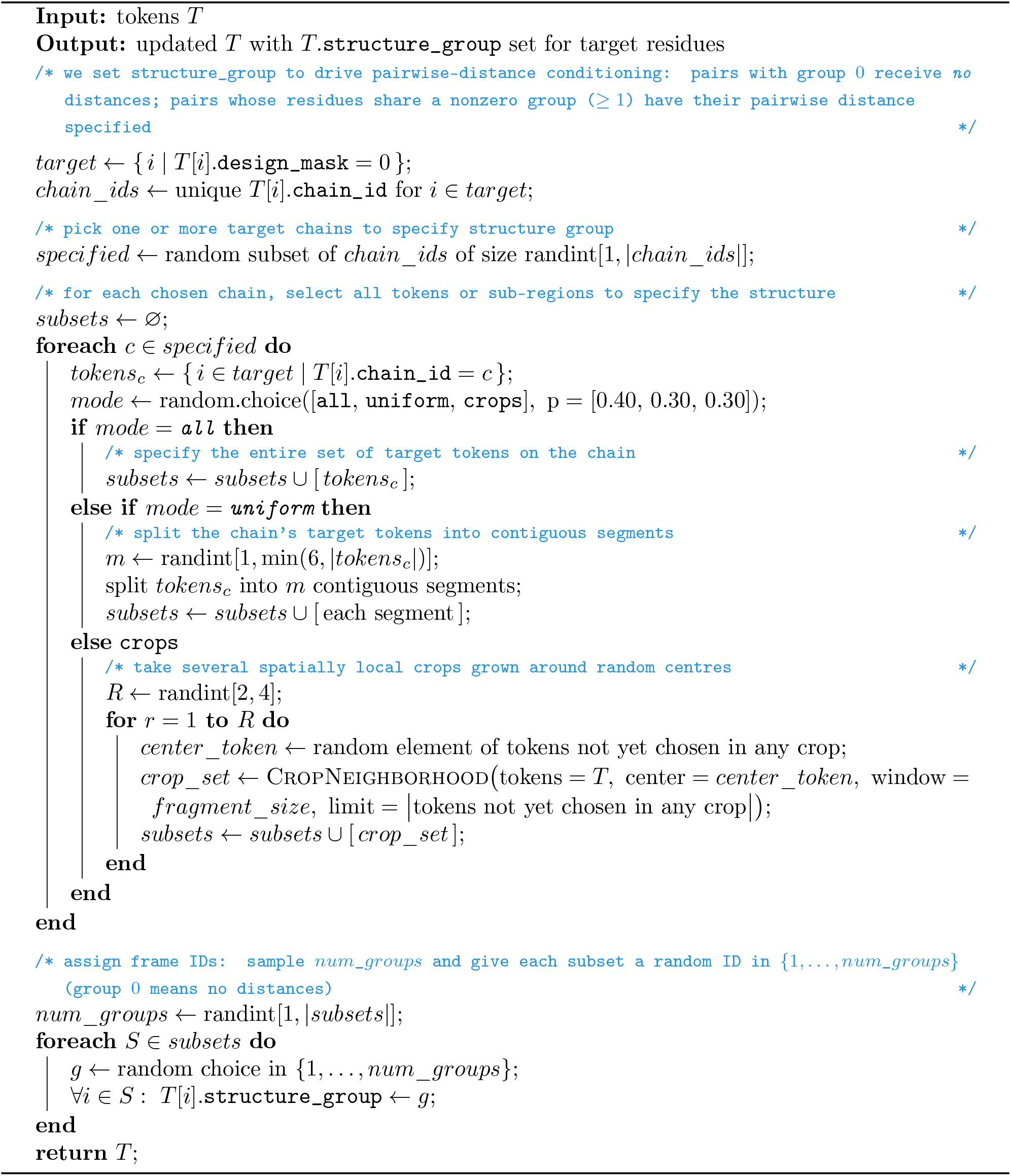

#### Algorithm 14 specify_secondary_structure_mask

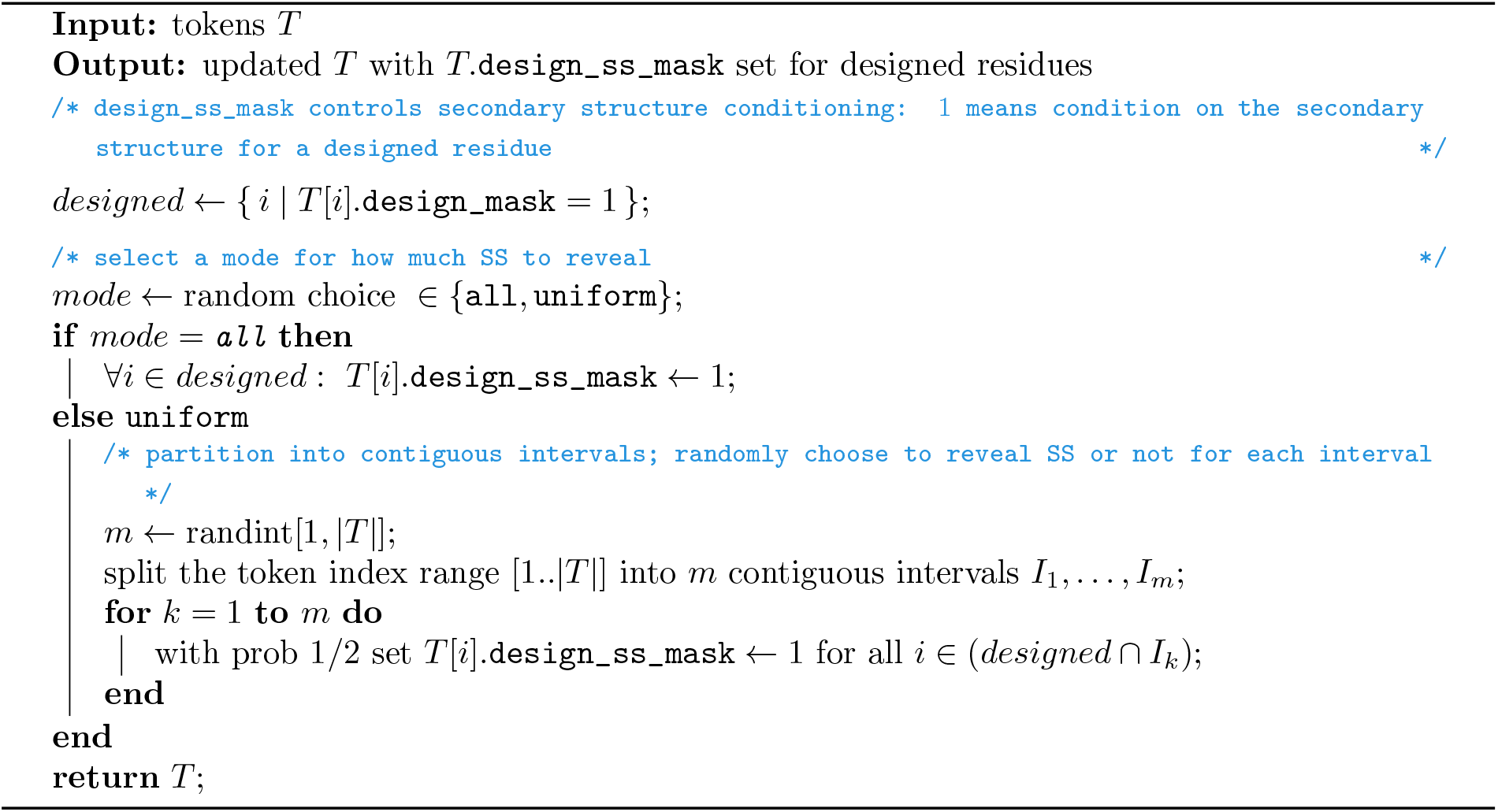

### B.4 Details about Computed Metrics

Table 6 provides a comprehensive reference for all metrics computed by the BoltzGen pipeline.

**Table 6:**
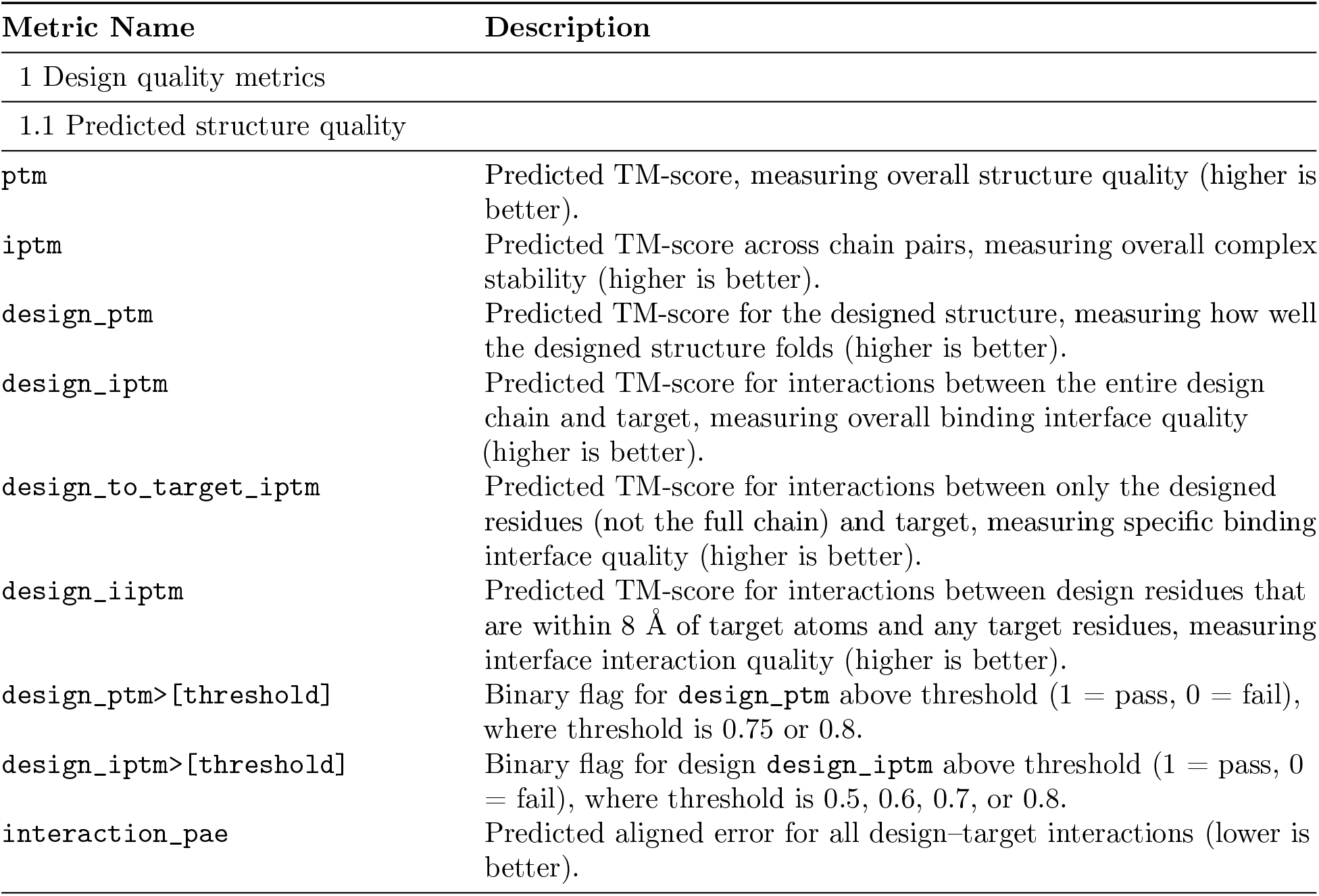
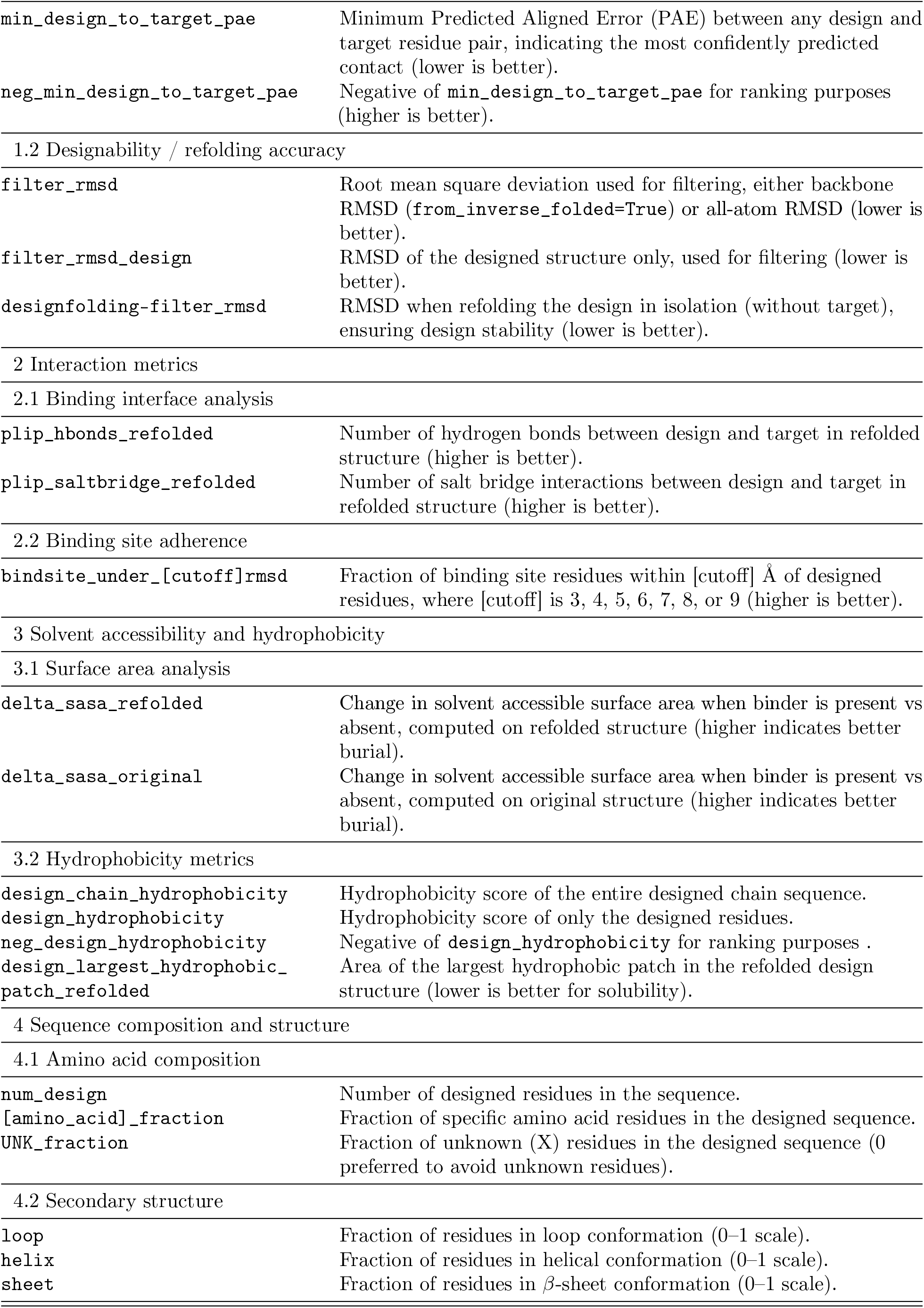
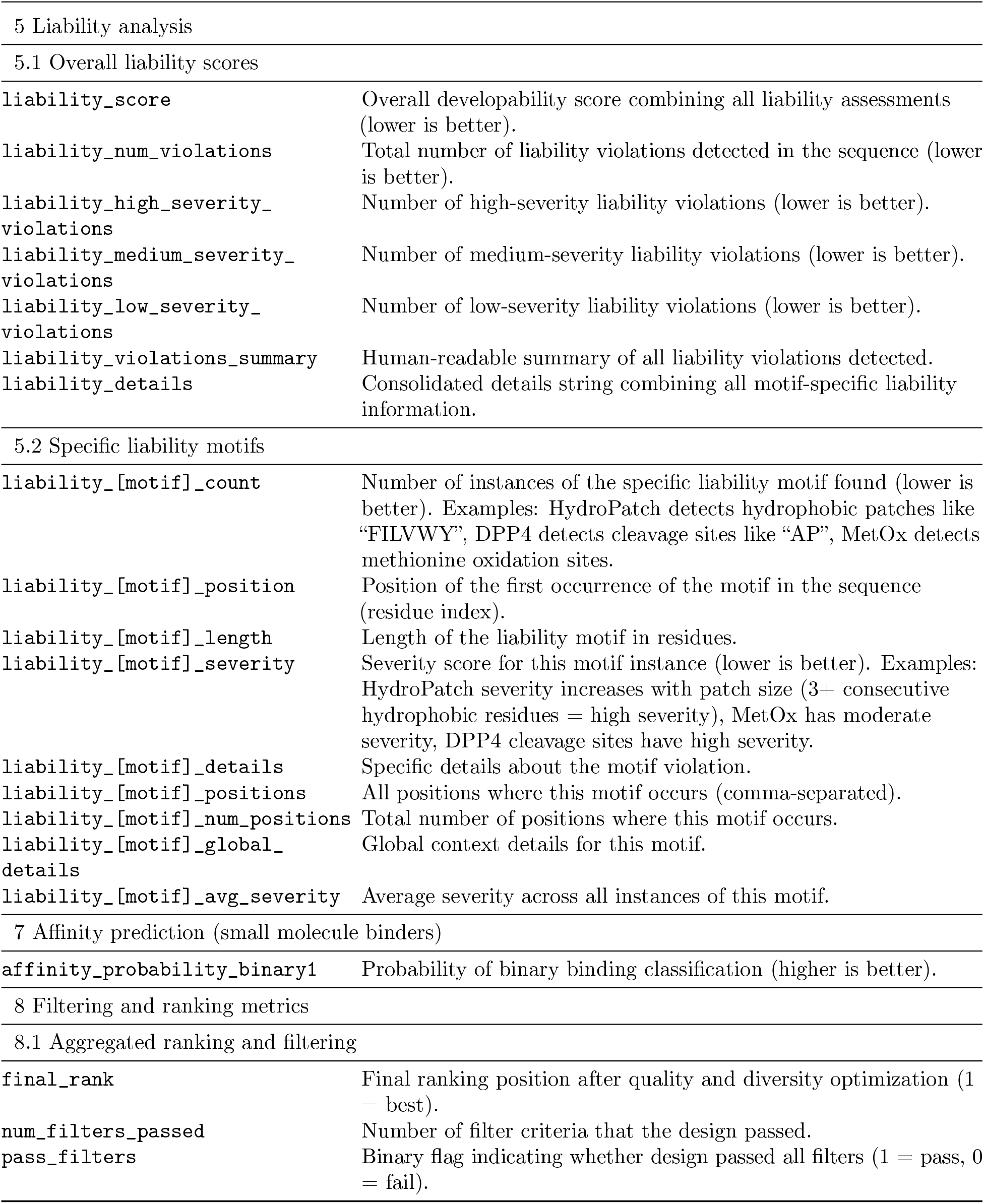
BoltzGen metrics reference.

| Metric Name | Description |
| --- | --- |
| 1 Design quality metrics |  |
| 1.1 Predicted structure quality |  |
| ptm | Predicted TM-score, measuring overall structure quality (higher is better). |
| iptm | Predicted TM-score across chain pairs, measuring overall complex stability (higher is better). |
| design_ptm | Predicted TM-score for the designed structure, measuring how well the designed structure folds (higher is better). |
| design_iptm | Predicted TM-score for interactions between the entire design chain and target, measuring overall binding interface quality (higher is better). |
| design_to_target_iptm | Predicted TM-score for interactions between only the designed residues (not the full chain) and target, measuring specific binding interface quality (higher is better). |
| design_iiptm | Predicted TM-score for interactions between design residues that are within 8 Å of target atoms and any target residues, measuring interface interaction quality (higher is better). |
| design_ptm>[threshold] | Binary flag for **design_ptm** above threshold (1 = pass, 0 = fail), where threshold is 0.75 or 0.8. |
| design_iptm>[threshold] | Binary flag for design **design_iptm** above threshold (1 = pass, 0 = fail), where threshold is 0.5, 0.6, 0.7, or 0.8. |
| interaction_pae | Predicted aligned error for all design–target interactions (lower is better). |
| min_design_to_target_pae | Minimum Predicted Aligned Error (PAE) between any design and target residue pair, indicating the most confidently predicted contact (lower is better). |
| neg_min_design_to_target_pae | Negative of min_design_to_target_pae for ranking purposes (higher is better). |
| 1.2 Designability / refolding accuracy |  |
| filter_rmsd | Root mean square deviation used for filtering, either backbone RMSD (from_inverse_folded=True) or all-atom RMSD (lower is better). |
| filter_rmsd_design | RMSD of the designed structure only, used for filtering (lower is better). |
| designfolding-filter_rmsd | RMSD when refolding the design in isolation (without target), ensuring design stability (lower is better). |
| 2 Interaction metrics |  |
| 2.1 Binding interface analysis |  |
| plip_hbonds_refolded | Number of hydrogen bonds between design and target in refolded structure (higher is better). |
| plip_saltbridge_refolded | Number of salt bridge interactions between design and target in refolded structure (higher is better). |
| 2.2 Binding site adherence |  |
| bindsite_under_[cutoff]rmsd | Fraction of binding site residues within [cutoff] Å of designed residues, where [cutoff] is 3, 4, 5, 6, 7, 8, or 9 (higher is better). |
| 3 Solvent accessibility and hydrophobicity |  |
| 3.1 Surface area analysis |  |
| delta_sasa_refolded | Change in solvent accessible surface area when binder is present vs absent, computed on refolded structure (higher indicates better burial). |
| delta_sasa_original | Change in solvent accessible surface area when binder is present vs absent, computed on original structure (higher indicates better burial). |
| 3.2 Hydrophobicity metrics |  |
| design_chain_hydrophobicity | Hydrophobicity score of the entire designed chain sequence. |
| design_hydrophobicity | Hydrophobicity score of only the designed residues. |
| neg_design_hydrophobicity | Negative of design_hydrophobicity for ranking purposes. |
| design_largest_hydrophobic_patch_refolded | Area of the largest hydrophobic patch in the refolded design structure (lower is better for solubility). |
| 4 Sequence composition and structure |  |
| 4.1 Amino acid composition |  |
| num_design | Number of designed residues in the sequence. |
| [amino_acid]_fraction | Fraction of specific amino acid residues in the designed sequence. |
| UNK_fraction | Fraction of unknown (X) residues in the designed sequence (0 preferred to avoid unknown residues). |
| 4.2 Secondary structure |  |
| loop | Fraction of residues in loop conformation (0–1 scale). |
| helix | Fraction of residues in helical conformation (0–1 scale). |
| sheet | Fraction of residues in β-sheet conformation (0–1 scale). |
| 5 Liability analysis |  |
| 5.1 Overall liability scores |  |
| liability_score | Overall developability score combining all liability assessments (lower is better). |
| liability_num_violations | Total number of liability violations detected in the sequence (lower is better). |
| liability_high_severity_violations | Number of high-severity liability violations (lower is better). |
| liability_medium_severity_violations | Number of medium-severity liability violations (lower is better). |
| liability_low_severity_violations | Number of low-severity liability violations (lower is better). |
| liability_violations_summary | Human-readable summary of all liability violations detected. |
| liability_details | Consolidated details string combining all motif-specific liability information. |
| 5.2 Specific liability motifs |  |
| liability_[motif]_count | Number of instances of the specific liability motif found (lower is better). Examples: HydroPatch detects hydrophobic patches like "FILVWY", DPP4 detects cleavage sites like "AP", MetOx detects methionine oxidation sites. |
| liability_[motif]_position | Position of the first occurrence of the motif in the sequence (residue index). |
| liability_[motif]_length | Length of the liability motif in residues. |
| liability_[motif]_severity | Severity score for this motif instance (lower is better). Examples: HydroPatch severity increases with patch size (3+ consecutive hydrophobic residues = high severity), MetOx has moderate severity, DPP4 cleavage sites have high severity. |
| liability_[motif]_details | Specific details about the motif violation. |
| liability_[motif]_positions | All positions where this motif occurs (comma-separated). |
| liability_[motif]_num_positions | Total number of positions where this motif occurs. |
| liability_[motif]_global_details | Global context details for this motif. |
| liability_[motif]_avg_severity | Average severity across all instances of this motif. |
| 7 Affinity prediction (small molecule binders) |  |
| affinity_probability_binary1 | Probability of binary binding classification (higher is better). |
| 8 Filtering and ranking metrics |  |
| 8.1 Aggregated ranking and filtering |  |
| final_rank | Final ranking position after quality and diversity optimization (1 = best). |
| num_filters_passed | Number of filter criteria that the design passed. |
| pass_filters | Binary flag indicating whether design passed all filters (1 = pass, 0 = fail). |

## C Additional Computational Results and Details

### C.1 Occurences of Benchmark Therapeutic Targets in PDB

1. **TNF***α* There are 18 matching PDB complexes, i.e., entries with more than 1 protein entity and with over 90% sequence identity to the structure used to design binders. See, for example, PDB
2. 5M2I, released in 2017, showing TNF*α* in complex with picomolar-affinity nanobodies [Beirnaert et al., 2017].
3. **PD-L1** There are 37 matching PDB complexes. See, for example, PDB 5JDS, released in 2017, showing PD-L1 in complex with a 3.0-nanomolar-affinity nanobody [Zhang et al., 2017].
4. **PDGFR** The PDB entry 3MJG used for making designs, released in 2010, shows PDGFR in complex with its binding partner PDGF, and previous work has reported *de novo* mini protein binders with nanomolar affinities [Cao et al., 2022].
5. **IL-7R***α* There are 6 matching PDB complexes, such as PDB entry 6P50, released in 2019, which shows IL-7R*α* in complex with a 1-nanomolar-affinity Fab [Hixon et al., 2020].
6. **InsulinR** There are 74 matching PDB complexes. The PDB entry 4ZXB used for design, released in 2016, shows the target in complex with four Fabs (83-7 and 83-14) [Croll et al., 2016].

### C.2 Calibrating Filtering Algorithm for Protein-Protein Complexes

The relative importance of the Boltz-2 confidence metrics and interaction metrics used to rank designs is calibrated on a benchmark of 11,000 validated binders across 11 target proteins, based on data from Cao et al. [2022]. These weights serve as default values and are manually adjusted for wetlab design experiments based on domain expert feedback.

#### Binder designs selection benchmark

For each of the 11 targets (InsulinR, FGFR2, EGFR, H3, IL7Ra, PDGFR, SARS-CoV-2 RBD, TGFb, Tie2, TrkA, and VirB8), we sample up to 100 positive examples (i.e., 4*µ*M binders) and fill the remainder up to 1,000 designs with negative examples (i.e., non-binders), resulting in a balanced subset of 11,000 designs.

When exploring different methods to prioritize designs, we optimize the mean enrichment factor at top 25 and top 50 designs. Following Zambaldi et al. [2024], due to the high variance of metric values across targets, we do not directly optimize the mean enrichment score. Instead, we optimize the mean rank across targets, where each target’s rank is based on its individual enrichment value. When computing enrichment factors, we normalize by the original binder-to-non-binder ratio from the full dataset, rather than our 11,000-design subset.

#### Binder designs selection method

In our initial experiments, we trained a decision tree to predict binary binder labels based on the given metric values. However, we find that learning decision thresholds for individual metrics is not optimal, as the best threshold values can be specific to the target protein (for example, optimal interaction count metric thresholds can vary depending on protein size). Instead of using absolute thresholds, we develop a design selection scheme that prioritizes candidates with the best worst-case ranks across all metrics (Algorithm 2). Each metric is assigned a single learnable weight from the set {0, 1, 1.2, 1.5, 2}, where 0 means that the metric is not used. We calibrate these weights by maximizing the enrichment factor on our benchmark of 11,000 experimentally validated binder designs. We do not run this optimization over all metrics but only over a representative subset of non-correlated ones (Figure 17). We include both `design_iiptm` and `neg_min_design_to_target_pae` despite their high correlation, as variations of these two metrics have been shown to be complementary [Zambaldi et al., 2024].

**Figure 17:**
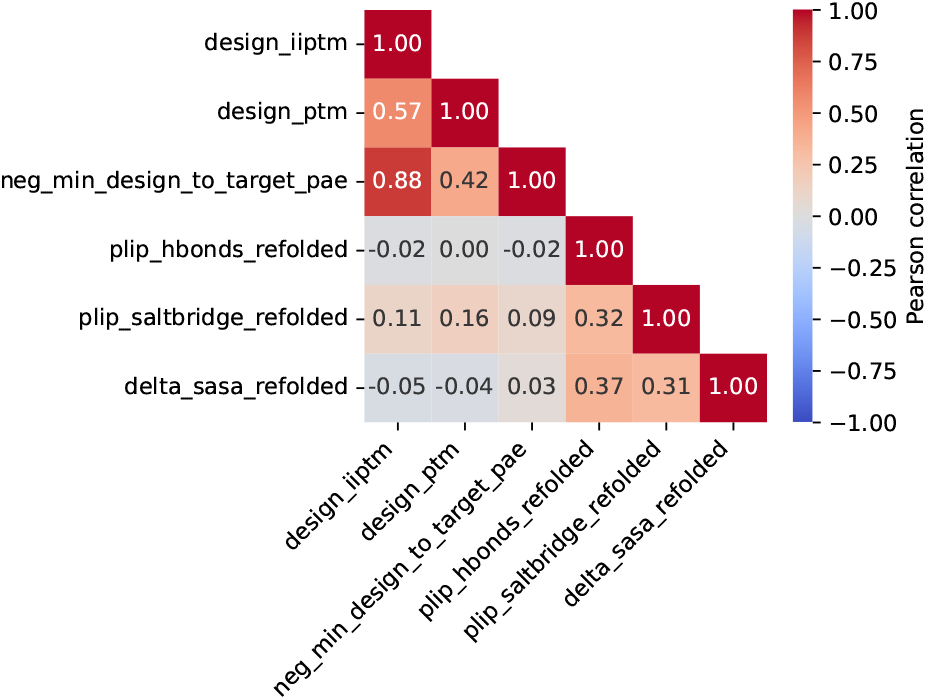
Final Metrics Used For Binder Design Filtering And Their Pairwise Correlations. Correlations were calculated on 11,000 BoltzGen-designed binders for 11 targets from our benchmark derived from Cao et al. [2022].

Our enrichment factor optimization, combined with wetlab experimental feedback, yields the following final combination of metrics and weights for Algorithm 2: `design_iiptm`: 1, `design_ptm`: 2, `neg_min_design_to_target_pae`: 1, `plip_hbonds_refolded`: 2, `plip_saltbridge_refolded`: 2, and `delta_sasa_refolded`: 2. When designing small-molecule binders, we slightly modify the weights of Boltz-2 metrics: `design_iiptm`: 1.1, `design_ptm`: 1.1, `neg_min_design_to_target_pae`: 1.1, `plip_hbonds_refolded`: 2, `plip_saltbridge_refolded`: 2, and `delta_sasa_refolded`: 2.

### C.3 Baseline methods

#### RFdiffusion

We employ RFdiffusion [Watson et al., 2023] as a baseline for binder generation, using the official implementation (https://github.com/RosettaCommons/RFdiffusion). We apply the standard settings (`diffuser.T=100`) and reduce the inference noise to improve design quality (`denoiser.noise_scale_ca=0`, `denoiser.noise_scale_frame=0`), following the configuration used in the official binder design example (https://github.com/RosettaCommons/RFdiffusion/blob/main/examples/design_ppi.sh).

We use ProteinMPNN [Dauparas et al., 2022] for inverse folding, as implemented in the official LigandMPNN [Dauparas et al., 2025] repository (https://github.com/dauparas/LigandMPNN). We use the standard checkpoint `proteinmpnn_v_48_020.pt`. Side-chain packing is performed with the following settings: `pack_side_chains=1`, `number_of_packs_per_design=1`, and `pack_with_ligand_ context=1`.

#### RFdiffusionAA

We use RFdiffusion All-Atom (RFdiffusionAA) [Krishna et al., 2024] as a baseline for generating protein binders against small molecules. We employ the official implementation (https://github.com/baker-laboratory/rf_diffusion_all_atom) with a standard configuration (`diffuser.T=150`, `inference.ckpt_path=RFDiffusionAA_paper_weights.pt`). Inverse folding is performed using LigandMPNN [Dauparas et al., 2025], as described above for RFdiffusion.

BoltzGen only requires a SMILES representation of the input small molecule and performs cofolding during the design process. In contrast, RFdiffusionAA uses a fixed ligand structure to generate a protein binder. To ensure a fair comparison between the two methods, we generate ligand conformers for RFdiffusionAA using RDKit [Bento et al., 2020]. Specifically, inspired by DiffDock [Corso et al., 2022], we employ the `AllChem.ETKDGv3` algorithm and, if it fails, fall back to initializing random coordinates followed by optimization with `AllChem.MMFFOptimizeMolecule`. We generate a single conformer per input ligand, as we observe that increasing the number of conformers to diversify designs has no significant impact on RFdiffusionAA performance.

### C.4 BoltzIF Inverse Folding Model

We verify whether BoltzIF behaves similarly to Protein MPNN (PMPNN) and Soluble (SMPNN) on a set of 64 monomer targets from the PDB. We evaluate each model’s ability to inverse fold both native and designed structures. For native ability, we inverse fold the targets themselves and evaluate 50 sequences for each one. For designed ability, we generate 50 binders for each target with BoltzGen and evaluate 1 inverse-folded sequence per design.

Table 7 reports both the backbone designability of the refolded sequences to the original structures as well as the size of the largest hydrophobic patch, which is relevant for protein expressibility. We see that BoltzIF attains the same designabilities as ProteinMPNN and SolubleMPNN and its hydrophobicity scores fall between the two models.

**Table 7:** Inverse Folding Model Comparison. “RMSD<2.5” denotes the success rate with which designed sequences refold into the inverse folded structure (using Boltz-2). “Hydrophobic” indicates the surface area of the inverse folded protein’s largest hydrophobic patch.

| Method | Native Backbones |  | Designed Backbones |  |
| --- | --- | --- | --- | --- |
|  | RMSD < 2.5 | Hydrophobic | RMSD < 2.5 | Hydrophobic |
| ProteinMPNN | 0.55 | 1343.30 | 0.31 | 1448.65 |
| SolubleMPNN | 0.55 | 1025.72 | 0.33 | 1143.31 |
| BoltzIF | 0.55 | 1185.98 | 0.32 | 1400.73 |

**Table 8:** Number of unique successes on the RFDiffusion benchmark for BoltzGen and 4 other methods, for 1000 backbones.

| Task Name | BoltzGen | Proteina | Genie2 | RFDiffusion | FrameFlow |
| --- | --- | --- | --- | --- | --- |
| 6E6R_short | **83** | 56 | 26 | 23 | 25 |
| 5TRV_med | **25** | 22 | 23 | 10 | 21 |
| 5YUI | **36** | 5 | 3 | 1 | 1 |
| 6EXZ_short | **11** | 3 | 2 | 1 | 3 |
| 5TRV_short | **6** | 1 | 3 | 1 | 1 |
| 4JHW | **3** | 0 | 0 | 0 | 0 |
| 5IUS | **3** | 1 | 1 | 1 | 0 |
| 1PRW | **2** | 1 | 1 | 1 | 1 |
| 6E6R_long | 289 | **713** | 415 | 381 | 110 |
| 6EXZ_long | 59 | 290 | 326 | 167 | **403** |
| 6E6R_medium | 164 | **417** | 272 | 151 | 99 |
| 1YCR | 102 | **249** | 134 | 7 | 149 |
| 5TRV_long | 155 | **179** | 97 | 23 | 77 |
| 4ZYP | 1 | **11** | 3 | 6 | 4 |
| 6EXZ_med | 32 | 43 | 54 | 25 | **110** |
| 7MRX_128 | 64 | 51 | 27 | **66** | 35 |
| 7MRX_85 | 22 | **31** | 23 | 13 | 22 |
| 3IXT | 10 | 8 | **14** | 3 | 8 |
| 5TPN | 3 | 4 | **8** | 5 | 6 |
| 7MRX_60 | 4 | 2 | **5** | 1 | 1 |
| 1QJG | 0 | 3 | 5 | 1 | **18** |
| 1BCF | 1 | 1 | 1 | 1 | 1 |
| 5WN9 | 2 | 2 | 1 | 0 | **3** |
| 2KL8 | 1 | 1 | 1 | 1 | 1 |

**Table 9:** Maximum sequence identity. (against non-monomeric PDB chains) for the selected targets.

| Target | Max sequence identity (%) |
| --- | --- |
| 1G13 | 25.4 |
| 1JQD | 0.0 |
| 1NB0 | 24.6 |
| 2A1X | 0.0 |
| 2PNY | 23.0 |
| 3APU | 19.8 |
| 3CH4 | 22.7 |
| 3QKG | 27.9 |
| 6M1U | 22.5 |
| 7AAH | 24.3 |

**Table 10:**
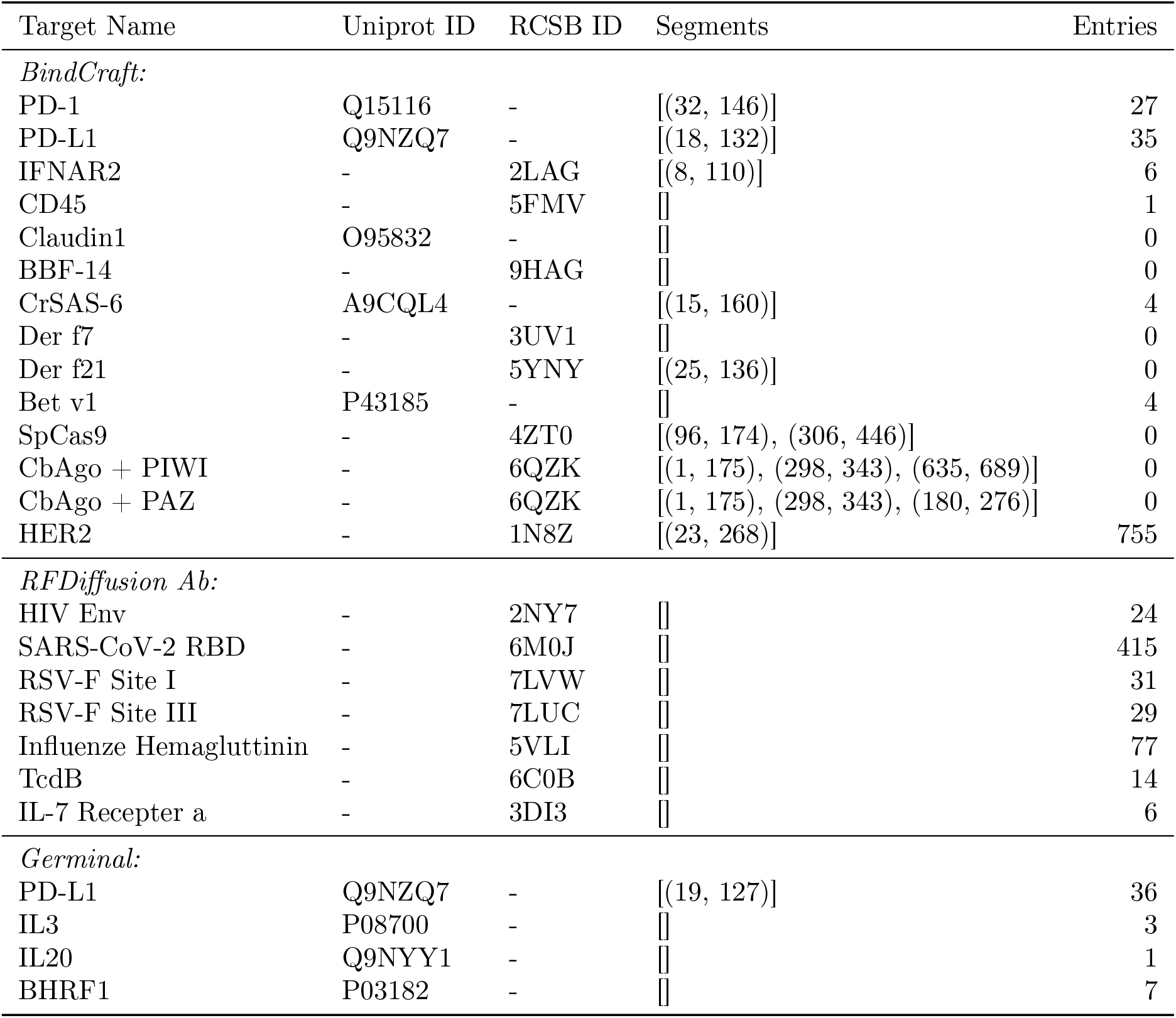
Number of RCSB Complexes Containing Targets From Previous Work. For each target we either use the sequence or subsequences from the Uniprot entry or an RCSB entry, according to the original authors’ descriptions. If no segments are specified we used the full sequence. We report the number of entries in RCSB with at least 90% sequence identity, and at least 2 protein entities. For RFDiffusion Ab, we additionally filter to entries deposited before their training data cutoff of August 2nd 2021.

**Table 11:** Selected binder design targets.

| Target | Assay valency | Material (catalog) | Oligomeric state | Tag | UniProt |
| --- | --- | --- | --- | --- | --- |
| TNFα | Trivalent | Acro · TNA-H4211 | Homotrimer | Untagged | P01375 |
| HNMT | Monovalent | Sino · 13096-H09E | Monomer | GST | P50135 |
| PD-L1 | Monovalent | Acro · PD1-H5229 | Monomer | His | Q9NZQ7 |
| ORM2 | Monovalent | Sino · 13299-H08H | Monomer | His | P19652 |
| Insulin Receptor | Bivalent | Acro · INR-H52Ha | Heterotetramer | His | P06213 |
| RFK | Monovalent | Sino · 13881-H07E | Monomer | His | Q969G6 |
| IL-7RA | Monovalent | Acro · IL7-H52H7 | Monomer | His | P16871 |
| MZB1 | Bivalent | Sino · 15682-H02H | Monomer | Fc | Q8WU39 |
| GM2A | Monovalent | Sino · 13246-H08B | Monomer | His | P17900 |
| IDI2 | Monovalent | Sino · 15292-H07E | Monomer | His | Q9BXS1 |
| AMBP | Bivalent | Sino · 13141-H05H1 | Homodimer | Fc | P02760 |
| PMVK | Monovalent | Sino · 14583-H07E | Monomer | His | Q15126 |
| PDGFR Beta | Bivalent | Acro · PDB-H5259 | Homodimer | Fc | P09619 |
| METTL16 | Monovalent | Sino · M336-380G | Monomer | GST | Q86W50 |
| PHYH | Monovalent | Sino · 13368-HNAE | Monomer | Untagged | O14832 |

**Table 12:**
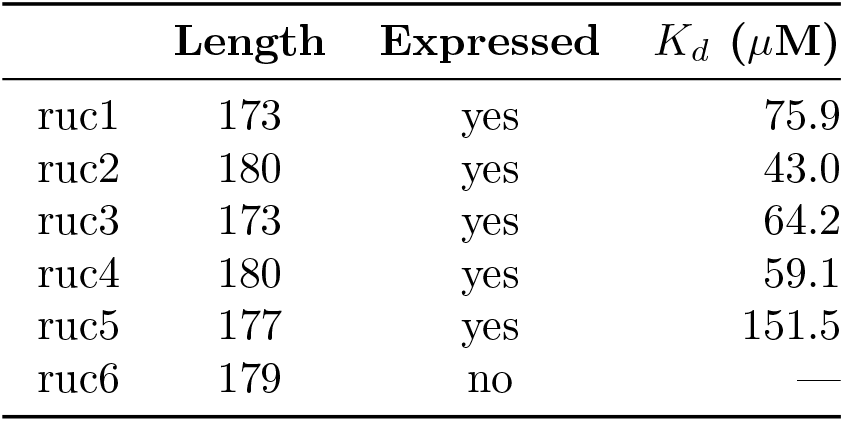
Small Molecule Binders. Experimental characterization of designed binders against rucaparib.

Fig. 18 shows the amino acid distribution over all sequences from each of the models. BoltzIF’s residue frequencies mostly fall between that of ProteinMPNN and SolubleMPNN. Likely, the reason for the higher hydrophobicity scores than SolubleMPNN is that it has been trained on crops (hydrophobic cores can be viewed as solvent-facing when it is the surface of a crop).

**Figure 18:**
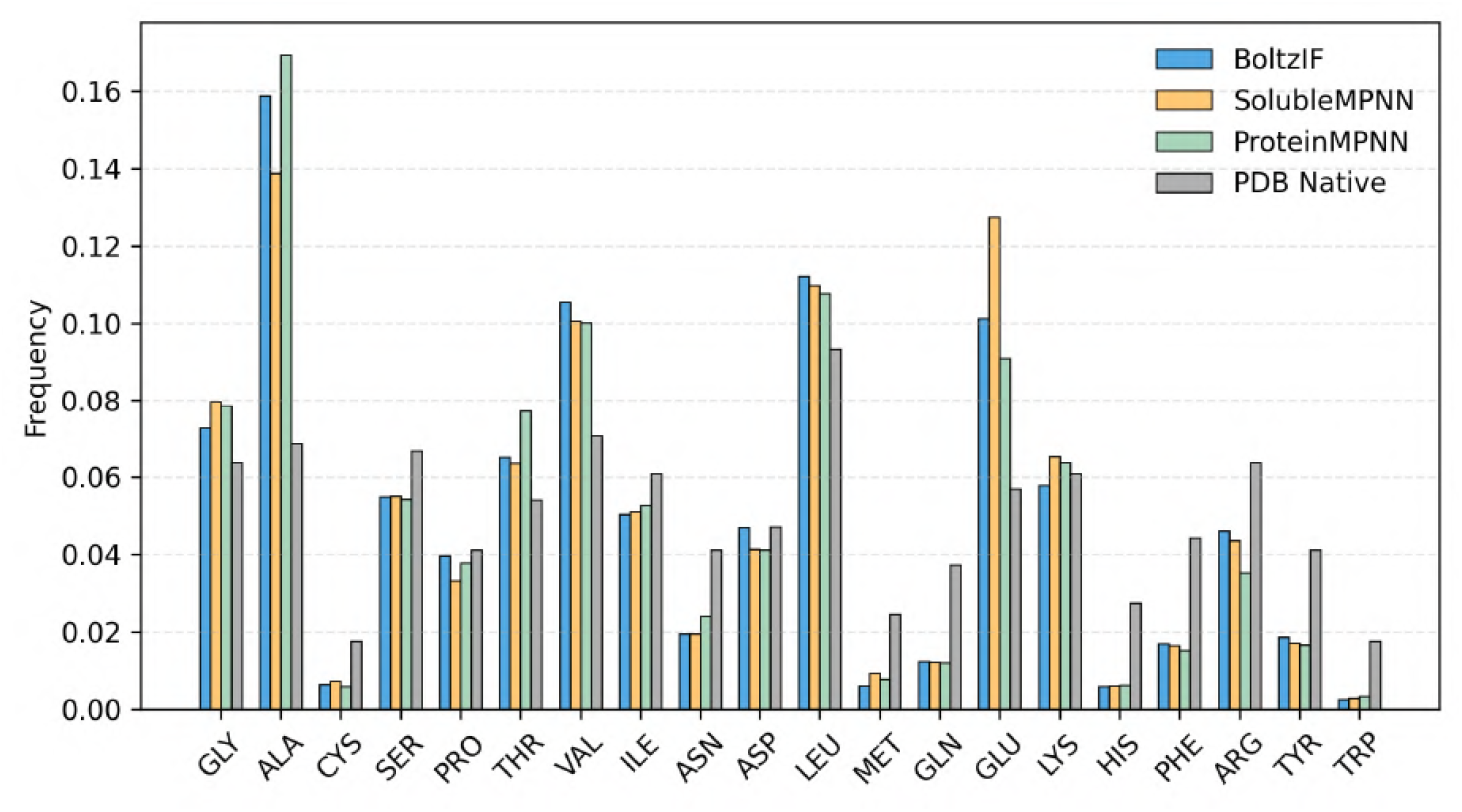
Amino acid distributions when inverse folding BoltzGen’s designed binders against 64 monomers in PDB. “PDB Native” denotes the amino acid distribution in our PDB training data.

### C.5 Motif Scaffolding Benchmark Results

We ran the following Motif-scaffolding performance benchmark performed in [Geffner et al., 2025b]:

- For each motif scaffolding task, we generate 1000 backbones.
- For each backbone, 8 sequences are generated by ProteinMPNN with fixed sequences in the motif region.
- All 8 sequences are refolded via ESMfold and the *C*_*α*_-RMSD and the motifRMSD are computed between the ground truth and the prediction.
- A backbone is categorized as a success when one of the ProteinMPNN sequences satisfy *C*_*α*_-RMSD ≤ 2Å, motifRMSD ≤ 1Å, pLDDT ≥ 70, and pAE ≤ 5.
- Hierarchical clustering with single linkage and TM-score threshold 0.6 is performed on all successful backbones to get a clustering to get the final unique successes.

BoltzGen has the highest number of sole best method in 8 tasks (compared to Proteína that wins in 6 tasks).

### C.6 Memorization of Ubiquitin

In a few design campaigns, we observed diminished sequence diversity and low filter pass rates for BoltzGen minibinder designs in the 73-76 amino acid length range. This is visualized in an analysis of 33,600 designs against 5 protein and 4 small molecule targets in Figure 19. Inspection of the sequences revealed that the pipeline (backbone design followed by inverse folding) is frequently recapitulating the sequence for ubiquitin in this length range. For example, in this analysis all designs of length 73 (n=156) had >97% sequence identity to “MQIFVKTLTGKTITLEVEPSDTIENVKAKIQDKEGIP-PDQQRLIFAGKQLEDGRTLSDYNIQKESTLHLVLRL”.

**Figure 19:**
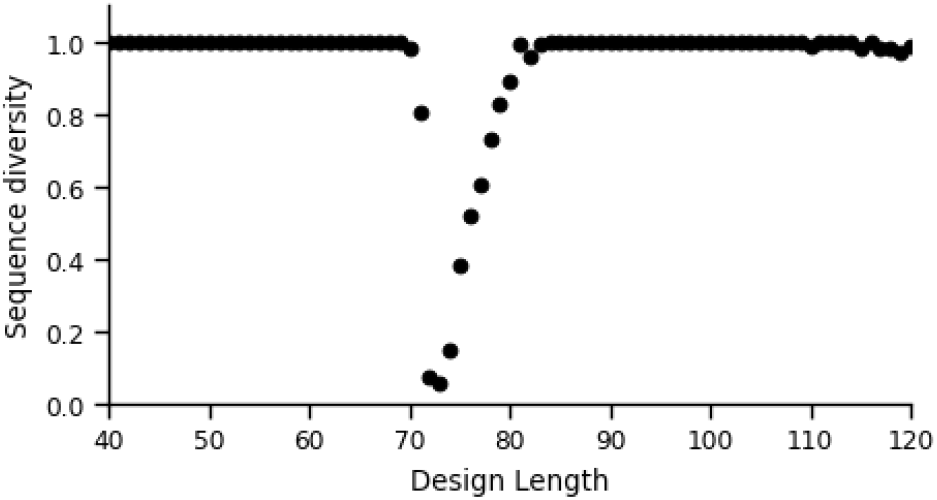
The Ubiquitin Memorization Issue. Shown is the sequence diversity (number of unique sequences divided by total number of designs) for designs with varying lengths against 9 targets (33,600 designs). The gap in the 73-76 region stems from BoltzGen’s bias toward Ubiquitin in that length range. The bias likely stems from Ubiquitin’s overrepresentation in the training data (>1000 entries in the PDB).

Likely, this arises since this sequence is present, often in complex, in >1000 entries in the PDB. In future versions of the model, we plan to down-sample this interaction during training.

## D Learnings from BoltzGenv0

### Template bug

A previous version of BoltzGen, which we term BoltzGenv0, had a flaw resulting in close-to-random ranking and filtering. The same failure case occurred for an attempt to design helical stapled peptides against a pMHC complex using BoltzGenv0.

The nature of this bug was in our handling of which residues are considered to be part of the target and which part of the design. In cases where the designed binder contains fixed residues, such as when designing helicons or nanobodies, BoltzGenv0 considered the fixed residues of the designs as being part of the target. The implications of this are that their relative position with respect to the target is provided in the refolding step via the templates that we employ for the targets. Thus, the resulting structure prediction is bound to recapitulate the generated structure, without providing any filtering power that enriches for binders. Furthermore, metrics such as the minimum interaction pAE and the ipTM would be influenced by the fixed residues that are in the design. For instance, in the overwhelming majority of cases, the minimum interaction pAE would not correspond to an interaction between the design and the target, but rather an interaction between a designed residue and a fixed residue in the designed binder. Hence, these scores also do not provide any filtering power for BoltzGenv0.

### Designfolding

Another improvement in BoltzGen over BoltzGenv0 is its “designfolding” step. In this step, we additionally refold the design in the absence of the target and compute the RMSD to the design in the generated structure. This serves as a proxy to assess whether the binder could attain the designed structure by itself, which we use to filter out designs that are likely to require a significant conformational change upon binding or do not express since they do not fold into a stable structure by themselves.

We introduced this improvement after a protein-protein binder design attempt where all 12 of BoltzGenv0’s designs failed to express. These designs would often partially or completely “envelop” the target and require a large conformational change to bind.

## E Wetlab Experimental Methods

### E.1 Nanobodies and Mini-proteins against Ten Novel and Five Benchmark Targets

#### Selection Process for Ten Novel Targets

Design campaigns frequently seek binders against novel targets for which no binders exist. Thus, we select a set of ten novel targets for which there are no existing bound structures in PDB. The detailed selection process is as follows:

1. **PDB Monomer** — The biological assembly has to be a monomer with exactly one protein polymer instance (`oligomeric_state = Monomer`, `polymer_entity_instance_count_protein = 1`).
2. **Monomer-only sequence cluster** — Each chain is either a singleton or a member of a sequence cluster (30% identity threshold, `mmseqs easy-cluster` with `–min-seq-id 0.30` and `-c 0.0`) in which *every* member is a PDB monomer (satisfies Condition 1).
3. **Catalog availability** — Must be available in the *Sino Biological* catalog (via mapping to a Swiss-Prot accession listed there).

With this selection method, we aim to ensure that each protein we keep is a **monomeric PDB entry** that has no close sequence homolog (MMseqs2 sequence identity ≥ 30%) anywhere in the PDB that appears in a multimeric or ligand-bound assembly. This makes the targets genuinely “hard” for our binder design: the model has not seen a closely related protein in a bound context during training.

We verify our target’s sequence identities are less than 30% to any non-monomeric protein in PDB as follows. For each target sequence, we ran MMseqs2 easy-search against non-monomer PDBs and kept the top identity hit that passed our coverage filter:

~~~
mmseqs easy-search queries.fa nonmonomer_db out.m8 tmp \
  --threads 32 \
  --min-seq-id 0.0 \
  --alignment-mode 3 \
  -e 1e5 -s 9.5 \
  --prefilter-mode 2 \
  --cov-mode 2 -c 0.9 \
~~~

- **Coverage policy**. We require *near-global coverage on the target* via `–cov-mode 2 -c 0.9`, i.e.

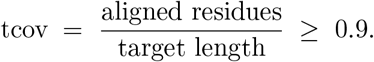

This enforces that the match spans (almost) the entire *target* chain. We initially used `-c 1.0`, but very long targets often fail to align end-to-end stably; we therefore relaxed to 0.9.

- **Zeros in the table**. Some entries appear as 0.0% (e.g., **1JQD, 2A1X**). This indicates that no hit met the tcov ≥ 0.9 threshold.

#### Biolayer Interferometry (BLI) affinity characterization

Ligand constructs were designed by reverse-translating target protein sequences and optimizing codon usage for expression in a prokaryotic cell-free system. A C-terminal assay tag was included to facilitate capture on biosensors. Gene fragments (Twist Bioscience) were assembled using NEBuilder HiFi DNA Assembly (NEB) in 2 *µ*L reactions. Assembly products were validated via capillary electrophoresis (Agilent ZAG DNA Analyzer) and quantified using the Qubit dsDNA assay (Invitrogen).

Proteins were expressed in 8 *µ*L reactions using an optimized in vitro transcription/translation system supplemented with 4 nM DNA template. Reactions were incubated at 37 °C for 8 hours. After expression, total protein levels were measured using an affinity-based detection assay and normalized across samples prior to binding analysis.

BLI experiments were carried out using a Gator Bio instrument with Strep-Tactin XT biosensors. Twin-Strep-tagged ligands were captured on the biosensors using the following protocol:

- **Baseline 1:** 120 s in running buffer
- **Ligand loading:** 120 s (target shift 0.5–1.0 nm)
- **Baseline 2:** 200 s in running buffer

The running buffer consisted of 50 mM HEPES, 100 mM NaCl, and 0.5% Triton X-100 at pH 7.4. All steps were performed at 25 °C with data collected at 5 Hz. Binding was assessed using a multi-cycle format with four antigen concentrations (30–1000 nM, half-log dilution series). Each kinetic cycle consisted of a 220 s association phase in antigen solution followed by a 240 s dissociation phase in running buffer. After each cycle, biosensors were regenerated with 10 mM glycine-HCl (pH 1.5) applied five times for 10 s each, followed by a wash step in running buffer to restore baseline. Buffer-only and non-binding ligand controls were used for reference subtraction and signal drift correction.

#### Surface Plasmon Resonance (SPR) affinity characterization

SPR measurements were performed on a Carterra LSA XT system. DNA constructs encoding ligands with C-terminal Twin-Strep tags were synthesized (Twist Bioscience), assembled using NEBuilder HiFi DNA Assembly, and validated using capillary electrophoresis and Qubit fluorometry. Proteins were expressed in a prokaryotic in vitro translation system and normalized post-expression using an affinity-based quantification assay. Sensor chip surfaces were functionalized by covalently attaching Strep-Tactin XT to a carboxymethylated surface using EDC/NHS coupling. The chip preparation procedure included:

- **Conditioning:** 50 mM NaOH
- **Activation:** EDC/NHS solution
- **Capture:** Strep-Tactin XT (50 *µ*g/mL in 10 mM sodium acetate, pH 4.5)
- **Quenching:** 1 M ethanolamine hydrochloride, pH 8.5
- **Wash:** 0.1 M sodium borate, 1 M NaCl, pH 9.0

Twin-Strep-tagged ligands were captured on the chip using a 96-channel printhead under bidirectional flow for 750 s, followed by a 600 s baseline step in running buffer. Antigens were diluted in running buffer (10 mM HEPES, 150 mM NaCl, 3 mM EDTA, 0.05% Tween-20, pH 7.4) and injected at seven concentrations (1–1000 nM, half-log dilution series) in a single-cycle kinetic format. Each cycle consisted of a 60 s baseline step in running buffer, a 300 s antigen association phase, and a 600 s dissociation phase in running buffer. After the completion of each injection series, the chip was regenerated with 10 mM glycine-HCl (pH 1.5) for 5 minutes, followed by a 20-minute wash in running buffer.

#### Data analysis and binder classification

Sensorgrams from both BLI and SPR assays were analyzed using Adaptyv Fitting software. Preprocessing included trimming to relevant kinetic phases (association and dissociation), correcting for signal jumps at buffer transitions, aligning phases, and subtracting signals from both baseline and reference channels.

Data were fit to a 1:1 Langmuir binding model using a global fitting approach across all antigen concentrations. If global fits were not feasible, alternative fitting strategies—such as dissociation-only or slope-based methods—were employed. In cases where individual curve fits could not be achieved, group-level models (e.g., equilibrium, flat, or linear) were used to approximate binding behavior.

Final kinetic parameters (*k*_on_, *k*_off_, and *K*_*D*_) were determined based on the best available fit. Under global fitting, *k*_off_ and *K*_*D*_ were estimated directly, with *k*_on_ calculated as *k*_off_*/K*_*D*_. Ligands were labeled as **binders (True)** or **non-binders (False)** based on the presence of quantifiable sensorgrams and successful kinetic model fitting. In cases where ligands generated a large signal during the association phase (at least 300% greater than the negative control) but could not be reliably fit, binding classification was assigned based on the observed magnitude of the shift.

#### Human Proteome Microarray Selectivity Profiling

Proteome-wide selectivity of biotinylated binder designs was assessed using the HuProt v4.0 Human Proteome Microarray (CDI Laboratories), which contains *>*21,000 full-length, individually purified human proteins and isoforms covering approximately 80% of the known human proteome. All proteins in the collection are expressed as N-terminal GST-His6 fusion proteins in Saccharomyces cerevisiae, purified via glutathione-agarose beads in a high-throughput 96-well format, Sanger sequence-verified, and printed in duplicate onto nitrocellulose-coated glass slides as previously described [Venkataraman et al., 2018]. Proteins are non-covalently yet irreversibly captured on the nitrocellulose surface. Each array additionally contains twenty blocks of control spots including titrated GST, BSA, biotinylated BSA, anti-biotin antibody, and fluorescent dye landmarks (Rhodamine + Alexa Fluor 647 IgG).

Prior to assay, arrays were blocked for 1 hour at room temperature with CDIArrayBlock buffer. Biotinylated binder constructs were diluted to 100 ng/mL or 1,000 ng/mL in CDIArrayBlock buffer and incubated on blocked arrays for 1 hour at room temperature with gentle shaking. Arrays were washed three times for 10 minutes each with TBST (1xTBS, 0.1% Tween-20), then incubated for 1 hour in a light-protected box with fluorescent streptavidin to detect biotin-tagged binders via the 635 nm channel. After three additional TBST washes and three rinses with ddH_2_O, arrays were dried under compressed CO_2_ and scanned on a GenePix 4000B scanner. A secondary-only control array (streptavidin without binder) was processed in parallel to capture background fluorescence arising from the small subset of human proteins that exhibit intrinsic streptavidin binding.

The target protein was represented on the arrays both by the standard HuProt collection spots and by two additional custom spots printed specifically for this study. Raw fluorescence intensities were extracted from .gpr files as median foreground values averaged across duplicate spot pairs and log2-transformed. Hits were called when the sample signal exceeded the secondary-only signal for that specific spot by more than 4-fold (*log*_2_ > 2), and the Z-score within that array exceeded 3, following the standard CDI Labs informatics pipeline [Venkataraman et al., 2018].

### E.2 Inhibiting Bioactive Peptides

#### Visual Inspection

The top 100 computationally ranked designs were manually examined in PyMOL. Visual inspection criteria included: (i) extent of peptide burial within the binding pocket, (ii) number and geometry of hydrogen bonds between binder and target peptide, (iii) overall packing density and complementarity at the binding interface, and (iv) internal packing quality of the apo binder in the absence of the target peptide. The top 6 designs exhibiting consistent burial, multiple well-oriented hydrogen bonds, and tightly packed interfaces were prioritized for experimental characterization.

#### Solid phase peptide synthesis and purification

Protegrin-1 was purchased from MedChemExpress (catalog #HY-P1633)/ Melittin, and Indolicidin were synthesized following the procedure below.

Melittin and Indolicidin were synthesized on a Biotage Initiator Alstra microwave synthesizer using standard Fmoc solid-phase peptide synthesis on a TentaGel S Ram resin. Resin (417 mg, 0.24 mmol) was swollen in DMF for 10-15 mins prior to synthesis. The general synthetic steps included: (a) Fmoc deprotection with 20% (v/v) piperidine in DMF, (b) resin washing (3x, DMF), (c) amino acid (0.50 mmol) coupling with DIPEA (0.50 mmol) and HCTU (0.50 mmol) for 5 min at 75 °C, (d) resin washing (3x, DMF), and (e) repeat deprotection/coupling until sequence completion.

The peptides were globally deprotected in a 10 mL solution of TFA:H2O:TIPS (95:2.5:2.5) for 3 h. The solution was then filtered, with the filtrate concentrated under the flow of nitrogen. The concentrate was precipitated in cold diethyl ether (40 mL), centrifuged, and the pellet dissolved in 5 mL H2O:ACN (1:1, 0.1% TFA).

Crude peptides were purified by reverse-phase HPLC on a C18 column using H2O/ACN (0.1% TFA) at a 10 mL/min gradient of 5-100% ACN (0.1% TFA) for 50 min. Pure fractions were identified by analytical HPLC and MALDI-TOF, pooled, and lyophilized. The final products were white powders with ≥95% purity.

#### Protein expression and purification

Codon-optimized genes encoding the designed candidate proteins with an N-terminal 6×His tag and a TEV protease cleavage site (HHHHHHENLYFQS) were synthesized and obtained from Twist Bioscience. To facilitate Gibson assembly into the pET-28a(+) vector, short sequences were added at the 5’ end (CTCTAGAAATAATTTTGTTTAACTTTAA-GAAGGAGATATACC) and 3’ end (GATCCGGCTGCTAACAAAGCCCGAAAG) of each gene. The recombinant plasmids were transformed into the Escherichia coli strain E. coli BL21(DE3). A single colony was picked from an LB agar plate and inoculated into LB medium supplemented with kanamycin (50 *µ*g/mL) for overnight growth. The culture was then transferred into 200 mL of TB medium containing kanamycin (50 *µ*g/mL) and incubated at 37 °C until reaching an OD600 of 0.6 - 0.8. Protein expression was induced with 0.5 mM isopropyl *β*-D-1-thiogalactopyranoside (IPTG), and cultures were incubated overnight at 30 °C. Cells were harvested by centrifugation and resuspended in 25 mL PBS buffer (10 mM Na2HPO4, 1.8 mM KH2PO4, 2.7 mM KCl, 137 mM NaCl, pH 7.4) supplemented with 20 mM imidazole. Cells were lysed by ultrasonication (Sonic Dismembrator Model 500, Fisher Scientific), and the lysate was clarified by centrifugation (35,000g, 30 min). The supernatant was loaded onto a gravity column containing Ni-NTA agarose resin (HisPur, Thermo Fisher, 1.0 mL or 3.0 mL). The resin was washed with three column volumes (CVs) of PBS buffer containing 20 mM imidazole, and bound proteins were eluted with 7 mL PBS buffer containing 250 mM imidazole. The eluted proteins were concentrated and subjected to three rounds of buffer exchange with PBS buffer using a 15 mL, 10 kDa cutoff centrifugal filter unit (EMD Millipore).

#### Circular dichroism

Protein samples were prepared at 10 *µ*M in sterile filtered 10 mM sodium phosphate with 50 mM NaCl, pH 7.4. A280 was measured via UV-Vis spectroscopy in a 0.1 mm Quartz cuvette, and protein concentration was calculated from A280 via Beer’s law using the extinction coefficient calculated from protein sequence via ExPasy ProtParam. Circular dichroism measurements were performed on a Jasco J-810 spectropolarimeter. Spectra were collected from 200-250nm in continuous scanning mode at 50 nm/min and 1nm band width with six accumulations per sample. CD spectra were converted from millidegrees to molar ellipticity using the equation *m · M/*(10· *L* ·*C*) where *C* is concentration in g/L (derived from A280 signal in UV-Vis experiments), *M* is the average molecular weight (g/mol), and *L* is the path length of the cell.

#### Analytical size exclusion chromatography

The oligomeric state of binder samples was assessed via analytical size exclusion chromatography using a Superdex 75 5/150 analytical gel filtration column (Cytiva) on an AKTA FPLC. Samples (50 *µ*L) were prepared at 100 *µ*M in sterile filtered 1X Phosphate Buffered Saline (PBS), pH 7.4, and centrifuged in a microfuge at 21,000g for 15 minutes before loading onto the FPLC. Chromatography runs were conducted at 0.2 mL/min for 1.2 column volumes and absorbance was measured at 220 and 280 nm. For measurement of protein:peptide complexes, protein and peptide were mixed at a 1:1 ratio (50 *µ*M each) and incubated overnight at 4 °C. After equilibration to room temperature, samples were centrifuged at 21,000g for 15 minutes before loading onto the FPLC.

#### Change in intrinsic tryptophan fluorescence for *in vitro* assessment of peptide binding

*In vitro* binding was assessed via tryptophan quenching, in which either peptide or protein was held constant, with the other binding partner varied, depending on which species contained tryptophan residues. All peptides were solubilized in DMSO at > 1mg/mL prior to dilution into sterile filtered 1X PBS, pH 7.4, for binding experiments. All binding assays were conducted in PBS in non-binding 96-well half-area black plates (Corning 3686,) and tryptophan quenching was measured as endpoint fluorescence intensity measurements (*λ*_ex_ = 295 nm, *λ*_em_ = 330 nm) in a BioTek Synergy Neo-2 multi-mode plate reader. For melittin and indolicidin, which both contain tryptophan residues, assays were conducted with constant [peptide], and binder sequences were designed to exclude tryptophan. [Melittin] was fixed at 10 *µ*M and [melittin binder] was varied from 0 to 40 *µ*M. [Indolicidin] was initially fixed at 10 *µ*M, with [binder] varying from 0-20 *µ*M. For subsequent global fitting experiments, [Indolicidin] was held constant at either 5, 7.5, or 10 *µ*M and [binder] was varied from 0-25*µ*M. Samples were prepared in triplicate and incubated overnight at 4 °C. After equilibrating to room temperature for 30 minutes, tryptophan fluorescence was measured as described. As a control, a gradient of [binder] without peptide was included for background subtraction. For all samples, the A295 and A330 of the binder was below 0.1; thus, protein fluorescence was subtracted as background instead of being treated with inner filter effect correction. For protegrin, which contains no tryptophan residues, [binder] was held constant while [peptide] was varied. All protegrin binder designs contain at least one tryptophan. For initial experiments, [protegrin] was held constant at 5 *µ*M and [binder] was varied from 0-30*µ*M. Samples were prepared in triplicate and incubated at room temperature for 3h before tryptophan fluorescence was measured. Here, a gradient of [protegrin] without binder was included for background subtraction.

Binding was fit with the following✓quadratic binding equation in Prism (GraphPad): 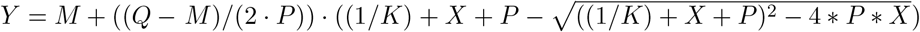 where *M* = fully unbound signal (baseline signal), *Q* = fully bound (saturation) signal, *P* = concentration of fixed species, *X* = ligand concentration, *Y* = observed fluorescence intensity, and *K* = association constant. *M, Q*, and *K* were not constrained, and *P* was fixed. For global fitting, *K* was shared for all datasets.

#### Surface plasmon resonance for validation of indo4-indolicidin binding

The binding of the highest affinity peptide/binder pair was further validated by surface plasmon resonance (SPR). SPR was performed on a Bruker SPR-24 Pro Instrument using an NTA derivatized SPR chip (SPR sensor prism NiHC1000M; Xantec bioanalytics). The surface was preconditioned with 350mM ethylenediaminetetraacetic acid (EDTA) and running buffer (10 mM HEPES pH 7.4, 150 mM NaCl, 50 µM EDTA, 0.05% Tween-20) prior to loading with 5mM Ni2+. Hexahistidine-tagged indo4 binder was immobilized on the surface prior to exposure to the analyte (indolicidin). Indolicidin was solubilized in water to 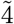mg/mL to afford a stock solution. From the stock solution, a concentration gradient of 0-20*µ*M indolidicin was prepared in running buffer. The analyte solutions were flowed over the immobilized protein surface for 80 seconds at a 25 *µ*L/min flow rate from low to high concentration and a 120-second dissociation time. Blank (running buffer only) injections were interspersed between the analyte injections to confirm that the analyte was dissociating between injections. Following the cycle of injections, binding affinity was calculated by plotting the pre-injection stop point signal (RU) versus protein concentration. High concentration samples were omitted because of a bulk shift from buffer mismatch in the preparation of the samples from the stock solution, solubilized in water. Affinity was calculated via a Langmuir fit of the response units (RU) at the pre-injection stop point.

#### Bacterial growth assays to measure neutralization of antimicrobial activity

Minimum inhibitory concentrations (MICs) of melittin, indolicidin, and protegrin-1 were determined against *Bacillus subtilis* (ATCC 23857). Peptides were prepared at 100 *µ*g/mL and serially diluted 2-fold in Mueller-Hinton Broth (MHB). A glycerol stock of *B. subtilis* was inoculated into 10 mL MHB and grown overnight at 37°C. The following day, 100 µL of starter culture was added to 10 mL MHB and grown to an OD600 of 0.6–0.8, then diluted to OD600 = 0.001 and added to the peptide dilutions in a 96-well non-treated cell culture plate (Gen-Clone 25-104). Absorbance at 600 nm was measured using a BioTek Synergy Neo-2 plate reader at 37°C with cycles of 7 min shaking and 3 min rest for 15 hours. The MIC was defined as the lowest peptide concentration that completely inhibited growth, and was 1.1 *µ*M, 1.70 *µ*M, and 1.16 *µ*M for melittin, indolicidin, and protegrin-1, respectively.

For neutralization assays, peptides were held constant at their respective MICs, and protein binders were serially diluted 2-fold starting from 40X their target peptide MIC. The peptide was added to the protein binders, transferred to a 96-well plate, and then the diluted *B. subtilis* culture was prepared as described above. Samples were prepared in triplicates. Absorbance at 600 nm was measured as previously described, with the exception that the indolicidin assays were conducted for 7.5 hours instead of 15 hours to account for peptide degradation. For the protegrin binders, protein and peptide dilutions were incubated overnight at 4°C before measuring absorbance. % neutralization was calculated with the following equation: % neutralization = ((*A*_obs_ − *A*_min_)*/*(*A*_max_ − *A*_min_))*cdot*100 where *A*_obs_ is the observed endpoint A600 for the varying protein concentrations, *A*_min_ is the observed endpoint A600 of the peptide and bacteria only control, and *A*_max_ is the observed endpoint A600 of the bacteria only control.

#### Hemolysis assays

Sheep red blood cells (25 mL) were transferred into a 50 mL conical tube and centrifuged at 500 x g for 5 min. The plasma layer was aspirated, leaving the pellet. The cells were resuspended in 150 mM NaCl solution to 25 mL, with gentle inversion and centrifugation (500 x g, 5 min). The supernatant was aspirated, and washed once more with 150 mM NaCl solution. The pellet was then resuspended in 1X PBS (pH 7.4), centrifuged (500 x g, 5 min), and aspirated. The pellet after the PBS wash, was then resuspended in 1X PBS to 25 mL and stored at 4 °C. For hemolysis assessment, 190 *µ*L of RBC (1:100 in 1X PBS) was added per well of a 96-well plate, followed by the addition of 10 *µ*L of either 1X PBS, 20% Triton X-100, or melittin. Cells were treated with serial dilutions of melittin (0.08 to 10 *µ*M final). The plate was incubated at 37 °C with gentle shaking for 1 h, followed by centrifugation (500 x g, 5 min). From each well, 100 *µ*L of supernatant was transferred to a fresh plate, ensuring pellets were undisturbed. Absorbance was measured at 400 nm using a microplate reader (SpectraMax M5). Values were normalized to PBS and Triton X-100 controls (N= 4, performed in duplicate). To assess melittin’s hemolytic activity in the presence of its binders, 10 *µ*L of melittin (1.2 *µ*M final) was added to wells of a 96-well plate, followed by serial dilution of protein (0.05 *µ*M to 6 *µ*M final). This was incubated for 1h at room temperature. Then, 180 *µ*L of red blood cells (1:100 in 1X PBS) was added to each well and incubated at 37 °C for 1 h. After centrifugation, 100 *µ*L of supernatants were transferred to a fresh plate and absorbance measured at 400 nm. Absorbance values were normalized to PBS and melittin only (1.2 *µ*M) controls (N= 4, performed in duplicate).

### E.3 Small Molecule Binders and Inhibitors

#### E.3.1 Rucaparib Binders

##### Rational inspection

The top 100 computationally ranked designs were examined based on the number and geometry of potential hydrogen bonds formed between rucaparib and each designed binder. Rucaparib was conceptually fragmented into three hydrogen-bonding functional groups: carboxamide, indole NH, and secondary amine. Hydrogen bonds were defined by the distance between oxygen or nitrogen atoms of these rucaparib fragments and those of the binder residues within 3.2 Å. The presence of hydrogen bonds involving the carboxamide group was given the highest priority during selection. Six candidate designs were subsequently chosen by visual inspection, considering both burial within the binding pocket and diversity of the protein scaffolds.

##### Protein expression and purification

Codon-optimized genes encoding the designed candidate proteins with an N-terminal 6×His tag and a TEV protease cleavage site (HHHHHHENLYFQS) were synthesized and obtained from Twist Bioscience. To facilitate Gibson assembly into the pET-28a(+) vector, short sequences were added at the 5’ end (CTCTAGAAATAATTTTGTTTAACTTTAA-GAAGGAGATATACC) and 3’ end (GATCCGGCTGCTAACAAAGCCCGAAAG) of each gene. The recombinant plasmids were transformed into the Escherichia coli strain E. coli BL21(DE3). A single colony was picked from an LB agar plate and inoculated into LB medium supplemented with kanamycin (50 *µ*g/mL) for overnight growth. The culture was then transferred into 200 mL of TB medium containing kanamycin (50 *µ*g/mL) and incubated at 37 °C until reaching an OD600 of 0.6 - 0.8. Protein expression was induced with 0.5 mM isopropyl *β*-D-1-thiogalactopyranoside (IPTG), and cultures were incubated overnight at 30 °C. Cells were harvested by centrifugation and resuspended in 25 mL PBS buffer (10 mM Na2HPO4, 1.8 mM KH2PO4, 2.7 mM KCl, 137 mM NaCl, pH 7.4) supplemented with 20 mM imidazole. Cells were lysed by ultrasonication (Sonic Dismembrator Model 500, Fisher Scientific), and the lysate was clarified by centrifugation (35,000 g, 30 min). The supernatant was loaded onto a gravity column containing Ni-NTA agarose resin (HisPur, Thermo Fisher, 1.0 mL or 3.0 mL). The resin was washed with three column volumes (CVs) of PBS buffer containing 20 mM imidazole, and bound proteins were eluted with 7 mL PBS buffer containing 250 mM imidazole. The eluted proteins were concentrated and subjected to three rounds of buffer exchange with PBS buffer using a 15 mL, 10 kDa cutoff centrifugal filter unit (EMD Millipore).

##### Fluorescence emission and fluorescence polarization assays

Fluorescence emission and fluorescence polarization spectra for assessment of rucarparib binding: To assess rucaparib binding, rucaparib dissolved in DMSO was mixed with proteins in PBS buffer (137 mM NaCl, 2.7 mM KCl, 10 mM Na2HPO4, 1.8 mM KH2PO4, pH 7.4) to a final DMSO concentration below 2%, and incubated for 5 min prior to measurement. Fluorescence emission spectra were recorded in black, flat-bottom 96-well plates using a BioTek Synergy Neo-2 plate reader with an excitation wavelength of 355 nm. Protein aliquots from 10 or 100 *µ*M stocks in PBS were combined to make 200 *µ*L samples containing 10 *µ*M rucaparib. Each condition was measured in triplicate. Fluorescence polarization (FP) assays were performed using the same samples on a BioTek Synergy 2 plate readerequipped with excitation and emission filters of 405 nm and 516 nm, respectively. FP values were recorded in polarization (P) units. The polarization values were plotted against protein concentration, and the data were fitted to a one-site binding model using nonlinear regression in GraphPad Prism 10 to determine the dissociation constant (*K*_*d*_).

#### E.3.2 Heme Binders

##### Rational inspection

The top 50 computationally ranked designs were visually inspected to ensure the heme was buried within the protein core, while the ligand placeholder remained exposed to the solvent to allow substrate access to the catalytic site. Structure predictions for the potential candidates were generated using Chai-1. From these, three candidates were selected based on their high structural similarity (backbone RMSD ≤ 1.0 Å) to the original Boltzgen design models.

##### Protein expression and purification

Codon-optimized genes encoding the designed candidate proteins with an N-terminal 6×His tag and a TEV protease cleavage site (HHHHHHENLYFQS) were synthesized and obtained from Twist Bioscience. To facilitate Gibson assembly into the pET-28a(+) vector, short sequences were added at the 5*′* end (CTCTAGAAATAATTTTGTTTAACTTTAA-GAAGGAGATATACC) and 3*′* end (GATCCGGCTGCTAACAAAGCCCGAAAG) of each gene. The recombinant plasmids were transformed into Escherichia coli strain E. cloni BL21(DE3). A single colony was picked from an LB agar plate and inoculated into LB medium supplemented with kanamycin (50 *µ*g/mL) for overnight growth. The culture was then transferred into 200 mL of TB medium containing kanamycin (50 *µ*g/mL) and incubated at 37 ◦C until reaching an OD600 of 0.6 - 0.8. Protein expression was induced with 0.5 mM isopropyl *β*-D-1-thiogalactopyranoside (IPTG), and cultures were incubated overnight at 30 ◦C. Cells were harvested by centrifugation and resuspended in 25 mL PBS buffer (10 mM Na2HPO4, 1.8 mM KH2PO4, 2.7 mM KCl, 137 mM NaCl, pH 7.4) supplemented with 20 mM imidazole. Cells were lysed by ultrasonication (Sonic Dismembrator Model 500, Fisher Scientific), and the lysate was clarified by centrifugation (35,000 g, 30 min). The supernatant was loaded onto a gravity column containing Ni-NTA agarose resin (HisPur, Thermo Fisher, 1.0 mL or 3.0 mL). The resin was washed with three column volumes (CVs) of PBS buffer containing 20 mM imidazole, and bound proteins were eluted with 7 mL PBS buffer containing 250 mM imidazole. The eluted proteins were concentrated and subjected to three rounds of buffer exchange with PBS buffer using a 15 mL, 10 kDa cutoff centrifugal filter unit (EMD Millipore).

##### UV-vis spectroscopy assays

UV-vis spectrometry were used to determine the binding affinities of the designs to FeDPP and iron heme (Protophophyrin IX). To assess the binding the cofactors, 1mM FeDPP or heme were dissovled in DMSO and mixed with proteins in PBS buffer (137 mM NaCl, 2.7 mM KCl, 10 mM Na 2 HPO 4, 1.8 mM KH 2 PO 4, pH 7.4) for a final cofactor concentration of 5*µ*M, and increasing concentration of protein (0.05, 0.1, 0.25, 0.5, 0.75, 1.0, 2.0, and 5.0 equivalent of cofactor) to a total volume of 200 *µ*L. The mixture was incubated for 1 hour prior to measurement. The spectra from 250 to 800nm were recorded on a UV-vis spectrometer. No concentration-dependent spectral change was observed for FeDPP titration, most likely due to self-aggregration of the cofacotr in aqueous buffer. For titration of proteins to heme, the absorbance at 413nm was used for dissociation constant (K d) determinination, using a quadratic ligand binding equation [Hulme and Trevethick, 2010].

#### E.3.3 Inhibitors against Brilacidin

##### Protein purification and expression of brilacidin binders

Synthetic, codon-optimized genes encoding the designed candidate proteins with a C-terminal 6×His tag and a TEV protease cleavage site (ENLYFQSGGHHHHHH) were obtained from Twist Bioscience. To facilitate Gibson assembly into the *pET-28a(+)* vector, short sequences were added at the 5’ end (CTTTAAGAAGGAGATATACCATGG) and 3’ end (CTCGAGCACCACCACCACCACCAC) of each gene. The recombinant plasmids were transformed into *Escherichia coli* strain BL21(DE3). A single colony was picked from an LB agar plate and inoculated into LB medium supplemented with kanamycin (50 *µ*g/mL) for overnight growth. The culture was then transferred into 1 L of LB medium containing kanamycin (50 *µ*g/mL) and incubated at 37 ◦C until reaching an OD_600_ of 0.6–0.8. Protein expression was induced with 1 mM isopropyl *β*-D-1-thiogalactopyranoside, and cultures were incubated for 4 hours at 37 ◦C. Cells were harvested by centrifugation and resuspended in 25 mL PBS buffer (10 mM Na_2_HPO_4_, 1.8 mM KH_2_PO_4_, 2.7 mM KCl, 137 mM NaCl, pH 7.4). Cells were lysed by ultrasonication (Sonic Dismembrator Model 500, Fisher Scientific), and the lysate was clarified by centrifugation (35,000 g, 30 min). The supernatant was loaded onto a gravity column containing Ni-NTA agarose resin. The resin was washed with three column volumes of PBS buffer containing 20 mM imidazole, and bound proteins were eluted with 20 mL PBS buffer containing 300 mM imidazole. The eluted proteins were concentrated and subjected to an overnight buffer exchange using 10 kDa cutoff dialysis cassettes (Slide-A-Lyzer, ThermoFisher) in PBS. Further purification of designs bri1 and bri5 were performed via FPLC using a Superdex75 Increase 10/300 (cm/cm) column. Proteins were concentrated to 1 mL and injected onto the column with a flow rate of 0.5 mL/min with UV detection at 220 nm and 280 nm. The monomeric species were then collected and tested for neutralization.

##### Bacterial growth assays to measure neutralization of antimicrobial activity of brilacidin

A glycerol stock of *E. coli* K-12 was inoculated into 10 mL Mueller Hinton Broth (MHB) and grown overnight at 37 ◦C. The following day, 100 *µ*L of starter culture was added to 10 mL MHB and grown to an OD_600_ of 0.6–0.8, then diluted to OD_600_ = 0.001. Brilacidin concentration was held constant at its *E. coli* minimum inhibitory concentration (MIC) of 1.56 *µ*M and designed proteins (bri1–bri5) were serially diluted 2-fold in PBS from 62.4 *µ*M to 0.98 *µ*M protein concentration. Bacteria were then added to the protein dilutions and brilacidin in a 96-well non-treated cell culture plate (Gen-Clone 25-104) in triplicate. Absorbance at 600 nm was measured using a BioTek Synergy Neo-2 plate reader at 37 ◦C with cycles of 7 min shaking followed by 3 min resting for 15 hours.

Initial neutralization assays (Supplementary Figure 27.) were tested with proteins prepared after Ni-NTA elution and buffer exchange, which show that designs bri1 and bri5 are the most potent neutralizers. Bri1 and bri5 were then retested for neutralization after further purification from the FPLC.

### E.4 Peptide Binders Targeting Rag GTPase

#### Materials

Peptides targeting Rag C were synthesized by Genscript.

#### Expression and purification of the Rag A: Rag C GTPase heterodimer

*E. coli* LOBSTR [Andersen et al.,2013] cells carrying a pETDuet-1 vector encoding codon optimized, C-terminally His-tagged RagA with a mutation (T21N) that favors the GDP-loaded state [Yang et al.,2020,Kim et al.,2008,Shen et al.,2017] and tagless wildtype (WT) RagC in its state (Addgene: 99664), were grown in Terrific broth (TB) and protein expression induced with an overnight IPTG treatment at 18 ◦C b. The purification follows a protocol described previously [Shen et al.,2017]. The complex was purified using Ni-NTA affinity chromatography followed by ion-exchange chromatography (IEX) using a Capto HisRes Q anion-exchange column and then size-exclusion chromatography (SEC) using a Superdex200 column. The protein corresponding to the heterodimer was concentrated in a final buffer containing 50 mM HEPES (pH7.5), 100 mM NaCl and 2 mM MgCl2 and used for SPR studies.

#### Binding studies of peptides to the Rag GTPase heterodimer using a high-throughput SPR instrument

Surface plasmon resonance (SPR) experiments were performed on a Cytiva Biacore 8K instrument. 0.2 *µ*M of the Rag GTPase heterodimer with the His-tag on RagA was immobilized on a Biacore NTA sensor chip using a Ni-NTA-Histag immobilization technique. All the peptides, at a concentration range from 0-100 *µ*M were flown over the protein-bound NTA sensor chip as ‘analyte’ and their binding responses were recorded. The Rag GTPase heterodimer was first loaded with 1 *µ*M GDP after immobilization prior to each peptide run. The protein, GDP, and the peptide samples were prepared in a buffer containing 50 mM HEPES (pH7.5), 100 mM NaCl and 2 mM MgCl2. Rag GTPase heterodimer was immobilized on the NTA sensor chip for 200 sec and all the runs were performed at 25 °C. For each peptide run at each concentration, the association and dissociation times were 120 and 300 sec, respectively. The sensor chip was regenerated after each run using 0.35 M EDTA and subsequently reused throughout the entire run.

Association and dissociation sensorgrams were plotted and binding affinities were analyzed using GraphPad Prism (Version 10.6.0). Each sensorgram corresponding to a single peptide concentration (100*µ*M) was plotted using GraphPad Prism (Version 10.6.0) with time in seconds in the x-axis and relative response in the y-axis. (Supplementary Figures 30 and 32). For peptides that showed detectable as well as suboptimal binding response (Supplementary Figures 30,32 and Tables 13,15), a concentration-based binding curve with peptide concentration (*µ*M) in the x-axis and relative response (RU) in the y-axis were plotted based on the individual sensorgrams at each concentration. The fit was generated using GraphPad Prism (Version 10.6.0) using a non-linear regression curve fit with a ‘one-site specific binding’ model and a single dissociation constant (KD) was computed (Supplementary Figures 31, 33 and Tables 14,16).

**Table 13:**
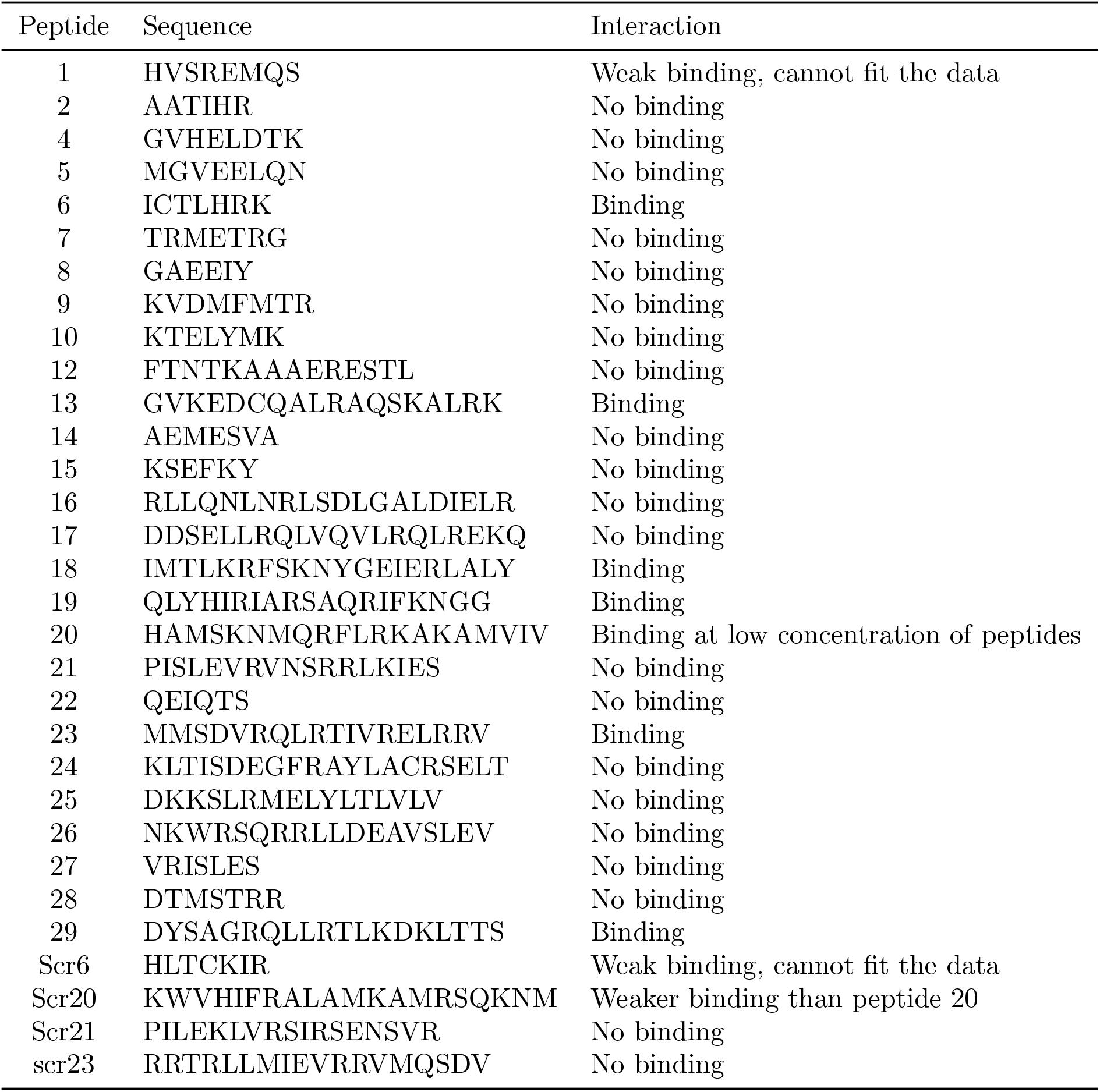
Binding of linear peptides to Rag GTPase designed with BoltzGen.

**Table 14:** Dissociation constant of binding of linear peptides designed with BoltzGen.

| Peptide | Sequence | KD in μM |
| --- | --- | --- |
| 6 | ICTLHRK | 48 |
| 13 | GVKEDCQALRAQSKALRK | 137 |
| 18 | IMTLKRFSKNYGEIERLALY | 1073 |
| 19 | QLYHIRIARSAQRIFKNGG | 400 |
| 20 | HAMSKNMQRFLRKAKAMVIV | 11 |
| 23 | MMSDVRQLRTIVRELRRV | 300 |
| 29 | DYSAGRQLLRTLKDKLTTS | 181 |
| Scr20 | KWVHIFRALAMKAMRSQKNM | 36.5 |

**Table 15:**
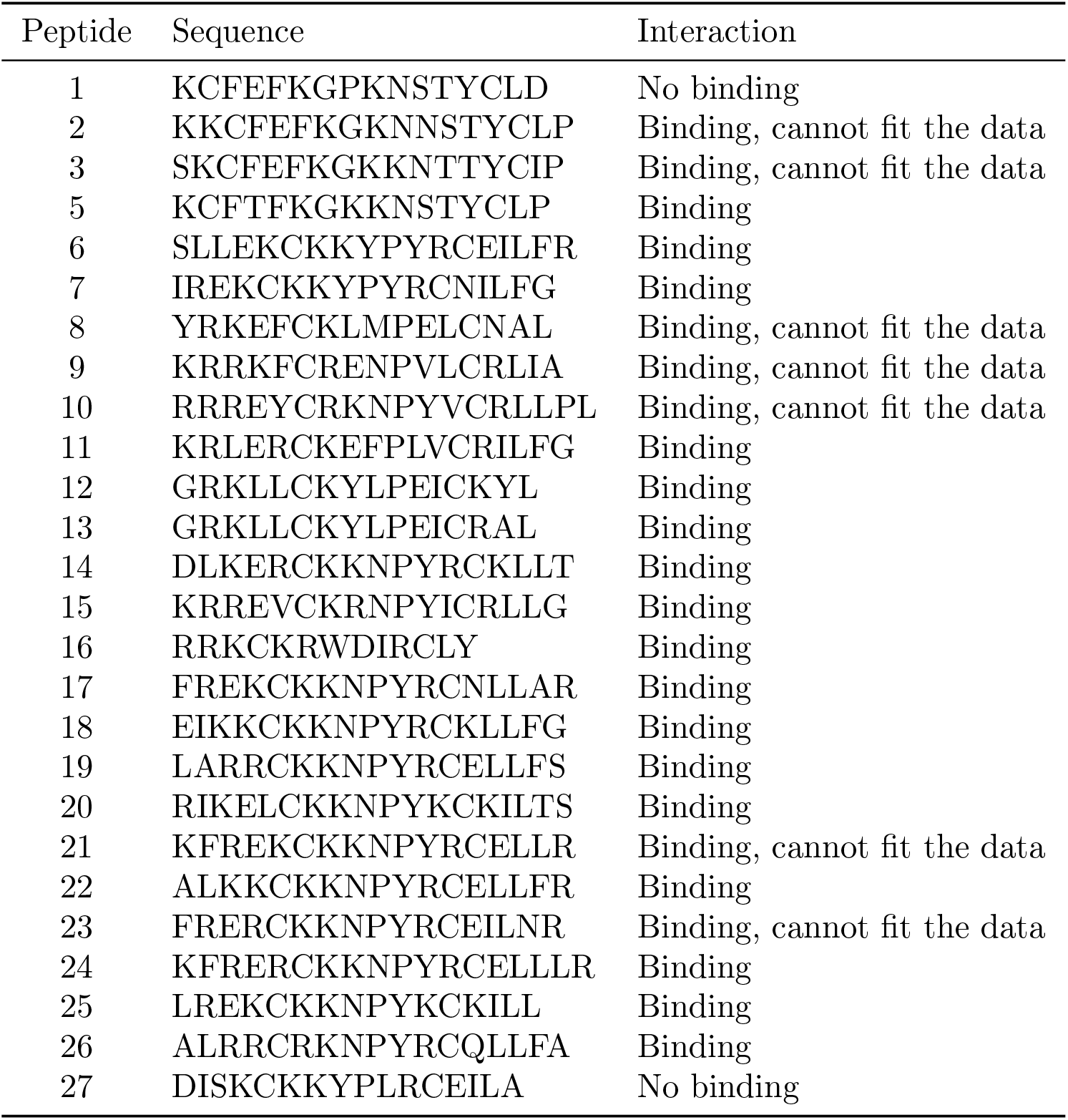
Binding of cyclic disulfide bonded peptides to Rag GTPase.

**Table 16:**
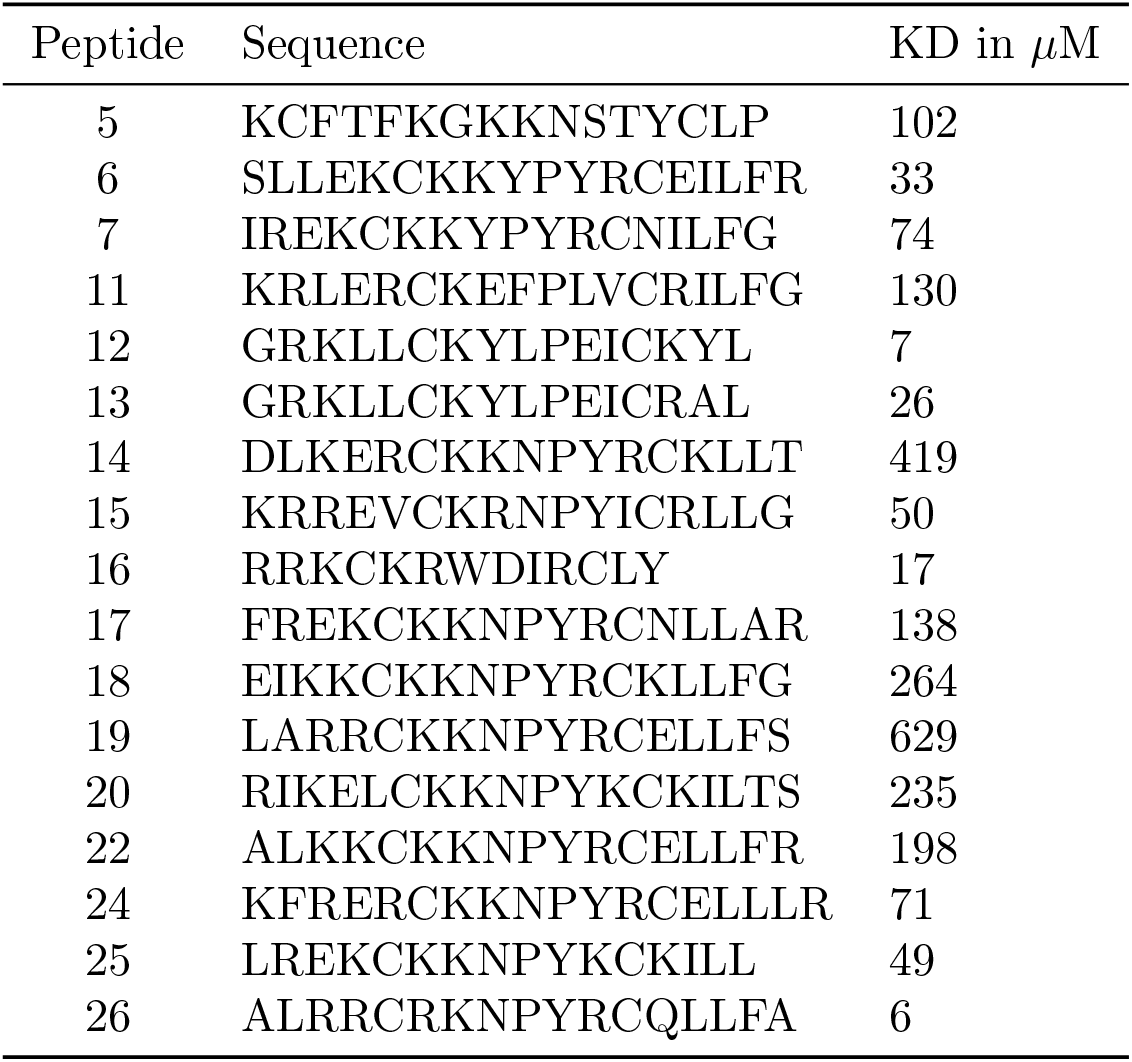
Dissociation constant of binding of cyclic disulfide bonded peptides.

| Peptide | Sequence | KD in μM |
| --- | --- | --- |
| 5 | KCFTFKGKKNSTYCLP | 102 |
| 6 | SLLEKCKKYPYRCEILFR | 33 |
| 7 | IREKCKKYPYRCNILFG | 74 |
| 11 | KRLERCKEFPLVCRILFG | 130 |
| 12 | GRKLLCKYLPEICKYL | 7 |
| 13 | GRKLLCKYLPEICRAL | 26 |
| 14 | DLKERCKKNPYRCKLLT | 419 |
| 15 | KRREVCKRNPYICRLLG | 50 |
| 16 | RRKCKRWDIRCLY | 17 |
| 17 | FREKCKKNPYRCNLLAR | 138 |
| 18 | EIKKCKKNPYRCKLLFG | 264 |
| 19 | LARRCKKNPYRCELLFS | 629 |
| 20 | RIKELCKKNPYKCKILTS | 235 |
| 22 | ALKKCKKNPYRCELLFR | 198 |
| 24 | KFRERCKKNPYRCELLLR | 71 |
| 25 | LREKCKKNPYKCKILL | 49 |
| 26 | ALRRCRKNPYRCQLLFA | 6 |

### E.5 Peptides Targeting Disordered Proteins

#### Generation of mammalian expression vectors for expression of GFP-tagged IDR binders

pRK5-msfGFP-binder plasmids were constructed by amplifying msfGFP from pRK5_msfGFP-HMGB1-Shuffled 1 (Addgene #237650) [Zhang et al., 2025]. The IDR binder sequences were inserted into the C-terminus of msfGFP through primer sequences. The amplicons were assembled into AgeI + XbaI-digested pRK5 backbone (Addgene #194548) [Mensah et al., 2023] using the NEBuilder HiFi DNA assembly master mix.

#### Cell culture

Cells were cultured under standard conditions (37 ◦C and 5% CO_2_) in sterile, TC-treated, non-pyrogenic, polystyrene tissue culture dishes (Corning). U2-OS (ATCC, HTB-96) cell line was cultured in DMEM GlutaMAX (Gibco, 31966047). Culture medium included 10% FBS (Gibco, 10438-026) and 100 U ml^−1^ penicillin–streptomycin (Gibco, 15140148). All of the cell lines tested negative for mycoplasma using the LookOut Mycoplasma PCR Detection Kit (Sigma-Aldrich, MP0035) or the PCR Mycoplasma Test Kit II (Applichem, A8994). Mycoplasma testing was performed on 0.2–1 ml of cell culture medium taken from tissue culture dishes containing confluent monolayers of cells on a routine basis at least twice a year.

#### Live-cell imaging

All live-cell imaging experiments were performed using the LSM880 Airyscan microscope equipped with a Plan-Apochromat ×63/1.40 oil differential interference contrast objective, while incubating cells at 37 ◦C and 5% CO_2_. Cells were seeded onto eight-well chamber slides (Ibidi, 80826-90) at 40,000 cells per well, transfected 24 h later, and imaged 24 h after transfection. U2OS cells were transfected using FuGENE HD according to the manufacturer’s instructions. Hoechst 33342 (0.2 *µ*g ml^−1^, Thermo Fisher Scientific, 62249) was added into the cell culture medium for nuclear staining.

#### Immunofluorescence

For immunofluorescence experiments performed in U2OS cells, cells were seeded on 8-well chamber slides (Ibidi, 80826-90) with 40,000 cells per well, and transfected 24 h later and fixed 24 h after transfection with 4% PFA in PBS for 10 min. Cells were permeabilized with 0.5% Triton X-100 (Thermo Fisher Scientific, 85111) in PBS for 30 min, incubated in blocking buffer containing 1% BSA (BSA Fraction V, Gibco, 15260037) and 0.1% Triton X-100 in PBS for 1 hr, followed by staining with primary antibodies at room temperature for 1 hr with gentle rotation. Slides were washed five times with blocking buffer, incubated with secondary antibodies (AlexaFluor 594 donkey anti-mouse antibody, Jackson ImmunoResearch, 715-585-150; and AlexaFluor 594 donkey anti-mouse antibody, Jackson ImmunoResearch 711-605-152, 1:1,000) in blocking buffer for 1 h at room temperature, washed twice with blocking buffer, stained with 0.5 *µ*g ml^−1^ DAPI in PBS (Invitrogen, D1306), and washed three times with PBS. The following primary antibodies were used: NPM1 (B23) (Santa Cruz, sc-271737, 1:100), SURF6 (Abcam, ab221990, 1:1000). Imaging was performed using the LSM880 Airyscan microscope equipped with a Plan-Apochromat ×63/1.40 oil differential interference contrast objective.

#### Image analysis

Image analysis of the NUP98-binder expression assay was performed using ZEN Blue v.3.12 software. Nuclear segmentation was performed using Zeiss AI-powered nuclear segmentation software trained on Hoechst 3342-stained U2OS cells. Nuclear annotations were used to compute single-cell mean GFP intensity, mean mCherry intensity, and mCherry standard deviation. Nuclei with mean mCherry intensities less than 50 were thrown out of analysis. Further image processing was performed using FIJI (v.2.16.0).

### E.6 Nanobodies against Recently Deposited and Novel Pathogen Targets

The plots in Figure 20 represent the median Alexa Fluor 647 fluorescence intensity of nanobody-displaying cells (HA tag label positive) minus the median Alexa Fluor 647 fluorescence intensity of the non-nanobody-displaying cells (HA tag label negative) at different concentrations of antigen. The latter acts as an internal control and is subtracted to remove the signal from nanobody-independent background binding of cells to antigen. Higher fluorescence intensity indicates more binding. The background-subtracted median fluorescence intensities across concentrations for each nanobody design were then fit to a Hill function using `SciPy curve_fit`. For initializing curve fitting, the background value used was the minimum fluorescence intensity observed at 0 antigen concentration and the EC_50_ used was the median antigen concentration tested. The Hill coefficient, *n*, was explicitly set to 1, as we expect non-cooperative binding of the antigen to the antibody. The fluorescence value, *Y*, is fit as:

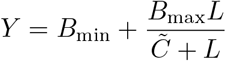

where *B*_min_ is the best-fit background fluorescence value, *B*_max_ is the best-fit maximum binding value, 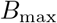 is the best-fit EC_50_ projection, and *L* is the concentration of antigen.

**Figure 20:**
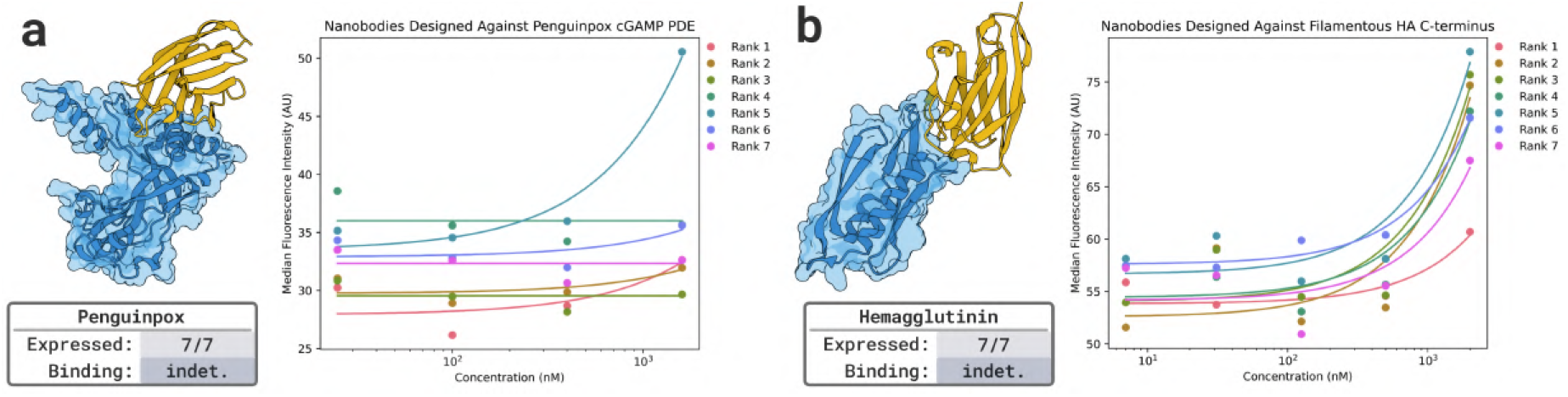
**a.** Penguinpox cGAMP PDE Binder Design. Structural model of the designed nanobody (yellow) in complex with penguinpox cGAMP PDE (blue), and nanobody binder YSD assays. **b**. Hemagglutinin FhaB Binder Design. Structural model of the designed nanobody (yellow) in complex with Hemagglutinin FhaB (blue), and nanobody binder YSD assays. The highest concentration of antigen in both cases was the only test case that demonstrated any binding signal for any nanobody design and no affinity can be determined.

**Figure 21:**
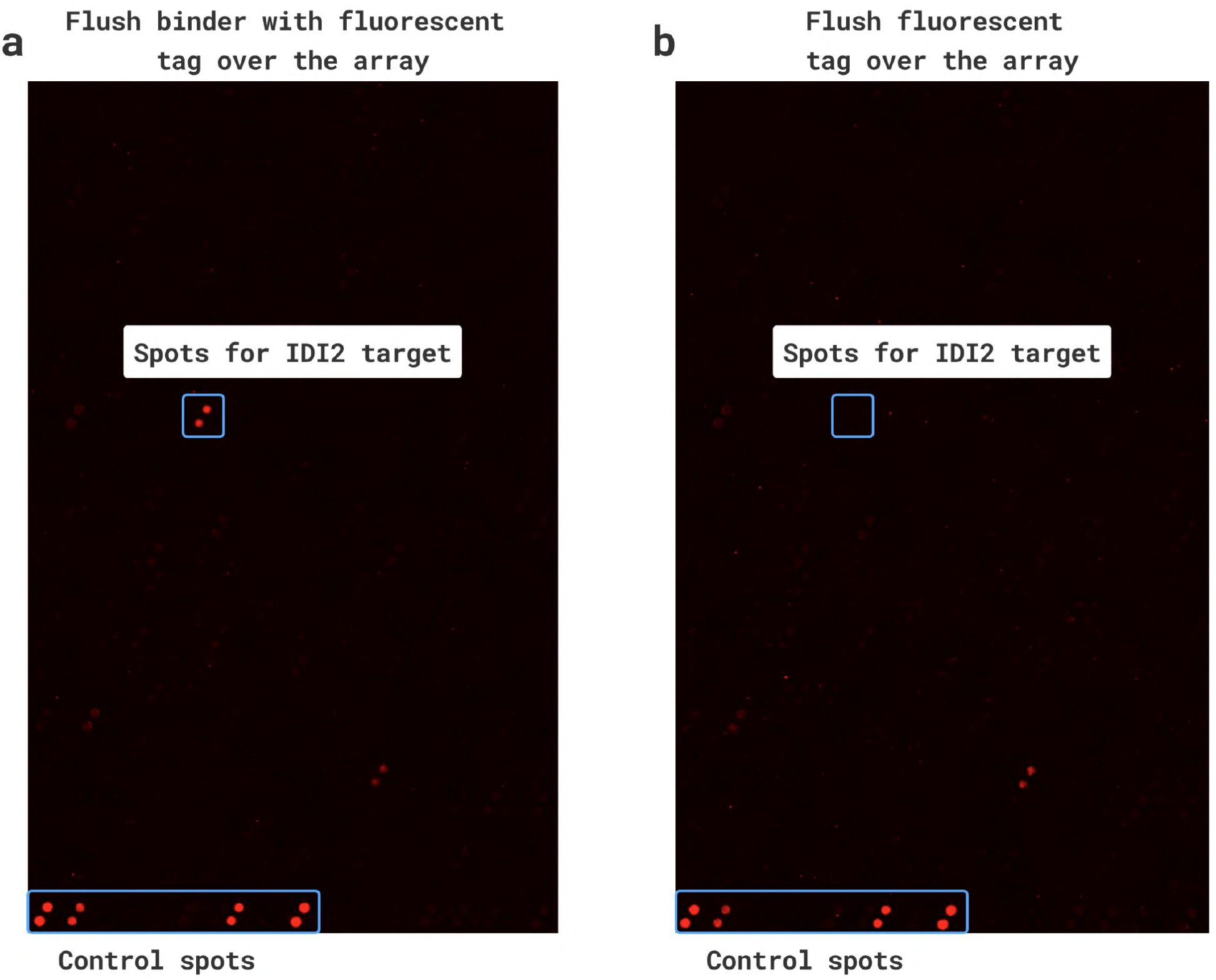
Proteome-wide selectivity profiling of a biotinylated mini-protein binder against IDI2. Off-target binding was assessed using the HuProt v4.0 Human Proteome Microarray (CDI Labs), in which *>*21,000 full-length human proteins are printed in duplicate on nitrocellulose-coated glass slides. Arrays were probed with a biotinylated binder design at 100 ng/mL followed by fluorescent streptavidin (panel a), or with fluorescent streptavidin alone as a secondary-only control (panel b). The sole signal exceeding the 4-fold secondary-only threshold (Z-score *>* 3) corresponds to the intended target IDI2, with no off-target interactions detected across the remainder of the human proteome.

**Figure 22:**
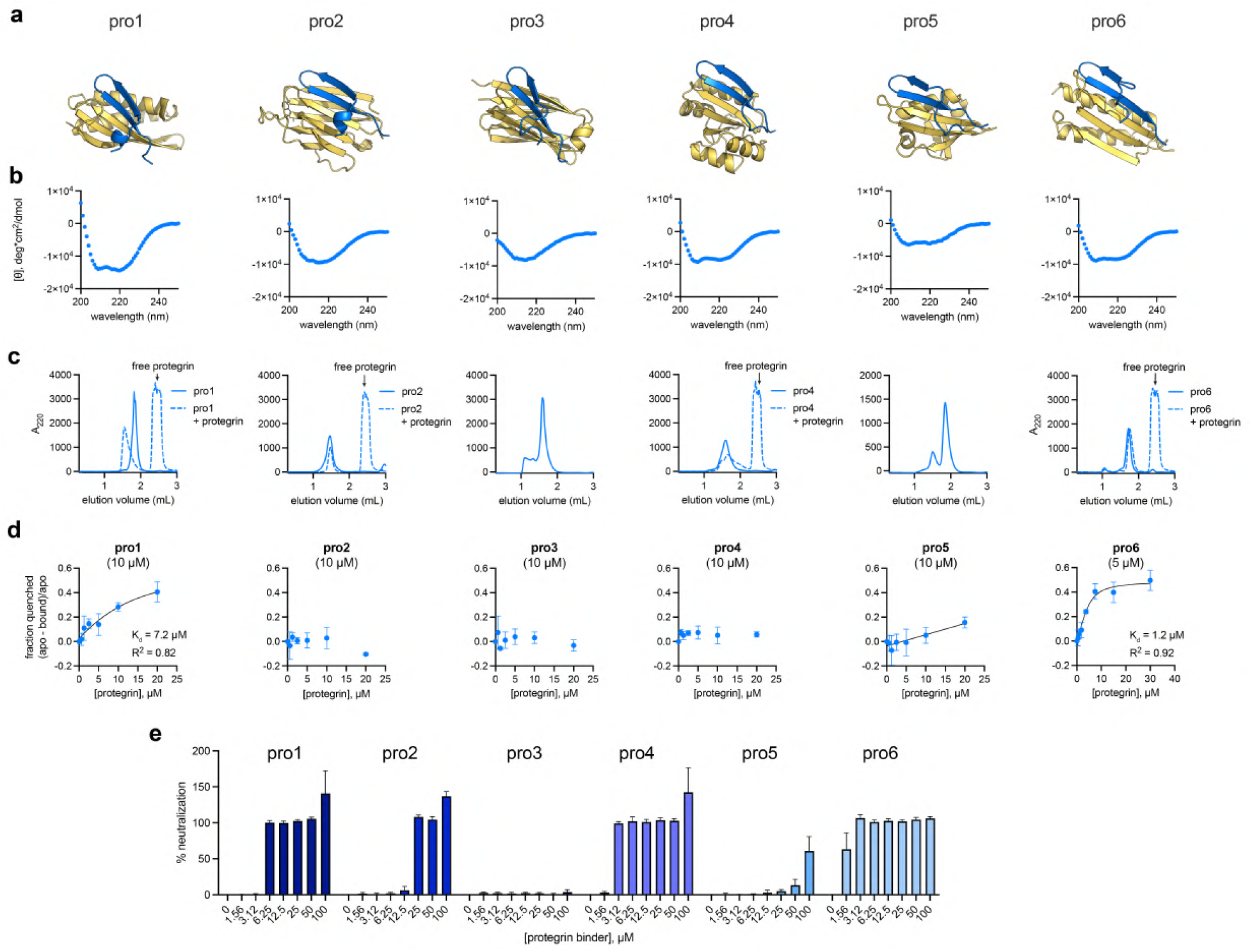
Experimental characterization of protegrin binder designs. **(a)** Structure prediction models of all selected protegrin binders are predicted to adopt a helical conformation. **(b)** Circular dichroism shows that Pro1, 4, 5, and 6 have partial helical character, while Pro2 and Pro3 have the expected *β*-structure in their apo form. **(c)** Analytical size exclusion chromatography traces of Pro1–Pro6 show varied oligomeric states. **(d)** Pro1, 5, and 6 show detectable indolicidin binding as measured by changes in intrinsic fluorescence of the binders’ tryptophan residues. **(e)** Pro1, 2, 4, and 6 fully neutralize protegrin antimicrobial activity against *B. subtilis* at the concentrations tested. (Data presented for pro6 is also presented in Figure 3)

**Figure 23:**
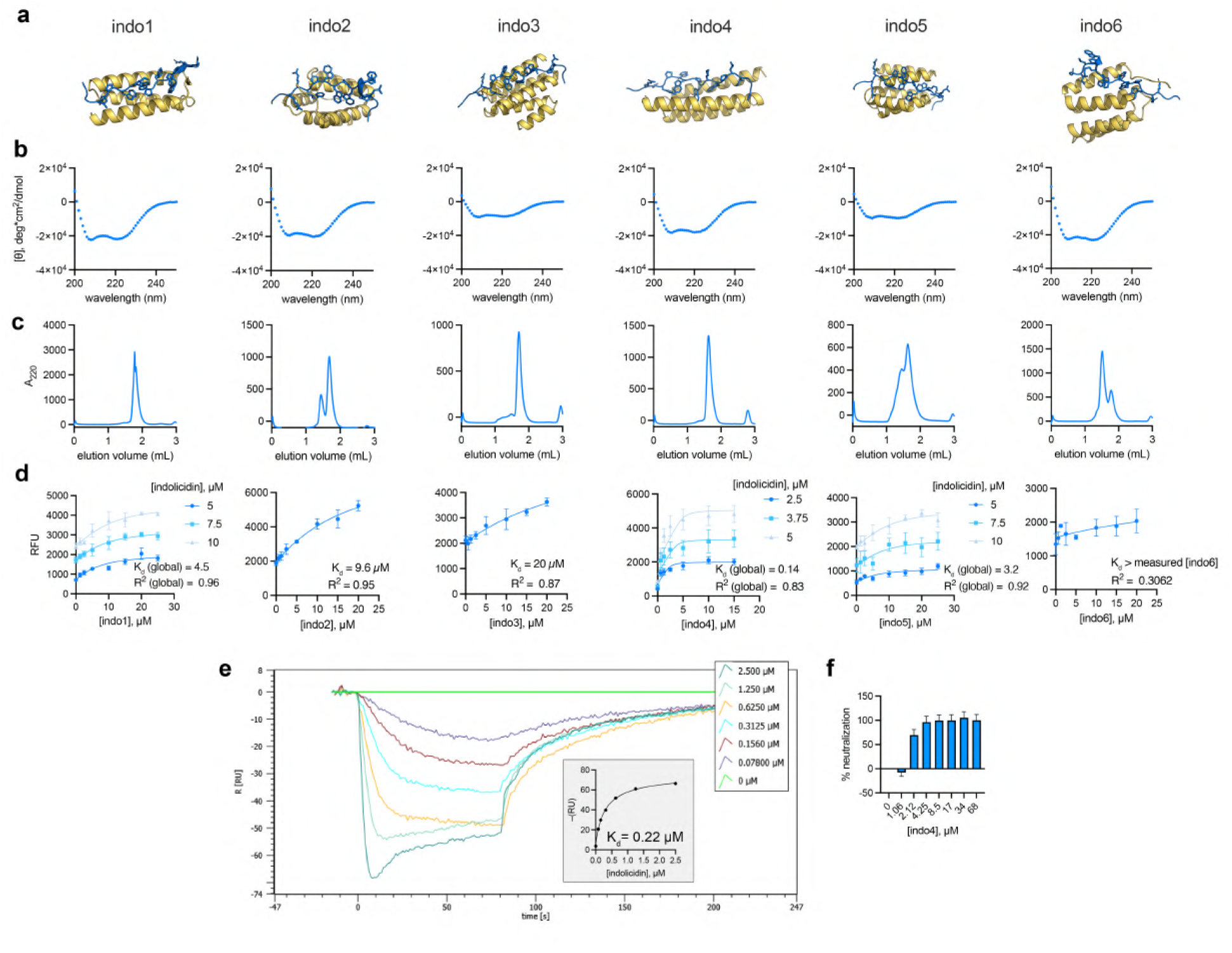
Experimental characterization of indolicidin binder designs. **(a)** Structure prediction models of all selected indolicidin binders are predicted to adopt a helical conformation. **(b)** Circular dichroism shows that Indo1–6 all have helical character. **(c)** Analytical size exclusion chromatography traces of Indo1–Indo6 show varied oligomeric states. **(d)** All Indo binders show detectable indolicidin binding as measured by changes in intrinsic fluorescence of indolicidin’s tryptophan residues. Indo1, Indo4, and Indo5 binding was measured with multiple indolicidin concentrations, and *K*_*d*_ was determined via a global fit. **(e)** Only Indo4 neutralizes indolicidin antimicrobial activity against *B. subtilis*. **(f)** For Indo4, binding was validated by surface plasmon resonance, in which Indo4 was immobilized and exposed to varied [indolicidin]. Affinity was determined as a function of response units at the pre-injection stop points of each [indolicidin]. (Data presented for indo4 is also presented in Figure 3)

**Figure 24:**
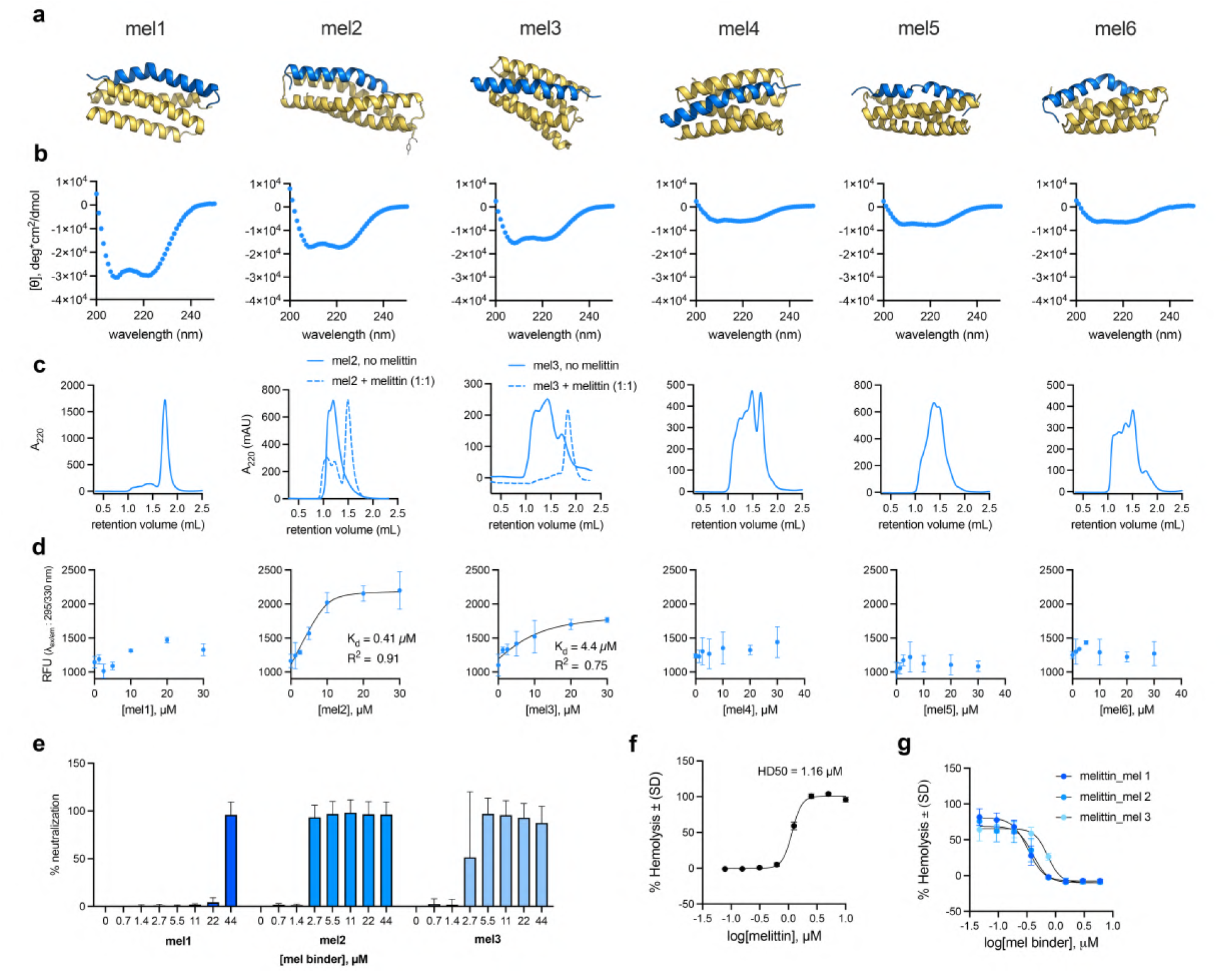
Experimental characterization of melittin binder designs. **(a)** Structure prediction models of all selected melittin binders are predicted to adopt a helical conformation and to bind melittin in its amphipathic helix state. **(b)** Circular dichroism shows that Mel1–3 have helical character, while Mel4–6 are only partially folded. **(c)** Analytical size exclusion chromatography traces of Mel1–Mel6 show varied oligomeric states. **(d)** Mel2 and Mel3 show detectable melittin binding as measured by changes in intrinsic fluorescence of melittin’s tryptophan residue. Melittin concentration is held constant at 10 µM while binders, which lack tryptophan residues, are varied from 0–40 µM. **(e)** Mel1–3 neutralize melittin’s antimicrobial activity against *B. subtilis*. **(f)** Mel1–3 also neutralize melittin’s hemolytic activity. (Data presented for mel2 is also presented in Figure 3)

**Figure 25:**
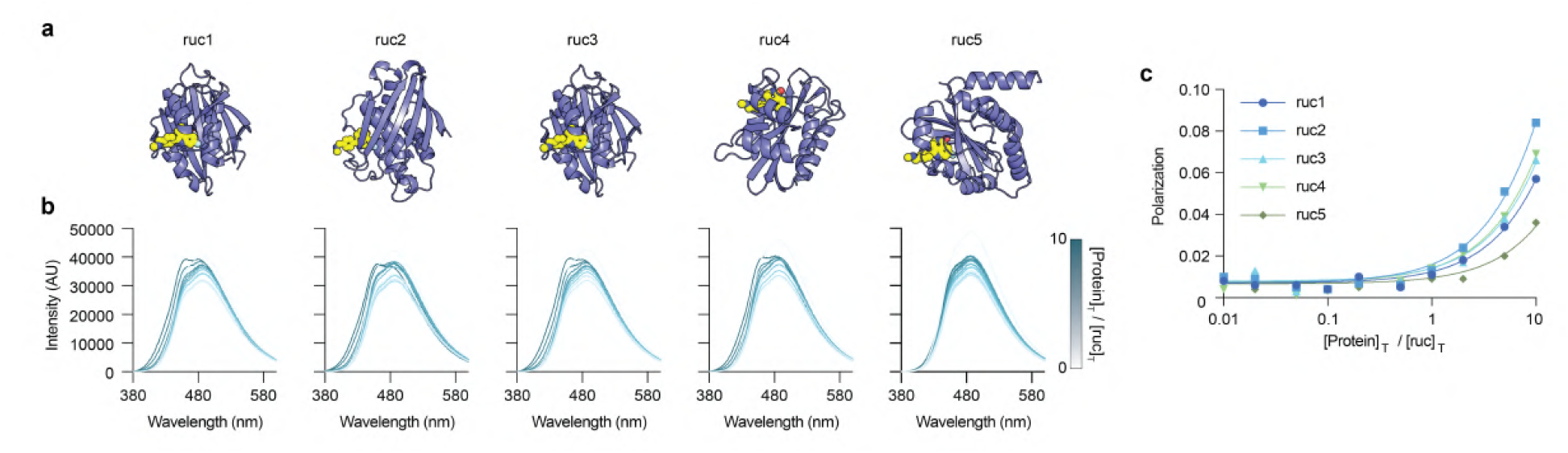
rucaparib Binder Design. **a)** Structural model of the designed rucaparib binder (purple) in complex with rucaparib (yellow). **b)** Fluorescence emission spectra of rucaparib (10 *µ*M) upon titration with increasing concentrations of the designed protein (0–10 equivalents). The emission spectrum of rucaparib exhibited a blue shift upon binding, indicating changes in its local environment. **c)** Fluorescence polarization assay of rucaparib binding to the designed protein, measured with an excitation wavelength of 405 nm and emission wavelength of 516 nm. Polarization values were plotted against the molar ratio of protein to rucaparib, and the data were fitted to a one-site binding model using nonlinear regression in GraphPad Prism 10 to determine the dissociation constant (*K*_*d*_).

**Figure 26:**
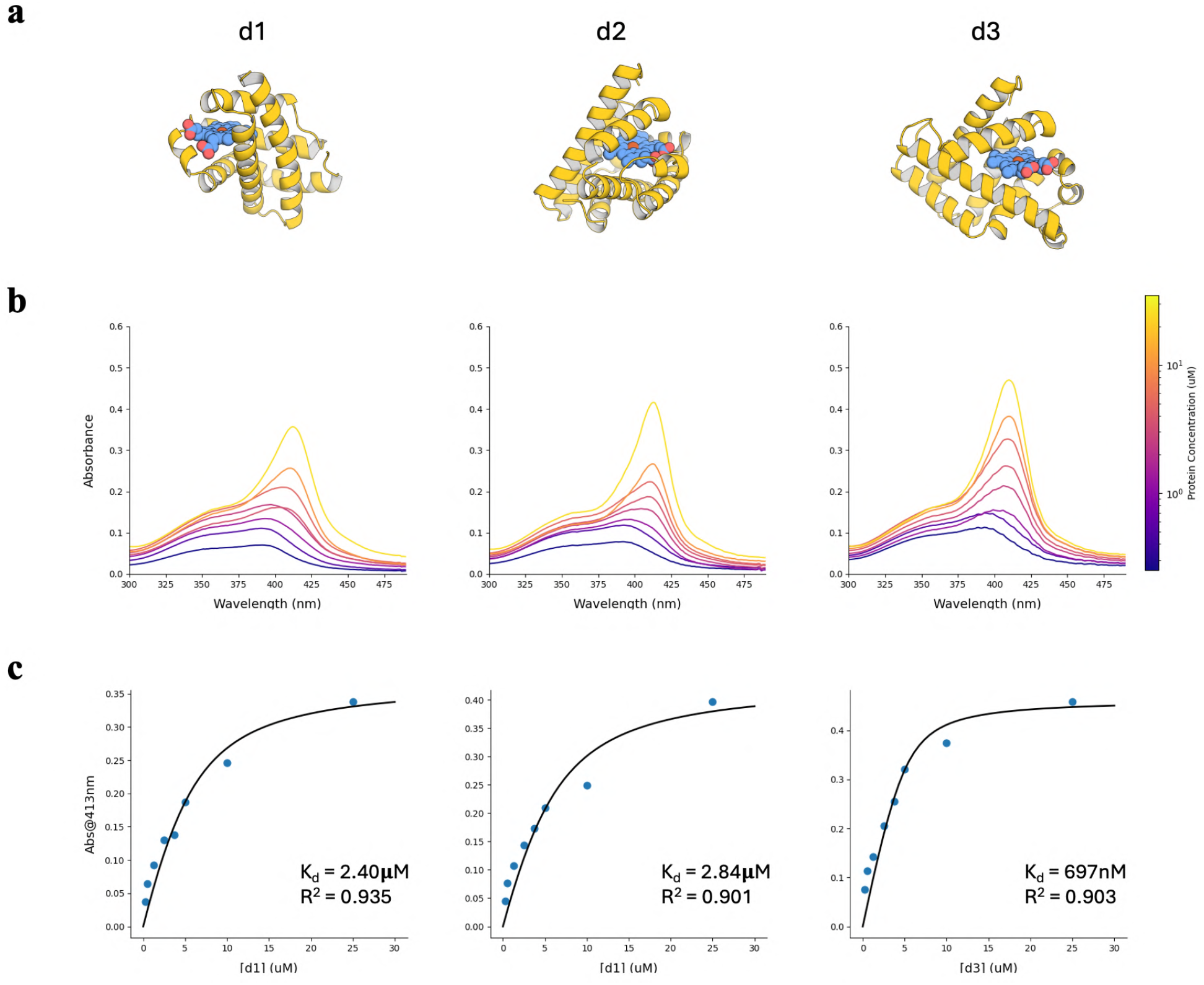
Heme Binder Design. **a)** Structural models of the designed porphyrin binders in complex with ferrous heme. **b)** UV–vis spectra of Fe(II) heme (5 *µ*M) upon titration with increasing concentrations of the designed proteins (0–5 equivalents). The Soret band shifts from 398nm to approximately 413 nm, consistent with histidine-ligated ferrous heme. **c)** Dissociation constants (*K*_*d*_) were determined from the absorbance at 413 nm by fitting to a quadratic ligand-binding model.

**Figure 27:**
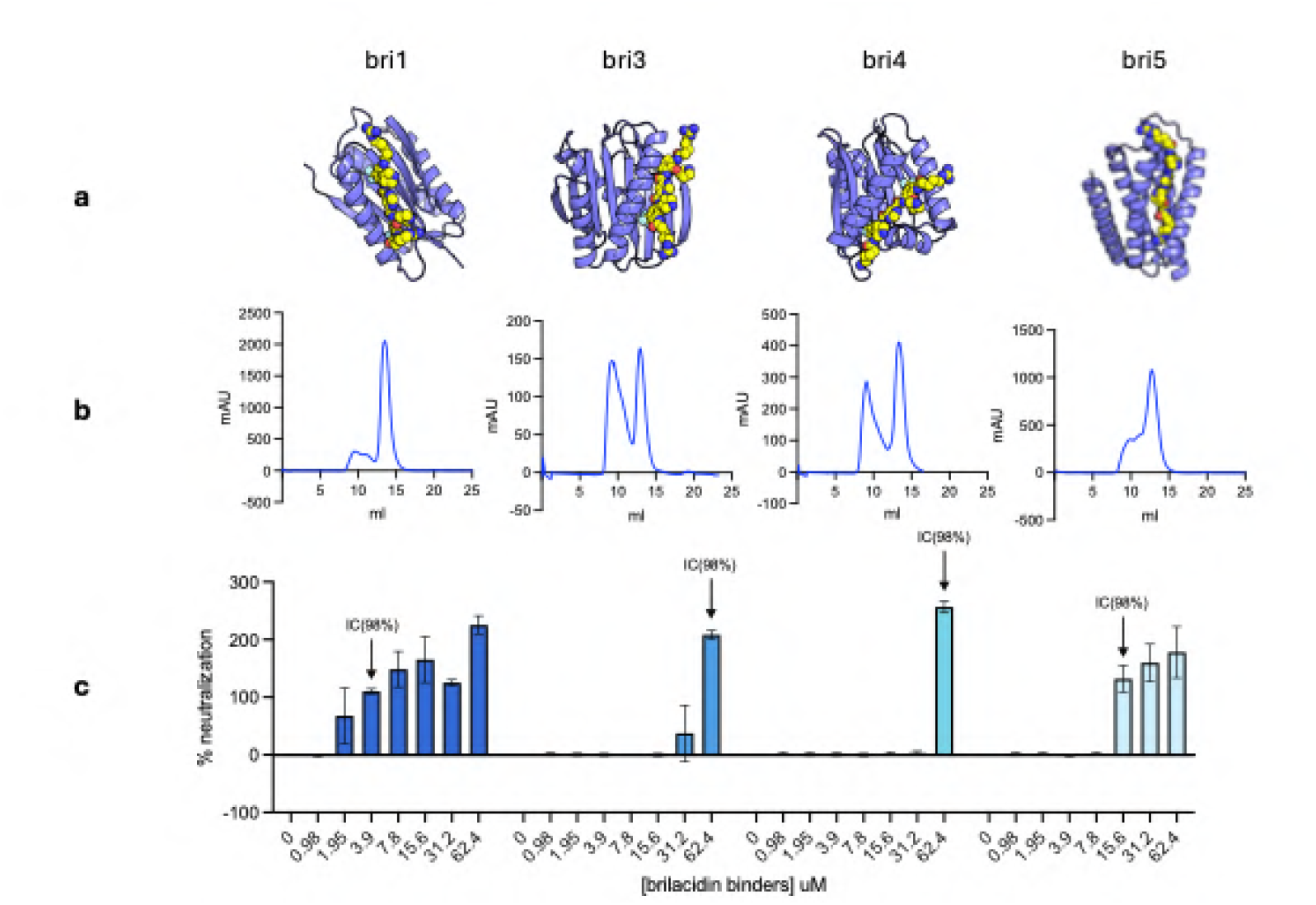
Brilacidin Binder Designs. **a)** Structural models of the designed brilacidin binders (purple) in complex with brilacidin (yellow). **b)** Size exclusion chromatography traces of bri1–bri5 show varied oligomeric states. **c)** Neutralization assays of brilacidin (1.56 *µ*M) upon titration with increasing concentrations of designed protein (0–62.4 *µ*M). Neutralization was assessed as: % neutralization 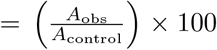, where *A*_obs_ and *A*_control_ are, respectively, the background-subtracted endpoint absorbance at 600 nm for the protein-treated sample versus the bacteria-only control (in the absence of both protein and brilacidin). At high concentrations, the addition of protein increased the turbidity, causing an increase in absorbance relative to control (resulting in a % neutralization value slightly larger than 100%).

**Figure 28:**
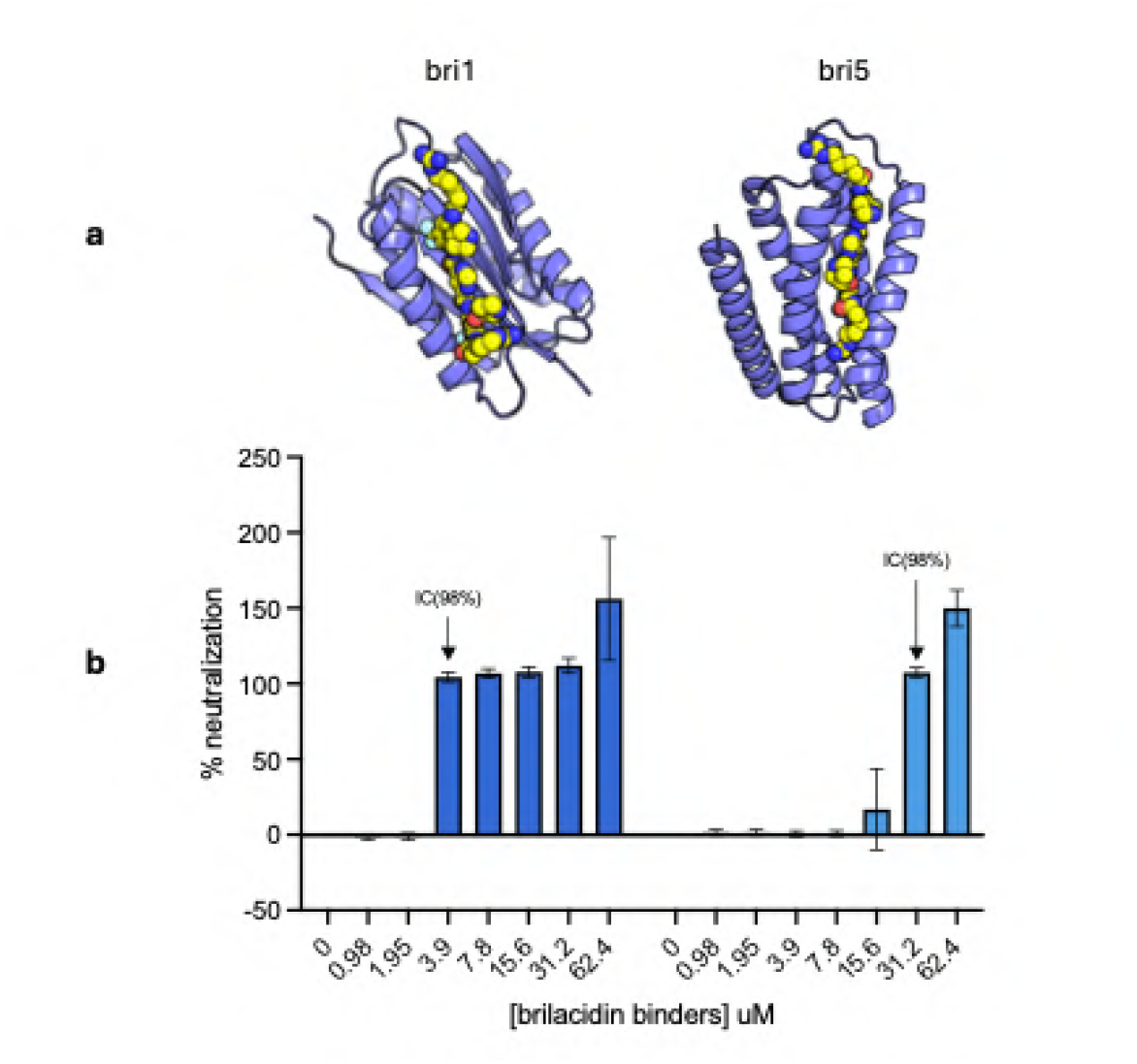
Brilacidin binder design and neutralization assays. **a**, Structural models of the designed brilacidin binders (purple) in complex with brilacidin (yellow). **b**, Neutralization assays of brilacidin (1.56 *µ*M) upon titration with increasing concentrations of designed protein (0–62.4 *µ*M). Neutralization was quantified as % neutralization = (*A*_obs_*/A*_control_) × 100, where *A*_obs_ and *A*_control_ denote the background-subtracted endpoint absorbance at 600 nm for the protein-treated sample and the bacteria-only control (in the absence of both protein and brilacidin), respectively. At high protein concentrations, turbidity increased, resulting in absorbance values above the control and thus % neutralization values slightly greater than 100%.

**Figure 29:**
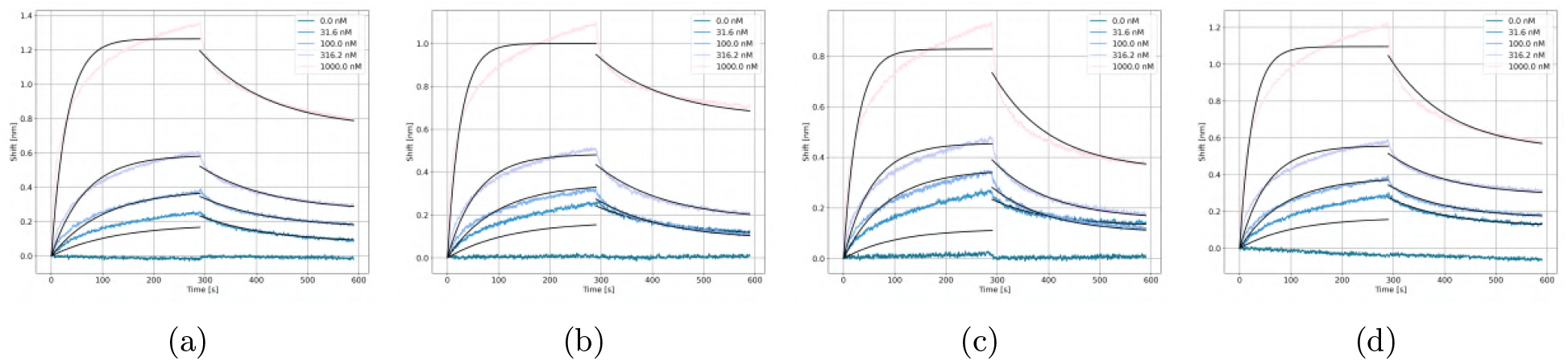
Sensorgrams for the two NPM1 binders.

**Figure 30:**
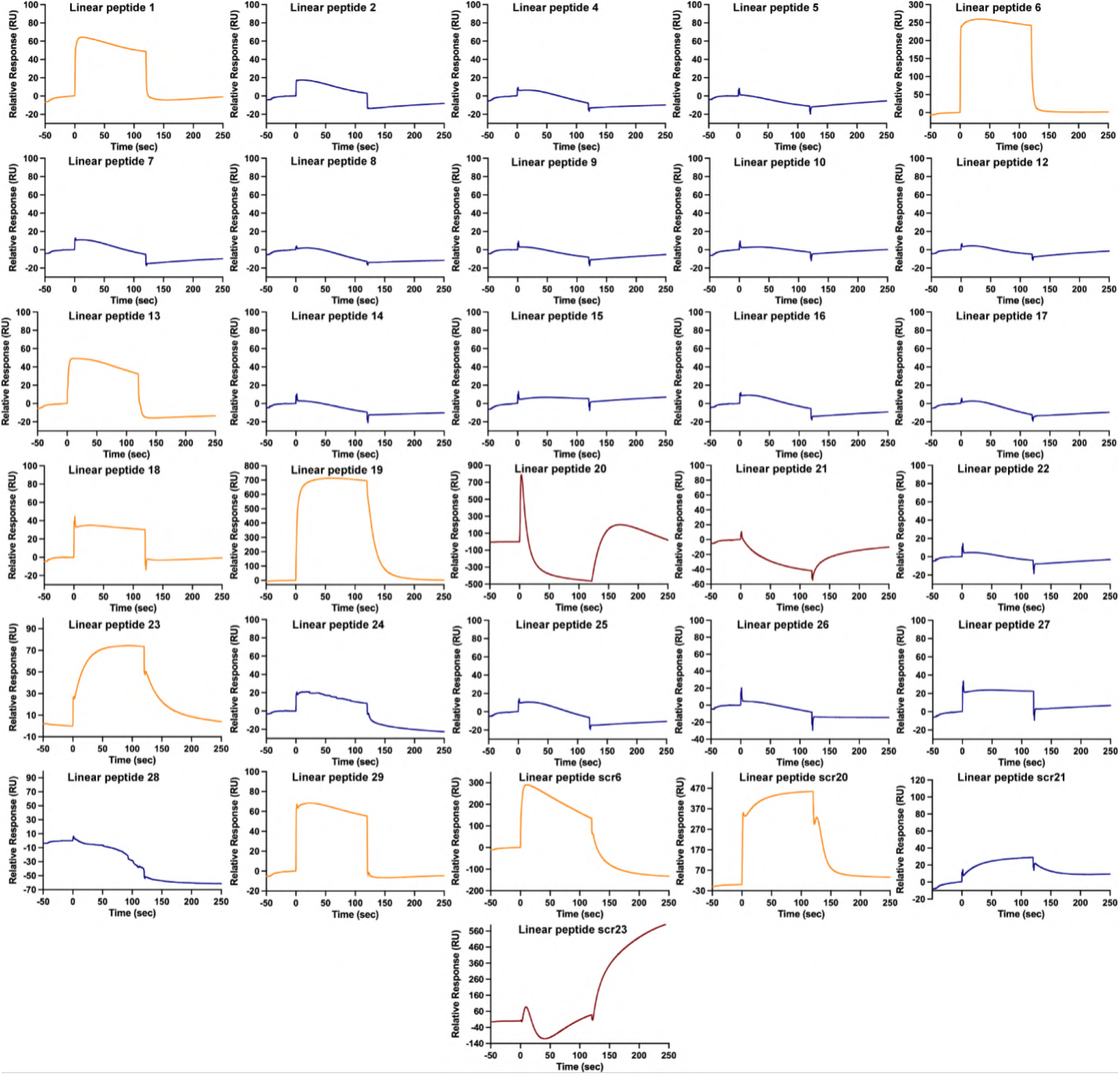
Linear peptide binding response to Rag GTPase. Individual sensorgrams corresponding to each peptide measured at 100*µ*M concentration. The measurements were taken through the same time range, and the plots were generated using GraphPad Prism (Version 10.6.0) with time in seconds on the x-axis and relative response in the y-axis. The curves shown in orange are annotated as binders, the curves in blue are non-binders, and crimson curves represent peptides with suboptimal binding response. The peptides designated as binders and suboptimal binders were further tested at different peptide concentrations to compute binding affinity (*K*_*d*_) as shown in Supplementary Figure 31 and Supplementary Table 14.

**Figure 31:**
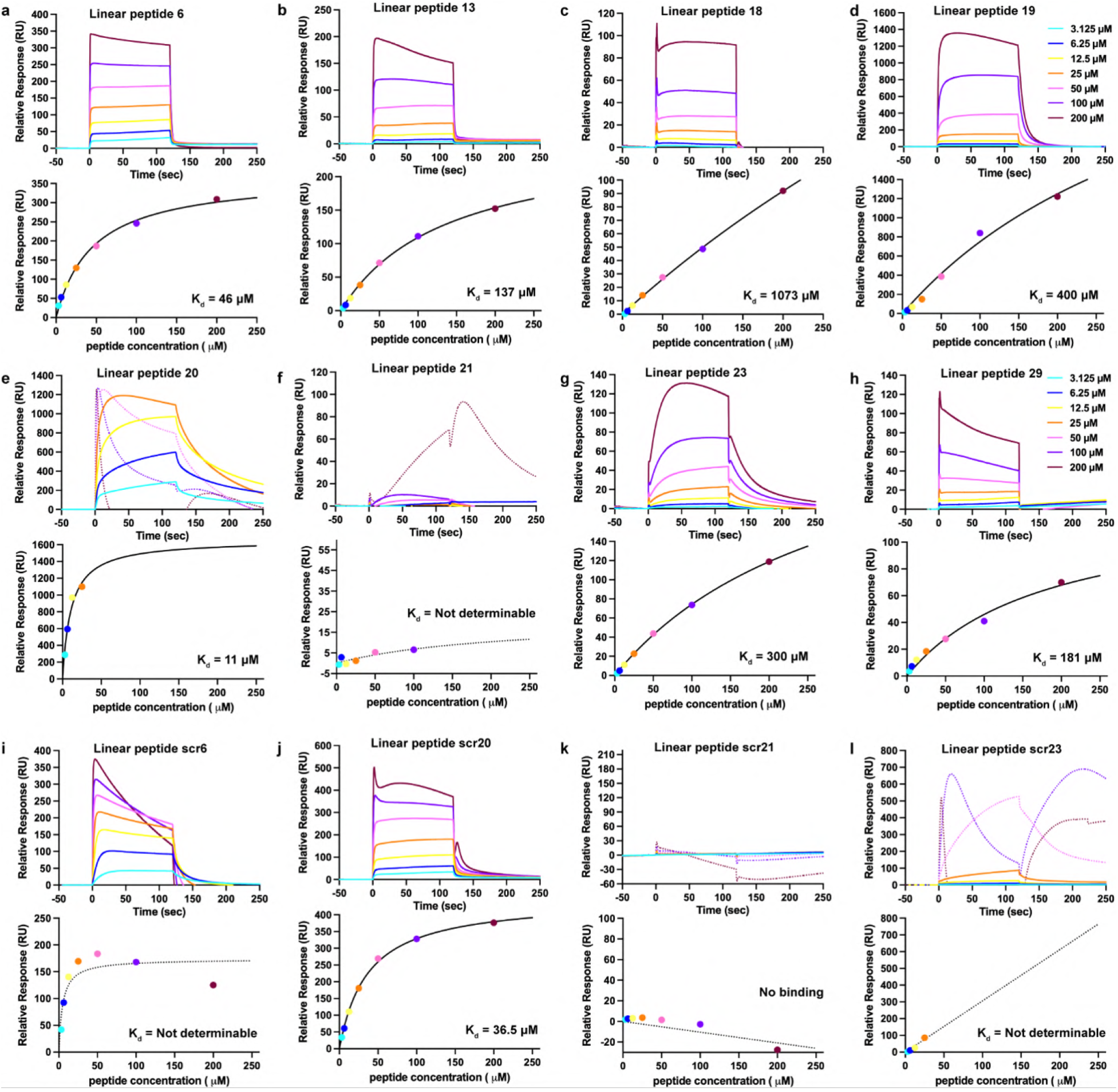
Binding affinity of linear peptides to Rag GTPase. Each panel shows the sensorgram corresponding to individual concentration ranging from 3.25–200 *µ*M for a particular peptide. Using the mean of the values obtained from the last 5 secs of the association step for each peptide as computed by the Biacore™ Insight Software, the binding curve was generated and the *K*_*d*_ (dissociation constant) computed using GraphPad Prism (Version 10.6.0). In all the sensorgrams, the suboptimal responses were plotted with dotted lines. Other than linear peptide 20 that has suboptimal response beyond 25 *µ*M (k), all other binders have detectable binding at all concentrations and have *K*_*d*_ ranging between 11–1073 *µ*M (Table 2). For linear peptide 20, the *K*_*d*_ was computed using values up to 25 *µ*M (g). For the scrambled peptides (f,j) corresponding to some of the binders (e,i), the response is suboptimal and the *K*_*d*_ cannot be determined (f,j). Linear peptide 21 and its scrambled version (k,l) has suboptimal response with non-determinable *K*_*d*_ and no binding respectively.

**Figure 32:**
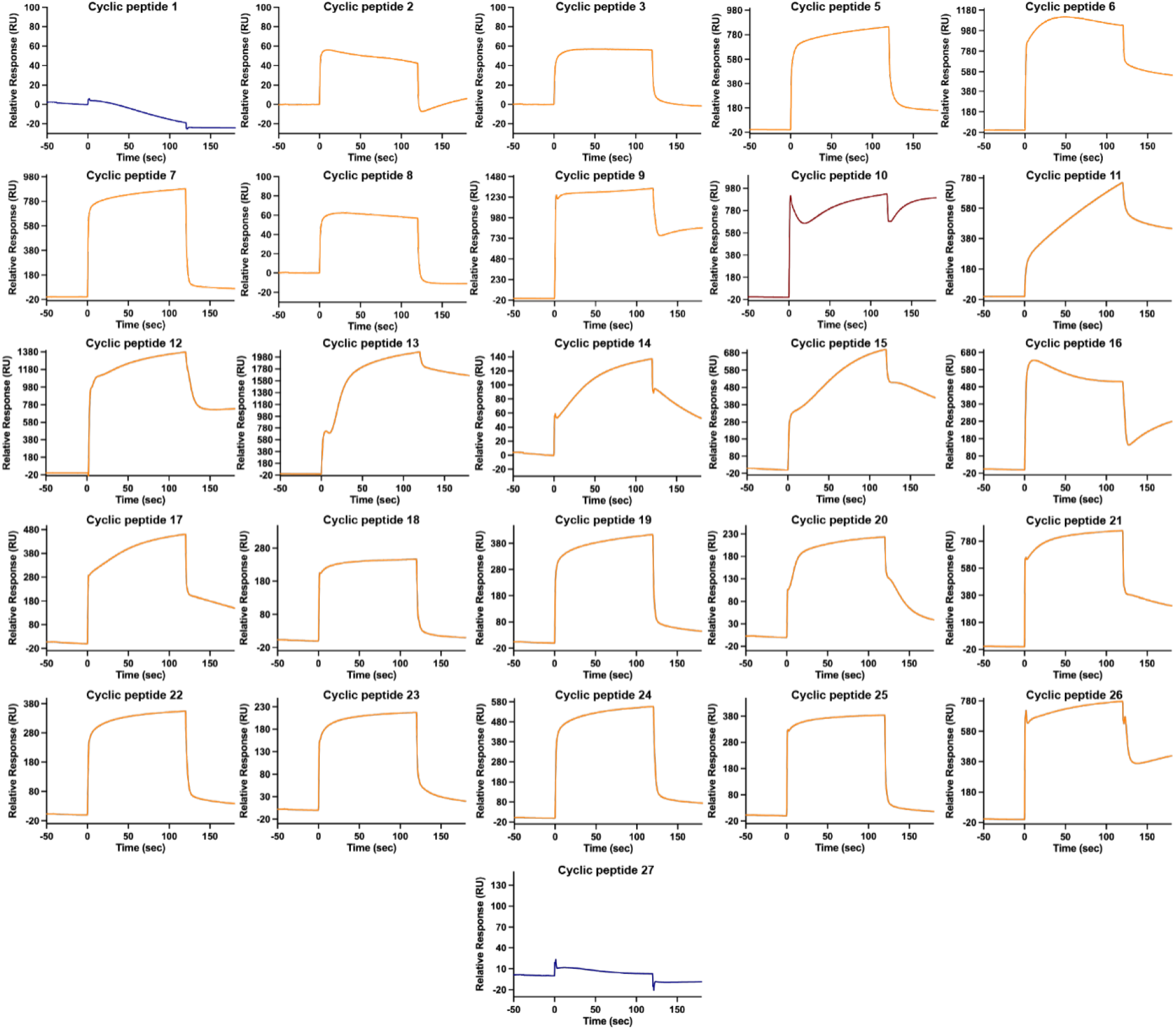
Cyclic disulfide bonded peptide binding response to Rag GTPase. Individual sensorgrams corresponding to each peptide measured at 100 *µ*M concentration. The measurements were taken through the same time range, and the plots were generated using GraphPad Prism (Version 10.6.0) with time in seconds in the x-axis and relative response in the y-axis. The curves shown in orange are annotated as binders, blue are non-binders. The binders were further tested at different peptide concentrations to be able to compute binding affinity (*K*_*d*_) as shown in Supplementary Figure 33 and Supplementary Table 16.

**Figure 33:**
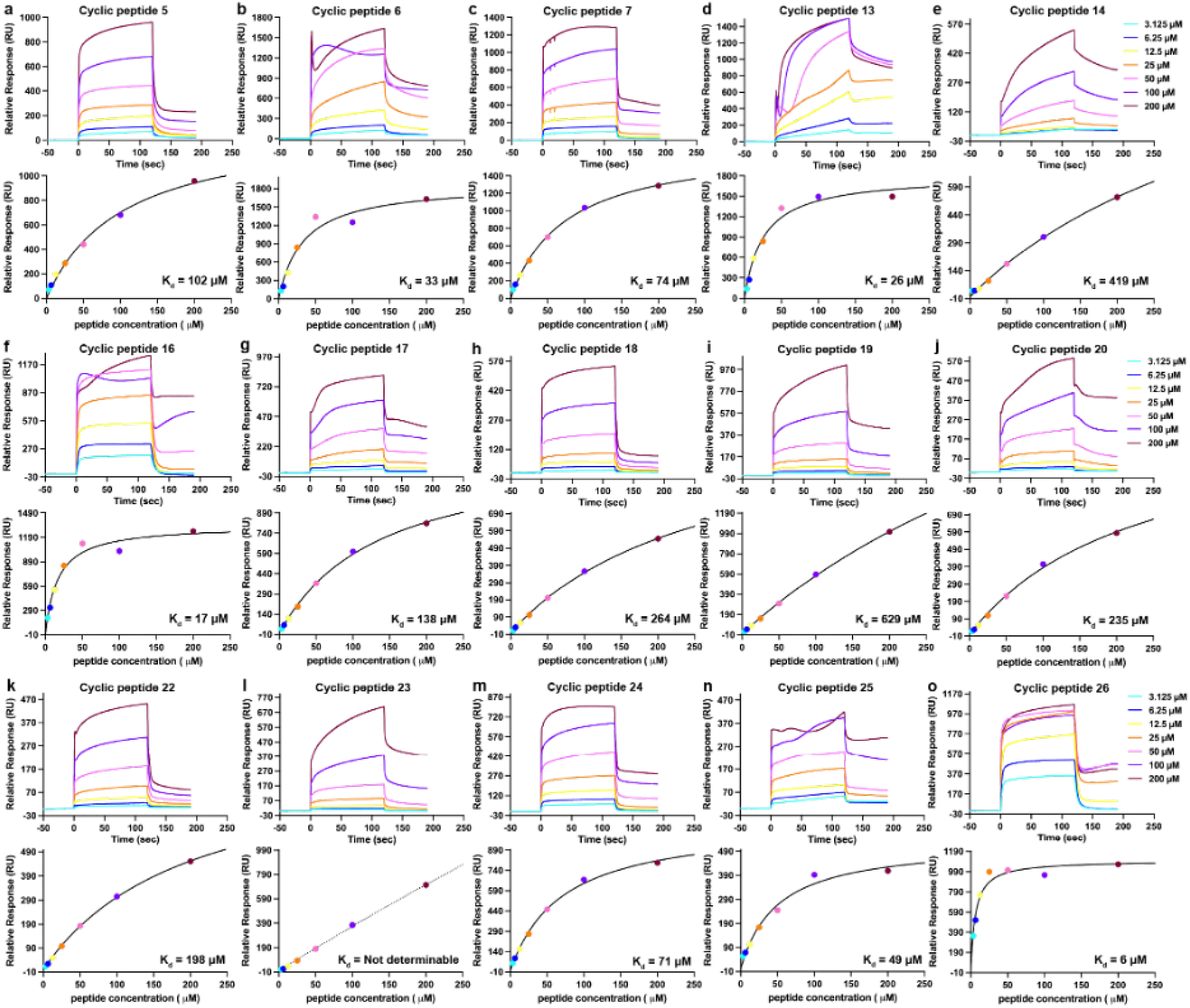
Binding affinity of cyclic disulfide bonded peptides to Rag GTPase. Each panel shows the sensorgram corresponding to individual concentration ranging from 3.25–200 *µ*M for a particular peptide. Using the mean of the values obtained from the last 5 secs for each concentration for each peptide as computed by the Biacore™ Insight Software, the binding curve was further generated and the *K*_*d*_ (dissociation constant) computed using GraphPad Prism (Version 10.6.0). Other than cyclic peptide 23 that has increasing signal beyond 200 *µ*M and couldn’t be fit into a model to obtain affinity (l), all other binders have detectable binding at all concentrations and have *K*_*d*_ ranging between 6–630 *µ*M (Table 16). Cyclic peptides – 6(b), 16(f) and 25(n) exhibit binding and have a determinable *K*_*d*_ under the tested conditions, but show suboptimal response in at least one concentration.

**Figure 34:**
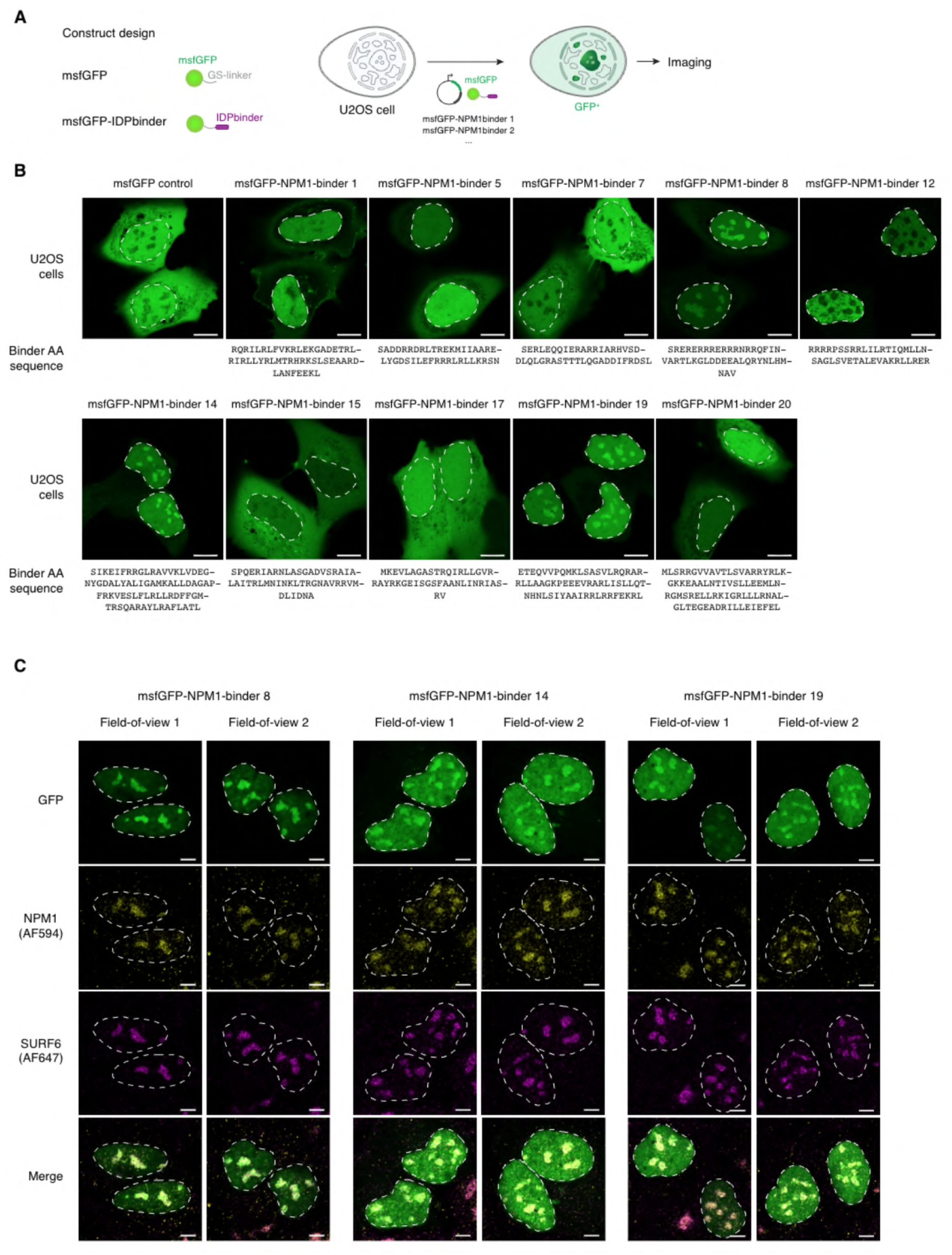
**A.** Schematic model of the msfGFP-tagged NPM1-binder design and cellular assay to visualize subcellular localization of the NPM1-binders. **B**. Representative live cell fluorescence microscopy images of U2OS cells expressing ectopic msfGFP-NPM1-binders. The cell nucleus is highlighted with a dashed white line contour. Scale bar: 10 µm. The experiment was repeated twice independently with similar results. **C**. Representative fixed-cell immunofluorescence of U2OS cells expressing exogenous msfGFP-NPM1-binder 8, 14, 19. Endogenous NPM1 and SURF6 are stained with antibodies. The cell nucleus is highlighted with a dashed white line contour. Scale bar: 5 µm. The experiment was repeated twice independently with similar results.

**Figure 35:**
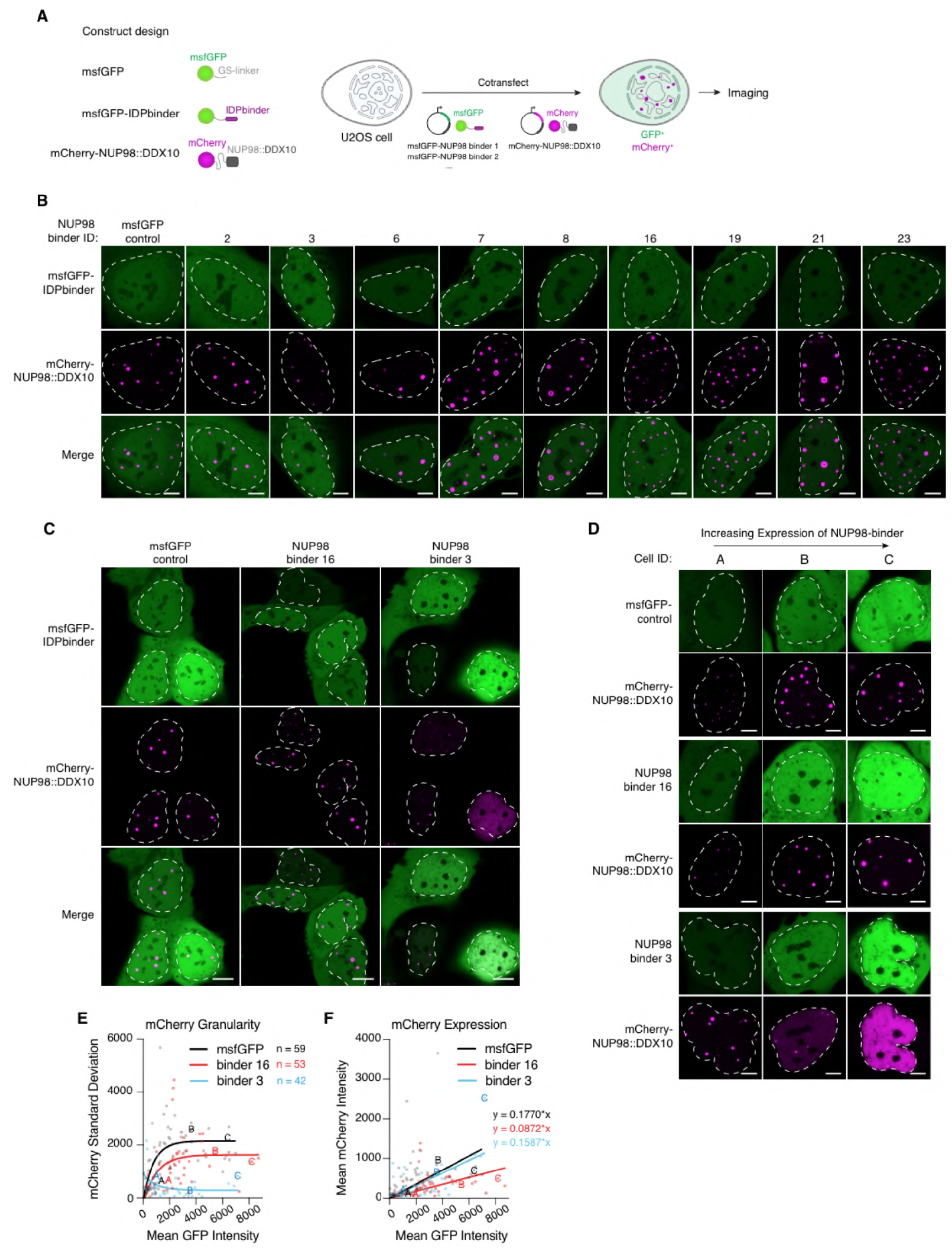
**A.** Schematic of the msfGFP-tagged NUP98-binder constructs and the mCherry-NUP98::DDX10 reporter (containing the NUP98 IDR), and the live-cell co-expression assay used to visualize their subcellular localization. **B**. Representative live-cell fluorescence microscopy images of U2OS cells co-expressing msfGFP-NUP98-binders and mCherry-NUP98::DDX10. The nucleus is outlined with a dashed white contour. Scale bar: 5 µm. The experiment was repeated twice independently with similar results. **C**. Representative widefield fluorescence microscopy images of U2OS cells co-expressing msfGFP control, msfGFP-NUP98-binder-16, or msfGFP-NUP98-binder 3 together with mCherry-NUP98::DDX10. The nucleus is outlined with a dashed white contour. Scale bar: 10 µm. The experiment was repeated twice independently with similar results. **D**. Representative single-cell examples showing increasing msfGFP-NUP98-binder expression (left to right) with corresponding mCherry-NUP98::DDX10 signal distribution. Cell IDs match the labeled data points in E and F. The experiment was repeated twice independently with similar results. **E**. Nuclear mCherry signal standard deviation (granularity) plotted against mean nuclear GFP intensity for msfGFP control, NUP98-binder 16, and NUP98-binder 3. Labeled points correspond to the cells shown in D. n = 59 (msfGFP control), 53 (msfGFP-NUP98-binder 16), and 42 (msfGFP-NUP98-binder 3) cells from two biologically independent experiments. Curves show one-phase decay fits. **F**. Mean nuclear mCherry intensity plotted against mean nuclear GFP intensity for the cells quantified in E. Labeled points correspond to the cells shown in D.

### E.7 Antimicrobial Peptides against Bacterial GyrA

A library of DNA templates encoding designed variants, mutated variants with 3 alanine substitutions at the binding interface, alongside a library encoding protein fragments tiling GyrA and eGFP, was generated (Twist Biosciences), and massively parallel relative growth measurements in E. coli were performed as previously [Savinov et al., 2022, 2025].

Specifically, the library of coding sequences was cloned into the pET-9a expression vector (Novagen) exactly as previously [Savinov et al., 2022, 2025]. The plasmid library encoding designed binders and protein fragments was then transformed into electrocompetent *E. coli* BL21 (DE3) (Sigma-Aldritch) at ≥ 110-fold coverage of the library size, and following 1-hr recovery from transformation, cells were immediately diluted into LB media (Gibco) containing kananycin (selecting for presence of the library) and 10 uM IPTG (inducer for library expression), beginning the massively parallel inhibition measurements. Cells were then grown to an OD_600nm_ of 1.5, at which point they were harvested. These experiments were performed in triplicate (3 biological replicates). Plasmids were extracted from each sample (Qiagen miniprep kit), and DNA from each output sample as well as the plasmid library input was prepared for high-throughput sequencing as previously [Savinov et al., 2022]. Paired-end sequencing was performed on a Singular G4 platform. From these measurements we determined designed peptide and protein fragment frequencies in the population (*f*) at the initial and final growth assay timepoints, allowing calculation of the enrichment *E* = log2(*f*_initial/*f*_final). The inhibition score for each peptide was then calculated as Inhibition = -*E*, such that larger positive values correspond to stronger inhibitory effects. Results across biological replicates of these measurements were highly reproducible as in prior work [Savinov et al., 2022, 2025].

The specificity of designed binders for the designed binding mode to GyrA was calculated as Δ(Inhibition) = Inhibition(designed binder) - Inhibition(mutated binder). Positive Δ(Inhibition) values therefore correspond to binders which are more inhibitory than their corresponding variants where 3 interface residues are mutated to alanine. Designed binders and protein fragments that substantially inhibit cell growth were defined as those with inhibition scores *E*≤ -2, corresponding to a ≥4-fold reduction in relative growth. Results were similar with a less stringent threshold of *E* ≤ -1, and so the more conservative threshold was employed. This threshold picks out previously identified inhibitory peaks from protein fragments tiling across GyrA [Savinov et al., 2022, 2025].

## F Additional Experimental Results

### F.1 Nanobodies against Recently Deposited and Novel Pathogen Targets

We selected two monomeric viral or bacterial targets deposited in the PDB in 2024 that share low similarity with any protein previously observed in a bound context in the PDB. This choice emulates the emergence of a novel pathogen and the subsequent identification of a therapeutically relevant target. Specifically, we selected the cyclic GMP–AMP phosphodiesterase from Penguinpox (cGAMP PDE), which shares 24% sequence identity with the closest protein in the PDB and inhibits host STING signaling by degrading cyclic dinucleotides [Hobbs et al., 2024]. As a second target, we selected Filamentous Hemagglutinin (FhaB), an adhesion protein expressed by *Bordetella* that promotes colonization and infection of host tissues [Costello et al., 2025] and shares 30% sequence identity with the closest protein in the PDB. We generated 60,000 nanobody designs per target and experimentally evaluated the seven highest-scoring candidates by displaying the antibodies on cells using yeast surface display (YSD) and testing what antigen concentrations resulted in detectable labeling of yeast populations.

All designs displayed well, suggesting proper expression and folding. However, binding was not strongly detected. Across antigen concentrations from 7 nM to ∼2 µM, we observed that one of seven yeast-displayed nanobody designs against cGAMP PDE showed detectable but weak binding only at 1.6 µM antigen. We observed that all seven yeast-displayed nanobody designs against FhaB—targeting a single epitope—showed detectable but weak binding only at 2 µM antigen (Figure 6). Because 1.6 µM or 2 µM was the highest antigen concentration used for labeling and was the minimal concentration that resulted in any binding signal, we cannot calculate an EC50 from these studies. Instead, we can conclude that the binders have much higher *K*_*d*_ than low µM, given that low µM concentrations of antigen were required to even begin to observe a labeling signal. Nevertheless, YSD is often used to experimentally optimize computational antibody designs, and the low binding signal observed should be sufficient to initiate YSD-based antibody affinity optimization campaigns.

### F.2 Sensorgrams from Adaptyv and Sino Biological

We provide all sensorgrams from Adaptyv Bio and Sino Biological for download here for miniprotein designs (https://huggingface.co/datasets/boltzgen/update_sensorgrams/resolve/main/protein_sensorgrams.xlsx?download=true) and nanobody designs (https://huggingface.co/datasets/boltzgen/update_sensorgrams/resolve/main/nanobody_sensorgrams.xlsx?download=true).

### F.3 Additional Tables and Figures

## Footnotes

1 For instance, the target could stick to the surface itself rather than the design that is attached to the plate (which is an issue that cannot be corrected for via a reference channel if the design attachment process altered the plate itself).

